# Conformational dynamics of exopolysaccharides underlie biofilm matrix mechanics in *Vibrio cholerae*

**DOI:** 10.64898/2026.07.22.739955

**Authors:** Kee-Myoung Nam, Nathan Fowler, Rajan Kandel, Yongqi Zhu, Yuchu Liu, Yi-Jhen Lai, Muhammad Faheem Hassan, Emma Gerace, Merrill Asp, Rich Olson, Ying Li, Mu-Ping Nieh, Mingjiang Zhong, Robert J. Woods, Alexis Moreau, Jing Yan

## Abstract

Polysaccharides remain the least understood biomacromolecules, particularly in terms of the relationship between their chemical structure and physical properties. On the other hand, polysaccharides often serve as the main structural components in biofilms: surface-attached aggregates of bacterial cells encased within a mechanically resilient extracellular matrix. The large chemical space explored by bacteria within biofilms provides excellent opportunities to establish the structure-function relationship for polysaccharides. In this paper, we systematically characterize various polymer properties of <u>V</u>ibrio <u>p</u>oly<u>s</u>accharide (VPS), the major exopolysaccharide in biofilms formed by *Vibrio cholerae*, the causative agent of pandemic cholera. Using a combination of shear rheology, dynamic and static light scattering, and small-angle X-ray scattering, we measure the viscosity, molecular weight, persistence length, radius of gyration, and hydrodynamic radius of this chemically unique biopolymer. Combining all-atom and coarse-grained simulations, we show how the conformational flexibility of a single glycosidic linkage within each VPS monomer can lead to dramatic compaction of the entire polymer chain and nonclassical entanglement behavior. Our comprehensive quantification represents a rare endeavor for bacterial biofilms, whose matrix composition and physical properties remain largely nebulous; it also represents a significant step towards a detailed understanding of the molecular origins of biofilm mechanics.

## INTRODUCTION

Biopolymers exhibit many interesting physical properties rarely found in conventional materials^1^. Compared to proteins and polynucleotides, polysaccharides remain the least understood among the major biopolymers important for life. The complex stereochemistry of monosaccharides, including that of the anomeric carbon (α versus β), as well as the variety of possible glycosidic linkage positions between monosaccharides, render the synthesis and characterization of polysaccharides much more challenging than for proteins and nucleic acids^2^. In bacteria, this endeavor is further complicated by the fact that bacteria can produce over 700 different kinds of monosaccharides, some with rare chemical structures, whereas mammalian cells rely on only 10 primary monosaccharides for function^3,4^. Over the past two decades, the study of microbial glycans has evolved into a burgeoning field that has generated many new insights into glycobiology unique to bacteria, as well as new tools for their study^5,6^.

Even less well understood is how the chemical structure of bacterial polysaccharides gives rise to their physical properties, including their conformational dynamics and solution properties. These properties are especially important in biofilms: surface-attached bacterial communities embedded within a self-produced extracellular matrix, which are often composed predominantly of exopolysaccharides^7–9^. The matrix serves as a scaffold that determines the biofilm’s architecture, mechanical stability, and resistance to environmental stresses^10–12^, and plays a key role in facilitating antibiotic tolerance in biofilm-associated chronic infections^13^ and biofouling of industrial surfaces^14^. The matrix is often described as a heterogeneous hydrogel composed of various polymers, including polysaccharides, proteins, and sometimes extracellular DNA. Among these polymers, exopolysaccharides often play a dominant role in determining the viscoelasticity, cohesion, and overall functionality of the matrix. Consequently, the mechanical properties of biofilms are closely linked to the physicochemical properties of their exopolysaccharides, including their molecular weight, conformational dynamics, local concentration, and interactions with other matrix components^10–12^. Despite this, our understanding of these properties and how they dictate biofilm mechanics remains limited, in large part due to the difficulty of purifying exopolysaccharides from biofilms and experimentally characterizing their properties.

Recent progress in understanding the biochemistry of the biofilm matrix in *Vibrio cholerae*, the causative agent of pandemic cholera^15^, offers a unique opportunity to investigate the structure-function relationship of exopolysaccharides (Fig. 1A). The formation and structural integrity of *V. cholerae* biofilms critically depend on a single secreted polysaccharide, <u>V</u>ibrio <u>p</u>oly<u>s</u>accharide (VPS)^16,17^. VPS has a chemically unique structure, consisting of a repeating tetrasaccharide unit containing a modified L-gulose residue (N-acetylated at C2, O- acetylated at C3, and bearing an amide-linked glycine at C6), followed by two glucose residues and one galactose residue^16^. All glycosidic linkages are α-(1 → 4), except for a β-(1 → 4) linkage connecting the two glucose residues. This unusual composition and linkage pattern determine how VPS interacts with other matrix components^18–20^, including the cell-cell adhesion protein RbmA^21,22^ and cell-surface adhesion proteins Bap1 and RbmC^23–25^.

**Fig. 1.**
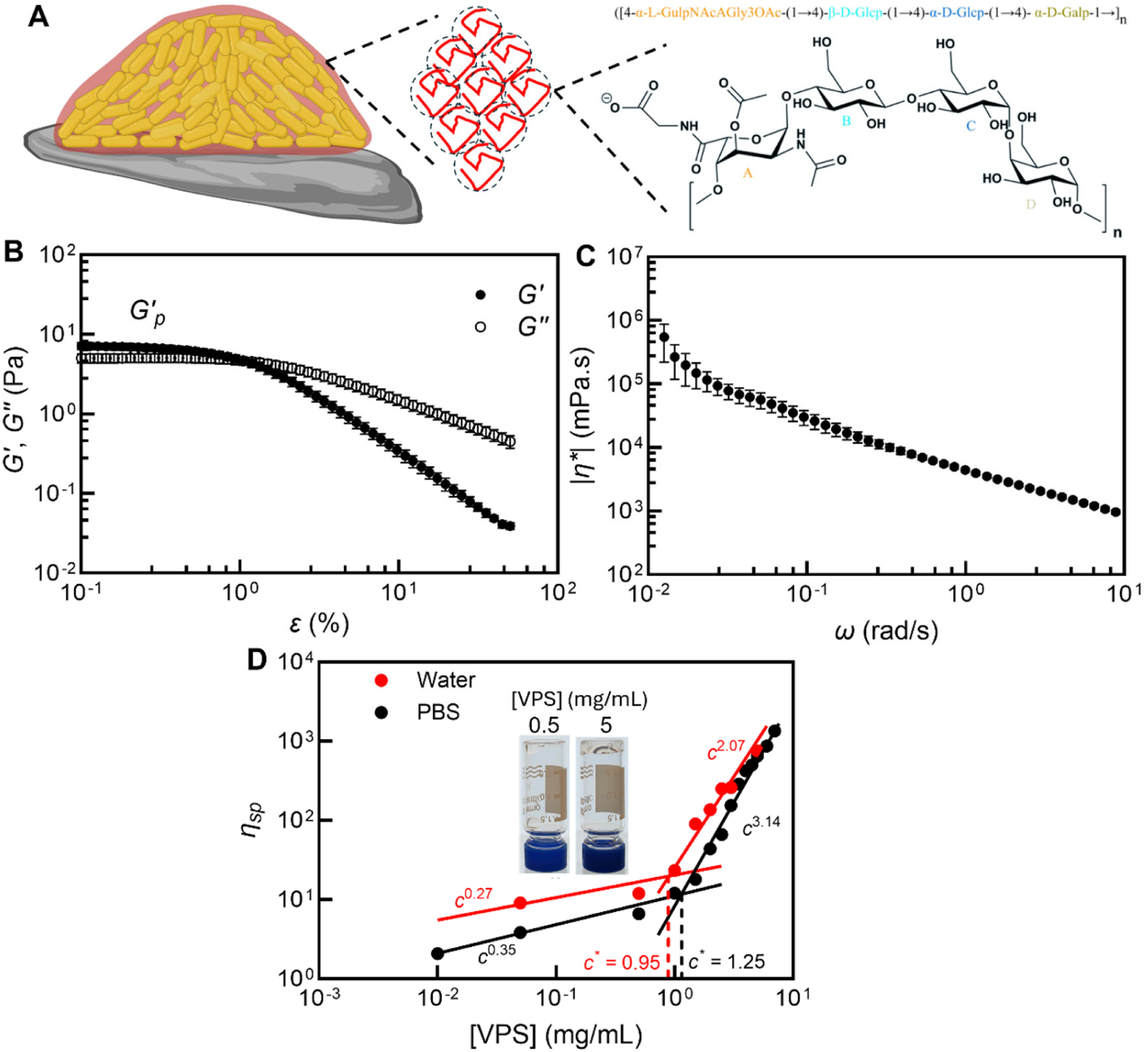
Rheological characterization of VPS, the main matrix component of *Vibrio cholerae* biofilms. (**A**) Schematic of a *V. cholerae* biofilm and its main matrix component <u>V</u>ibrio <u>p</u>oly<u>s</u>accharide (VPS). Shown on the right is the chemical structure of the repeating unit in VPS. Created with BioRender.com. (**B**) Representative storage modulus (*G′*) and loss modulus (*G′′*) curves as functions of the amplitude of oscillatory shear strain *ε*, measured for a VPS solution in phosphate-buffered saline (PBS) buffer at 7 mg/mL. The plateau modulus, *G′*_p_, is the value of *G′* in the plateau region. (**C**) Frequency sweep of complex viscosity, |*η*\*|, of a VPS solution in PBS at 5 mg/mL. Error bars correspond to standard deviations (*n* = 3). (**D**) Specific viscosity, *η*_sp_, as a function of VPS concentration in PBS (black) or in water (red). Each datapoint corresponds to the average of at least three independent replicates. Inset shows pictures of VPS solutions in flipped vials at two concentrations, one below (0.5 mg/mL) and one above (5 mg/mL) the crossover concentration, *c*\*.

Despite their importance, the physicochemical properties of VPS—including its molecular weight, viscosity, persistence length, and radius of gyration—have not yet been characterized. Determining these properties is essential for developing a deeper understanding of how VPS is organized within the biofilm and—in tandem with other matrix components— contributes to the biofilm’s mechanical resilience and viscoelasticity. In this study, we use a combination of biophysical tools, imaging, and multiscale simulations to characterize the physicochemical and polymeric properties of VPS, both *in vitro* and *in vivo*. Our findings suggest a model in which spontaneous rotation of one glycosidic linkage within each VPS monomer introduces discrete, sharp kinks along an otherwise stiff polymer, thus providing avenues to chain compaction and entanglement that are distinct from those available to classical flexible or semiflexible polymers.

## RESULTS

### Rheological characterization of VPS

We began by investigating the rheological properties of VPS. We purified VPS from *V. cholerae* cultures following a previously described protocol, using a strain that constitutively makes VPS but lacks all of the accessory matrix proteins^16,26–28^, yielding approximately 15–20 mg of solid polysaccharide per batch of purification (Fig. S1A). The purified polymer could be readily dissolved up to a concentration of ∼10 mg/mL in water or phosphate-buffered saline (PBS) buffer; PBS was chosen as a convenient salty buffer commonly used in microbiology laboratories, as it is compatible with cell viability and maintains pH and osmotic balance. This homogeneous solution was then transferred to a shear rheometer for mechanical characterization (Fig. S1A). Oscillatory shear experiments were then performed to determine the viscoelastic properties of VPS solutions (Figs. 1B, S1B, and S1C). By applying oscillatory shear strains (*ε*) and measuring the resulting stresses (*σ*), we extracted the storage modulus *G′*, which reflects the elastic, solid-like behavior of the polymer solution; and the loss modulus *G′′*, which reflects the viscous, fluid-like response. We found that VPS solutions are viscoelastic materials akin to hydrogels, with *G′* > *G′′* at low *ε*. The *G′*(*ε*) curve generally exhibits an initial plateau, where VPS deforms elastically with increasing strain, defining a plateau modulus *G′*_p_. Beyond a critical yield strain *ε*_Y_, *G′* sharply decreases, indicating network yielding.

To investigate the relationship between the concentration of VPS and its viscoelastic properties, we quantified the plateau storage modulus *G′*_p_, yield strain *ε*Y, and corresponding yield stress *σ*Y for VPS solutions of different concentrations (Fig. S1D). Both *G′*_p_ and *σ*_Y_ increase with VPS concentration, with a significant uptick around 2–3 mg/mL, indicating the emergence of a polymer network.

To further examine the viscoelastic properties of the polymer network, we also performed frequency-sweep experiments. We observed that the magnitude of the complex viscosity, |*η*^∗^|, decreases monotonically with the angular frequency, *ω* (Fig. 1C), consistent with shear-thinning of a viscoelastic polymeric system. Mechanistically, upon the application of shear, the VPS chains align with the shear direction, resulting in decreased flow resistance and consequently a reduction in viscosity^29^. By first extrapolating to the zero-shear viscosity *η*_0_, we calculated the specific viscosity *η*_sp_ of the polymer, defined as

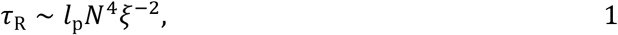

where *η*_s_ is the solvent viscosity. Plotting *η*_sp_ as a function of VPS concentration on a double- logarithmic scale revealed two distinct scaling regimes (Fig. 1D), with a marked crossover at approximately 0.95 mg/mL in water and 1.25 mg/mL in PBS. As a vivid demonstration of the increased viscosity, we show in the inset of Fig. 1D pictures of two VPS solution samples after flipping the vials containing them (see also Fig. S2); beyond the transition point, the VPS solution is sufficiently viscous such that it does not fall by its own weight on the experimental timescale (1 min)^30,31^. Given the presence of a negatively charged carboxylate group on the glycine side chain in the VPS tetrasaccharide unit at neutral pH, the generally higher viscosity in water compared to PBS may be explained by reduced chain expansion in PBS due to ionic screening^32^.

One way to interpret the observed crossover in the scaling of *η*_sp_ is as the overlap concentration, *c*\* (refs. ^29,33^): at concentrations below *c*\*, chains behave largely as isolated coils, whereas at concentrations above *c*\*, chains begin to interact and the solution’s rheological and structural properties are increasingly governed by polymer-polymer contacts. Alternatively, the crossover could be interpreted as the entanglement concentration, *c*_e_, above which the chains create geometrical constraints to restrict each other’s motion^33^. For complex biopolymers, distinguishing between these scenarios experimentally is often not straightforward^32,34^. A third possibility is that the polymer chains engage in attractive interactions, which, beyond a critical association concentration *c*_crit_, give rise to an associative network with increased viscosity and

*G′*_p_ (ref. ^34^). Indeed, potential hydrogen bonding between hydroxyl groups along the polysaccharide backbone and between the glycine side chains, as well as hydrophobic interactions between the N- and O-acetyl groups and the backbone carbon rings, constitute multiple possible modes for polymer-polymer association.

According to various scaling theories, the specific viscosity of a semidilute solution of flexible polymers in a good solvent (with a Flory exponent of v ≈ 0.588) should follow a power law with respect to the concentration *c*, with an exponent that depends on the presence of entanglement and/or associative interactions: without entanglement, *η*_sp_ ∼ *c*^1.3^ for non- associative polymers^33^ and *η*_sp_ ∼ *c*^1.15^ for associative polymers^35^; with entanglement, *η*_sp_ ∼

*c*^3.9–4.5^ for non-associative polymers^33,36^ and *η*_sp_ ∼ *c*^4.75^ for associative polymers^37^. On the other hand, analogous scaling laws for semiflexible polymers have not been as well-established, largely due to debate surrounding the molecular processes underlying the terminal relaxation timescale in the entangled regime^38^; a wide range of exponents have been observed across various settings^39^. Given these caveats, we note that combining the classical predictions of Odijk–Semenov theory with a recently reported scaling relation for the terminal relaxation time in entangled semiflexible chains^38,40,41^ yields a prediction of *η*_sp_ ∼ *c*^2.4^ (see Discussion), which lies in between our measured exponents in water (2.07) and in PBS (3.14). The scaling exponents we observe are also similar to those observed for the polysaccharides xanthan gum^29^ and hydroxypropyl cellulose^39^, as well as the *Staphylococcus epidermidis* exopolysaccharide PIA in the absence of associative interactions^42^. Regardless of the precise underlying mechanism, our rheological data suggest the formation of a network structure with overlapping VPS polymers above this crossover concentration, resulting in a sharply increasing viscosity that harbors critical implications for biofilm mechanics.

### Quantification of VPS concentration within *V. cholerae* biofilms *in situ*

One immediate question following the above analysis is: what is the concentration of VPS within a *V. cholerae* biofilm, and how does it compare with the crossover concentration? To measure this concentration, we adopted an approach described by Ganesan *et al.*^34^, originally developed for *S. epidermidis* biofilms. First, we grew biofilms from a mutant strain (Δ*rbmA*Δ*bap1*Δ*rbmC*) that produces VPS but lacks all the matrix proteins that interact with VPS, and imaged them at single-cell resolution^43^ using confocal microscopy (Fig. 2A). From the confocal *z*-stacks, we found that the volume occupied by each cell in a biofilm, *V*_VPS_, is ∼ 30 µm^3^, which is much larger than the physical size of the bacterium, *V*_cell_, which is on the order of 1 µm^3^ (ref. ^44^). Assuming that the remaining space is permeated with VPS, we sought to quantify the number of VPS molecules occupying this volume.

**Fig. 2.**
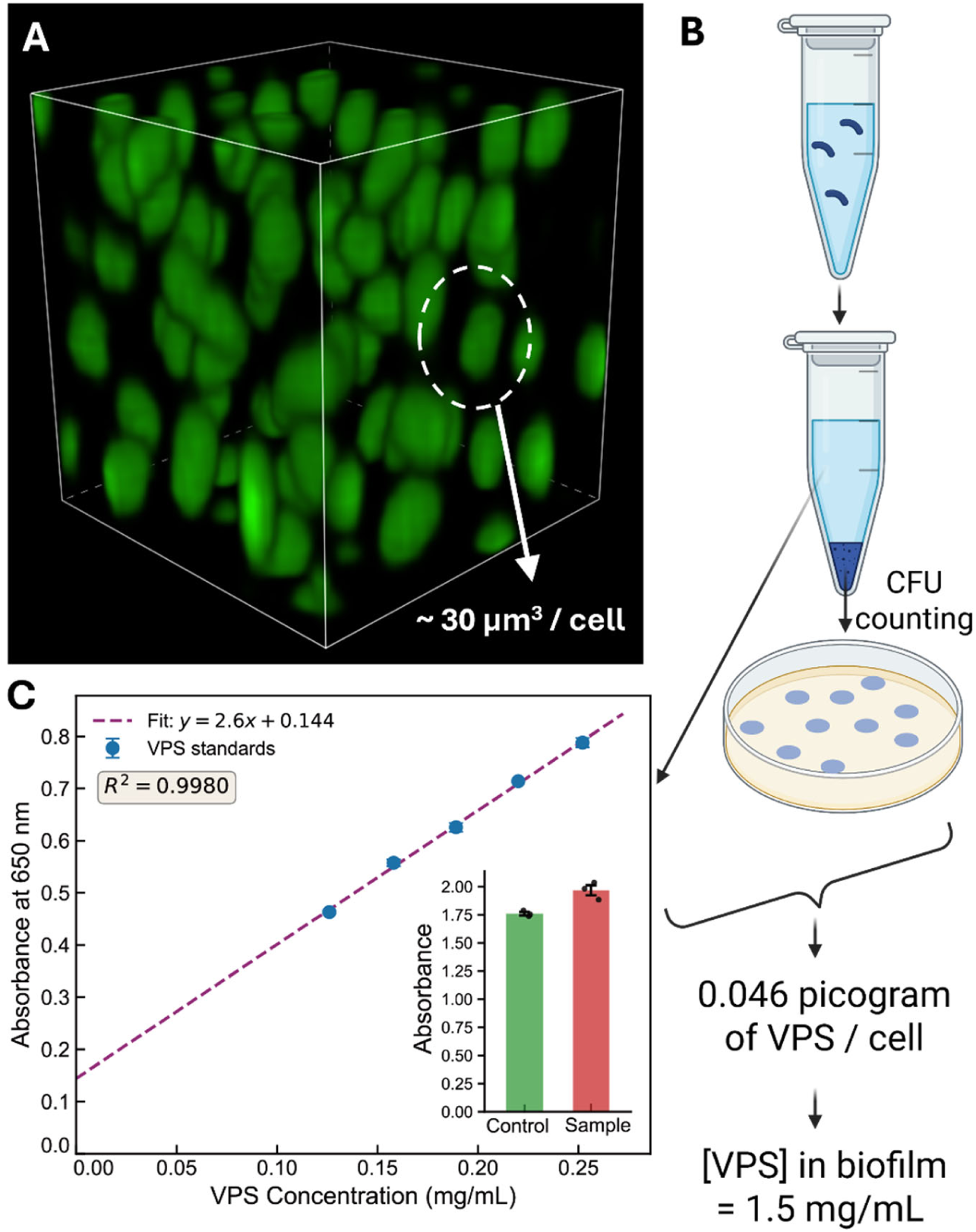
Quantification of VPS concentration within biofilms. (**A**) Three-dimensional rendering of a small region (10 µm × 10 µm × 10 µm) within a biofilm formed by a *V. cholerae* mutant strain in which all matrix proteins are deleted (Δ*rbmA*Δ*bap1*Δ*rbmC*). Cells constitutively express mNeonGreen (green). (**B**) Schematic for the procedure for quantifying VPS concentration within *V. cholerae* biofilms. Briefly, a Δ*rbmA*Δ*bap1*Δ*rbmC* mutant was grown in liquid culture to OD_600_ = 1 (corresponding to 1.75 × 10^9^ colony forming units (CFU) per mL). The culture was then centrifuged, and the supernatant was subjected to an amino- sugar-specific assay for concentration quantification, whereas the cell pellet was resuspended for CFU counting. Created with BioRender.com. (**C**) Results from the Smith–Gilkerson colorimetric assay for VPS quantification in solution. Blue dots correspond to purified VPS with known concentrations, which were used to generate a calibration curve. Green corresponds to results from a negative control strain (Δ*rbmA*Δ*bap1*Δ*rbmC*Δ*vpsL*) that does not produce VPS, and red corresponds to the VPS-producing sample; the difference between the two gives the VPS concentration in the sample.

To do this, we grew the Δ*rbmA*Δ*bap1*Δ*rbmC* mutant in static liquid culture to a known OD_600_ (and hence a known number of cells^26^), and we used the Smith–Gilkerson colorimetric assay for quantifying amino sugars^45^ (Fig. 2B). This assay relies on the N-acetyl group on the unique gulose moiety, as well as the 20% of glucose units (unit C in Fig. 1A) that are substituted with N-acetylglucosamine (GlcNAc). We first obtained a calibration curve using purified VPS; we then quantified the contributions of amino sugars other than VPS to this assay, by performing a control experiment with a non-VPS-producing mutant strain (Δ*rbmA*Δ*bap1*Δ*rbmC*Δ*vpsL*; Fig. 2C). Subtracting these extraneous contributions, the concentration of VPS in the Δ*rbmA*Δ*bap1*Δ*rbmC* supernatant was determined to be 0.081 ±

0.030 mg/mL. Upon converting the measured OD_600_ to colony forming units, we determined the total number of cells (1.75 × 10^9^) corresponding to 1 mL of the culture^26^. Dividing the total amount of VPS by the number of cells, we arrived at 0.046 picograms (pg) of VPS per cell. Note that the dry mass of a rod-shaped bacterial cell is about 0.4–0.5 pg (refs. ^44,46^), which means that the VPS produced by this constitutively VPS-producing strain accounts for a significant portion (about 10%) of the biofilm’s total dry biomass. This is consistent with the significantly decreased growth rate of constitutively VPS-producing *V. cholerae* cells, compared to those that do not produce VPS^47^.

Finally, by dividing 0.046 pg of VPS per cell by the volume *V*_VPS_ per cell (30 µm^3^), we arrived at a VPS concentration of about 1.5 mg/mL in the extracellular space of a *V. cholerae* biofilm (in the absence of any matrix proteins). Intriguingly, this calculation places the extracellular VPS concentration within the biofilm near or slightly above the crossover concentration in Fig. 1D (1.25 mg/mL). This implies that, even in the absence of any matrix proteins that interact with VPS, biofilm-dwelling *V. cholerae* cells produce and retain sufficient VPS to generate a network in their vicinity that imparts some mechanical strength and viscosity to the biofilm. This is consistent with prior measurements of biofilm mechanics^48,49^, and also explains why a Δ*rbmA*Δ*bap1*Δ*rbmC* mutant biofilm, despite having a loose structure, does not completely fall apart under static growth conditions^50^. We note that our calculation should be only considered as an estimate due to the various assumptions outlined above, in addition to other simplifications. In particular, we note that physiological conditions in a liquid culture (used to estimate the mass of VPS surrounding each cell) may differ from those in a biofilm culture (used to estimate *V*_VPS_).

### Hydrodynamic radius of VPS and concentration dependence of the solution structure, as revealed by DLS

To quantify the key molecular properties of VPS underlying its rheology, we performed a series of characterizations at the molecular level. We first performed dynamic light scattering (DLS) measurements of VPS solutions at various concentrations. Fig. 3A shows the resulting autocorrelation functions, *g*_2_(r): curves at low concentrations largely overlap, indicating similar half-lives r_1/2_, while r_1/2_ significantly increases at higher concentrations, signifying large, coherently moving structures. This upturn occurs around 1.5 mg/mL (Fig. 3B), which coincides roughly with the within-biofilm VPS concentration (Fig. 2) and the crossover concentration in the *η*_sp_ vs. *c* curve (Fig. 1D). This is consistent with our interpretation of the rheological transition. We converted the autocorrelation functions to an estimate of the hydrodynamic radius, *R*_h_, using datapoints obtained far below the crossover concentration, where the DLS signal should be dominated by single-chain diffusion; this yielded an average value of ∼ 130 nm for *R*_h_. Near or above the crossover concentration, the DLS signal could reflect collective motion or transient aggregates, as opposed to single-chain motion.

**Fig. 3.**
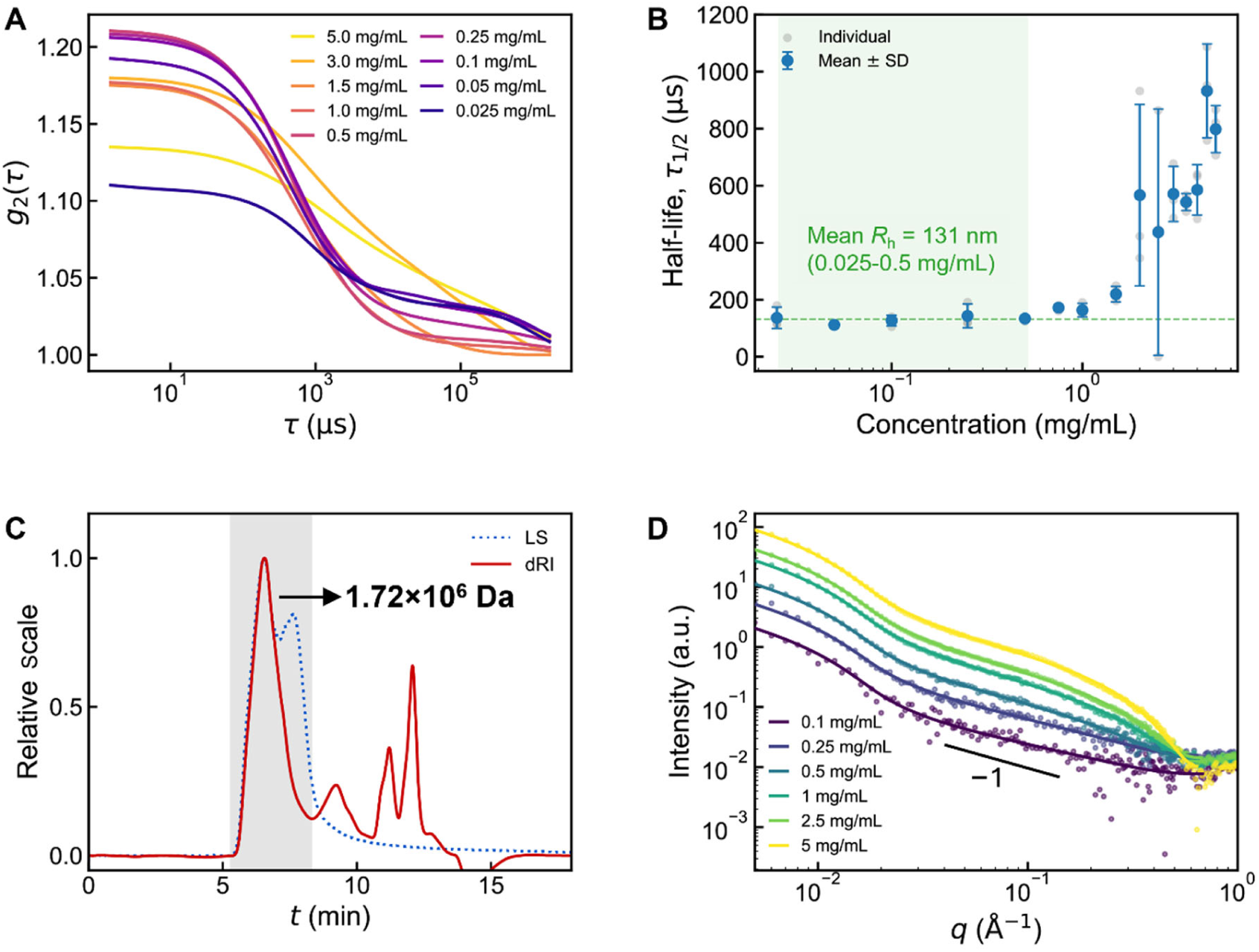
Characterization of VPS molecular weight, size, and conformation. (**A**) Autocorrelation functions of scattering intensity in DLS measurements for VPS solutions of different concentrations in PBS. For clarity, the average curve from three replicates is shown for each concentration. (**B**) Decorrelation time extracted from the autocorrelation functions across different VPS concentrations. The shaded region corresponds to the concentration range in which the calculated hydrodynamic radius corresponds to the size of one VPS molecule. (**C**) SEC-MALS analysis for estimating the molecular weight and radius of gyration of VPS. Shown are the light scattering intensity (LS, blue dashed line) and refractive index change (dRI, red solid line) as functions of time, measured in a Shodex LB-806M SEC column at a flow rate of 1 mL/min and a column temperature of 40°C. The shaded region corresponds to the peak from which the weight-averaged molecular weight was extracted to be 1.72 × 10^6^ Da (± 7.9%), with a dispersity of 1.05. (**D**) Scattering intensity, *I*, versus the magnitude of the scattering vector, *q*, for VPS solutions of different concentrations in PBS measured with SAXS, each fitted to a hybrid scattering model containing a Guinier–Porod scattering component and a cylindrical component. Solid line corresponds to a slope of −1.

### Molecular weight and radius of gyration of VPS, as revealed by SEC-MALS

Another critical feature of VPS that has not yet been characterized is its molecular weight. While the molecular weights of DNA and proteins are largely predetermined by their sequences (ignoring potential chemical modifications), the molecular weights of exopolysaccharides cannot be similarly inferred from the genome. Exopolysaccharides are generally synthesized by sets of enzymatic biosynthesis clusters from repeating units^51,52^; however, the mechanisms by which cells regulate their size remain largely unknown, as is the case for VPS. We performed size exclusion chromatography coupled with multi-angle light scattering (SEC-MALS) to determine both the weight-average molecular weight, *M*_w_, and the number-average molecular weight, *M*_n_. Note that SEC-MALS yields estimates of absolute *M*_w_ and *M*_n_, as opposed to values relative to a reference sample^53^. The resulting refractive index change (dRI) trace shows multiple peaks due to salts and other impurities (and potentially some low-molecular-weight fractions), while the light scattering (LS) trace shows a single major peak that colocalizes with the major peak in the dRI trace (Fig. 3C). Analyzing this peak yields an *M*_w_ value of 1.72 × 10^6^ Da (± 7.9%). This value is on the higher end of previously reported values for *M*_w_ of bacterial exopolysaccharides^54^. The measured number-average molecular weight, *M*_n_, is 1.64 × 10^6^ Da (± 7.9%), which leads to a dispersity (*M*_w_/*M*_n_) of 1.05. This is remarkably narrow compared to conventional synthetic polymers, as well as other exopolysaccharides (e.g., *S. epidermidis* PIA^34^ was reported to exhibit a dispersity of ∼ 2.8), and on par with values reported for some synthetic polymers obtained using living polymerization^55,56^. This strongly suggests that *V. cholerae* cells utilize some unknown mechanism to tightly regulate VPS chain length, which warrants further study. Finally, we estimated the average number of tetrasaccharide units along each VPS chain using the measured *M*_n_ value. The VPS tetrasaccharide unit (Fig. 1A) has a molecular weight of 760.6 Da without the O-acetyl group on the L-gulose moiety and 802.7 Da with the O-acetyl group; both forms exist in the VPS sample^27^. Hence, we estimate that, on average, a single VPS chain will contain around 2,100 tetrasaccharide units.

The SEC-MALS measurement also gives a radius of gyration *R*_g_ of 105 nm (± 5.4 %). This value is consistent with the hydrodynamic radius, *R*_h_, measured via DLS (∼ 130 nm). Assuming that the pervaded volume of each VPS chain is roughly spherical, we can then calculate a theoretical overlap concentration, *c*\*, as^33^

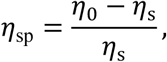

where *N*_A_ is the Avogadro number. This calculation yields a value of *c*\* ∼ 0.6 mg/mL, on the same order of magnitude as but slightly lower than the crossover concentration measured using *η*_sp_ (Fig. 1D).

The measured *R*_g_ value leads to another interesting question: the length of each VPS tetrasaccharide unit is around 1.8 nm head-to-tail, so the contour length of a VPS chain with 2,100 units is around 3.8 µm. Therefore, the VPS chain must accommodate a certain level of flexibility to achieve an *R*_g_ value of ∼ 105 nm. To investigate this, we turned to small-angle X-ray scattering (SAXS) to reveal the molecular conformation of the VPS chains.

### SAXS characterization reveals VPS as a semiflexible polymer with high persistence length

We performed SAXS measurements of VPS polymers both in water and in PBS, and we obtained qualitatively similar results between the two solvents (Figs. 3D and S3). All SAXS patterns exhibited two features: a Guinier–Porod decay at low *q*, where *q* is the magnitude of the scattering vector; followed by a scaling of *I*(*q*) ∼ *q*^‒1^ spanning more than one decade from intermediate to high *q*, suggesting that VPS molecules assume rigid, “rod-like” configurations at small length scales.

To resolve the detailed structure of VPS, we fitted the SAXS data to a hybrid model, containing both a Guinier–Porod component^57,58^ and a cylindrical component^59^ (see Methods). Due to the limited *q*-range in the low-*q* region, we focused on the fitting results in the high-*q* region, which corresponds to the local molecular conformation of VPS. Indeed, we found that, at these small length scales, the VPS conformation is best described as a rod with length ∼ 58 nm, which is presumed to be the persistence length *l*_p_ of the VPS molecules. We found that data obtained across VPS concentrations could be well-fitted to the hybrid model, with differential contributions from the two components. Importantly, the SAXS curve dramatically changes upon digestion of VPS using the native polysaccharide lyase from *V. cholerae*^27^ (Fig. S4A and B), rendering a significantly reduced rod length of ∼ 1.8 nm in the digested sample, close to the end-to-end distance of a tetrasaccharide unit. This outcome indicates that the *q*^‒1^ scaling in the high-*q* region indeed reflects the locally rigid conformation of the VPS chains. As another test, we filtered the VPS solution with a solid-phase extraction column to remove the large polymers, and indeed we found that the *q*^‒1^ scaling and associated spectral features disappear (Fig. S4C).

Altogether, the SAXS characterization shows that VPS molecules are rigid on length scales up to tens of nanometers, which is smaller than but not so far from the size of the entire molecule, as measured with DLS or SEC-MALS. To explain this behavior, we turned to multiscale molecular dynamics (MD) and Monte Carlo (MC) sampling.

### All-atom MD simulation and coarse-grained modeling of VPS segments

To examine how the chemical structure of VPS gives rise to its larger-scale conformational properties, we turned to all-atom MD simulations (Figs. 4A and S5A). Specifically, we simulated the dynamics of a VPS segment consisting of 10 tetrasaccharide units in explicit aqueous solvent with the Amber24 software package^60^, using the GLYCAM-06j force field for carbohydrates^61^ (three replicate trajectories, 500 ns each; see Methods). These simulations revealed that the VPS segment adopts a mostly linear conformation, which is qualitatively consistent with the large rod length obtained from the SAXS measurements; the VPS segment also displays a certain level of helicity. Quantifying the ф (O5–C1–O4’–C4’) and *ψ* (C1–O4’– C4’–C3’) dihedral angles along the four glycosidic linkage types revealed that each linkage preferentially assumes one highly stable conformation (Fig. S5B, Table S2). However, we also observed that the β-(1 → 4) linkage between the two glucose residues exhibits a discernible anti- *ψ* conformation (Fig. 4B; ф = —80° ± 4° , *ψ* = —6*η*° ± 4°; major conformation: ф = —71° ± 9°, *ψ* = 130° ± 13°), with an empirical probability of around 2.3% (9.5% including all nearby metastable conformations; Table S2). Visually examining snapshots of the simulations revealed that this minor conformation gives rise to sharp kinks along the contour of the VPS segment (Fig. 4C).

**Fig. 4.**
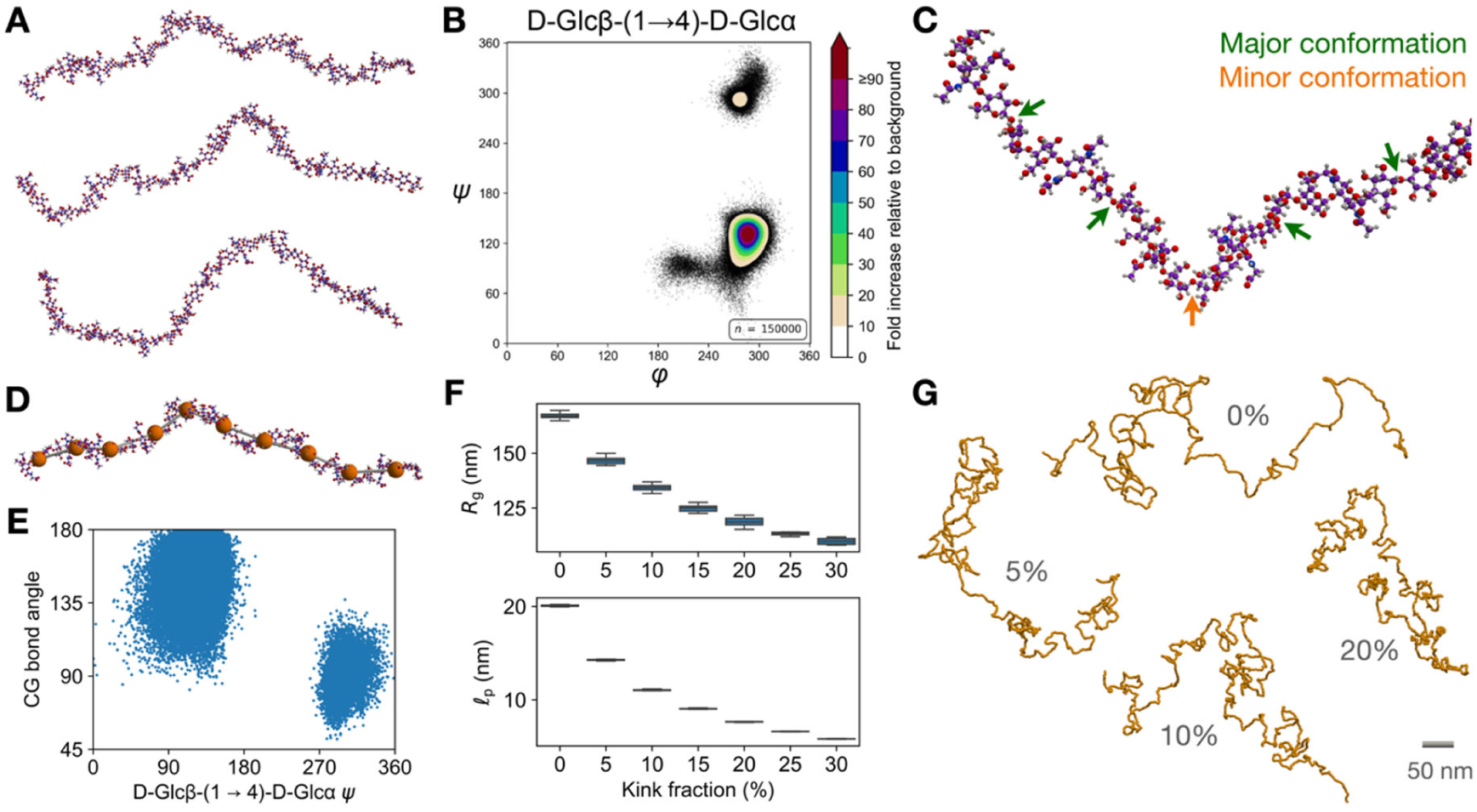
MD simulations and coarse-grained modeling of VPS chains. (**A**) Snapshots of an all-atom MD simulation trajectory of a VPS segment consisting of 10 tetrasaccharide units. See also Fig. S5A. (**B**) Contour map of the ф (O5–C1–O4’–C4’) and *ψ* (C1–O4’–C4’–C3’) dihedral angles along the D-Glcβ-(1 → 4)-D-Glcα glycosidic linkage, pooled over all three replicate trajectories. This plot reveals one major and one minor (anti-*ψ*) conformation. See also Fig. S5B and Table S2. (**C**) Representative snapshot of a portion of the VPS segment, with the D- Glcβ-(1 → 4)-D-Glcα linkages labeled according to their conformation. (**D**) Coarse-graining of the all-atom 10-tetrasaccharide VPS segment into a chain of 10 beads (orange), each representing a tetrasaccharide. Each bead is positioned at the center-of-mass of the atoms belonging to the corresponding tetrasaccharide. (**E**) Scatterplot of the D-Glcβ-(1 → 4)-D-Glcα *ψ* dihedral angle vs. the “bond angle” in the corresponding coarse-grained VPS segment (“CG bond angle”), showing that a VPS tetrasaccharide unit in which the glycosidic linkage assumes the anti-*ψ* conformation exhibits a ∼ 90° angle with its neighboring tetrasaccharide units. (**F**) Distributions of the root-mean-square radius of gyration (*R*_g_) and persistence length (*l*_p_) as functions of the kink fraction, obtained from MC conformational sampling of a coarse-grained VPS polymer consisting of 2,100 units. Each distribution is obtained from 10 measurements, each taken from an independent conformational ensemble. Whiskers correspond to 1.5 times the interquartile range below and above the first and third quartiles, respectively. See Methods for details; see also Fig. S8. (**G**) Representative conformations obtained using the MC sampling procedure for different kink fractions. For each kink fraction, the sampled conformation whose radius of gyration is closest to the corresponding mean value in panel **F** is shown. See also Fig. S7.

To investigate this connection more systematically, we developed a coarse-grained representation of the VPS segment as a chain of 10 beads, each representing a tetrasaccharide (Fig. 4D). The coordinates of each bead at each timepoint along each simulation trajectory were defined as the center-of-mass of each tetrasaccharide along the segment. We then related the *ψ* dihedral angles of the β-(1 → 4) linkages along the VPS segment to the corresponding “bond angles” along the coarse-grained chain, and observed that the major β-(1 → 4) conformation gives rise to a large bond angle (∼ 160°), whereas the anti-*ψ* conformation gives rise to a ∼ 90° kink (Fig. 4E). This mapping suggests that the conformational flexibility of the β-(1 → 4) glycosidic linkage between the two glucose residues may directly modulate the larger-scale conformational dynamics of the VPS chain.

### Coarse-grained conformational sampling of single VPS chains

The MD simulations suggested a putative mechanism by which a mostly rigid polymer segment can fold into a compact state through spontaneous rotation of a single glycosidic linkage. Since it is computationally infeasible to implement an all-atom MD simulation of a typical VPS chain consisting of 2,100 tetrasaccharide units (as inferred from our SEC-MALS measurements), we examined this idea more closely with a Monte Carlo (MC) conformational sampling procedure.

First, we quantified the empirical distributions of bond lengths, bond angles, and dihedral angles along the coarse-grained chain from the three replicate MD trajectories (Fig. S6A). This revealed that the bond lengths and bond angles both follow a multimodal distribution, each with a clear major peak: the bond lengths were concentrated around ∼ 1.8 nm, close to the end-to-end length of a tetrasaccharide unit, and the bond angles exhibited a major peak at ∼ 160°, as described above. Meanwhile, the dihedral angles exhibited a single broad peak at ∼ 145°, reflecting the helicity of the VPS segment, which also contributes to the stiffness of the VPS polymer. Fitting appropriately defined mixture models to these distributions (see Methods), we found that the major peaks in the bond length and bond angle distributions accounted for ∼ 90% of each observable, and that they were quite narrow: for instance, fitting the bond angle distribution to a folded two-component von Mises mixture revealed a bending stiffness of ∼20*k*_B_*T* (Fig. S6A, Table S3; see Supplementary Information).

To build an initial understanding of how these statistics dictate the conformation of a typical VPS chain, we combined these statistics with Svaneborg and Everaers’ modified Kremer–Grest model for a semiflexible polymer^62,63^ (see Supplementary Information). This model predicts that a semiflexible polymer with a bond length of 1.8 nm and a bending stiffness of 20*k*_B_*T* exhibits a Kuhn length of ∼ 70 nm (hence a persistence length of ∼ 35 nm), which is on the same order of magnitude as the rod length of ∼ 58 nm obtained from the SAXS measurements. However, this model, which assumes a smoothly changing director along the semiflexible polymer chain, cannot capture the consequences of a bimodal distribution of bond angles along the VPS polymer, nor the helicity of the polymer (Fig. S6A). To fill this gap, we implemented an MC conformational sampling procedure (Fig. S6B–F; see Methods and Supplementary Information). Here, we used a configurational-bias MC approach^64^ to iteratively sample conformations of a polymer consisting of 2,100 beads from the canonical ensemble (at 300 K), with bond lengths following a FENE potential^65^, bond angles following a bimodal Gaussian mixture potential^66^, and dihedral angles following a shifted harmonic potential. In particular, the weights of the two components in the bimodal angle potential—one centered at 160°, the other centered at 90°—control the “kink fraction” of the polymer, i.e., the fraction of bond angles along the chain that are near 90°. We sought to understand how the polymer’s dimensions are affected by this fraction.

We found that introducing even a small number of kinks along the polymer profoundly alters its conformation, with significant reductions in both *R*_g_ and *l*_p_ (Fig. 4F). Visually examining the sampled conformations revealed that, while a polymer with no kinks tends to exhibit an extended conformation, even a kink fraction of 5% along the polymer yields a significantly more compact conformation, and this effect increases with the kink fraction but quickly saturates (Figs. 4G and S7). In particular, we note that the *R*_g_ value we observe in the absence of kinks (*R*_g_ ≈ 170 nm) significantly exceeds our experimental estimate of ∼ 105 nm for *R*_g_ . This suggests that, given the intrinsic stiffness of the chain (Fig. S6A), kinks are necessary to achieve the experimentally observed degree of compaction. However, the *l*_p_ value we observe in the absence of kinks (*l*_p_ ≈ 20 nm) is shorter than the ∼ 58 nm rod length we obtained from the SAXS data; whether these values are directly comparable is unclear (see Discussion). In addition, we found that removing or strengthening the dihedral potential can also significantly change *R*_g_ and *l*_p_, but the qualitative effect of the kinks on *R*_g_ and *l*_p_ remains the same as we modulate the dihedral stiffness (Fig. S8). In sum, these findings suggest that even a small degree of conformational flexibility at the molecular scale (namely at the scale of a single glycosidic linkage) can dramatically alter the conformational statistics of single VPS chains at a much larger scale.

### Coarse-grained conformational sampling and analysis of entangled VPS networks

Our preceding analysis identified two structural features—the kink fraction and the stiffness of the dihedral angle potential—that dictate the conformations of single VPS chains (Figs. 4F and S8). We then asked how these two features jointly affect how multiple VPS chains interact and entangle with each other within a polymer solution. To do this, we adapted the MC conformational sampling procedure to multiple chains within a periodic domain (see Methods and Supplementary Information). Namely, we sampled conformations of semidilute polymer solutions, comprising multiple 200-bead chains situated within a periodic box, with the number of chains set to enforce a particular mass concentration *c*, using the same configurational-bias MC approach as in the single-chain analysis, but with an additional MD-based pre-equilibration step to initialize the system. We limited chain lengths to 200 beads to increase the number of unique chains within the system while maintaining a relatively small periodic box. We then varied the kink fraction (0%, 5%, 10%, 15%, and 20%); the dihedral stiffness, *K*_dihedral_ (0.5*k*_B_*T*, *k*_B_*T*, 2*k*_B_*T*, and 10*k*_B_*T*); and the concentration, *c*, to examine how these variables modulate the degree of entanglement among the chains. We set *c* to multiples of *c*_0_ = 6 mg/mL, which is slightly above the overlap concentration for a random coil of 200 beads with the same non-bonded interaction and FENE potentials as in the coarse-grained VPS model.

Visually inspecting the resulting configurations (Figs. 5A and S9A), we found that VPS-like chains for all choices of kink fraction, dihedral stiffness, and concentration assumed significantly interpenetrating conformations, whereas solutions of random coils of the same length at the same concentrations did not. To quantify the extent of entanglement in these configurations, we then used the Z1+ package^67^ to identify the primitive path^68,69^ corresponding to each chain in each configuration (Fig. S9B). We also computed various quantities related to entanglement (Figs. 5B and S10), including the mean number of entanglements per chain, (*Z*⟩; the mean primitive path length, (*L*_pp_⟩; and the ratio (*L*_pp_⟩/*R*_e_, where *R*_e_ is the root-mean-square end-to-end distance of the chains; this ratio measures the average *tortuosity* of the primitive paths. These quantifications revealed that, while the coils exhibited little to no entanglement even at *c* = *ηc*_0_, the VPS-like chains exhibited significant entanglement at all concentrations we examined (Fig. 5B); this is expected due to the more extended conformations of the VPS- like chains, which enable inter-chain contact and interpenetration at lower concentrations. Interestingly, we also found that kinks can modulate the primitive path geometry in different ways, depending on the dihedral stiffness (Fig. 5B). When the dihedrals are weakly constrained (*K*_dihedral_ = 0.5*k*_B_*T*), (*L*_pp_⟩/*R*_e_ decreases monotonically with the kink fraction. We speculate that this arises from increased chain compaction owing to a lower persistence length (Fig. 4F); according to Odijk–Semenov theory^40,41,70^, decreasing the persistence length increases the extent to which a chain can fluctuate transversely to its confining tube, resulting in a larger tube diameter (see Discussion). According to this picture, as the kink fraction increases, each chain is confined within a thicker tube that accommodates more chain contour distance per unit length along the primitive path, in a manner increasingly analogous to tubes in the flexible regime.

**Fig. 5.**
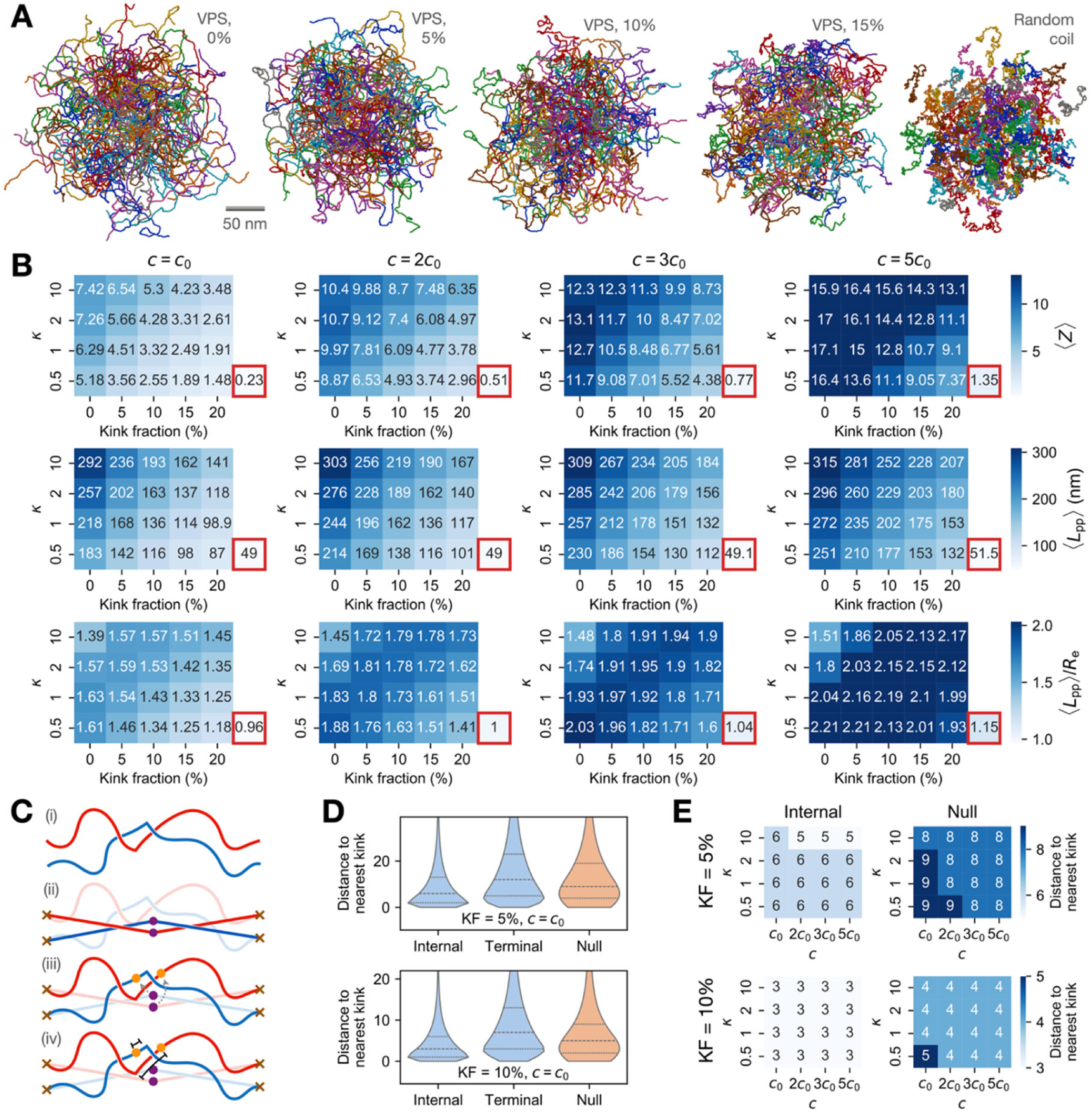
Coarse-grained modeling and entanglement analysis of VPS polymer solutions. (**A**) Representative configurations of polymer solutions at *c* = *c*_0_. Each chain is plotted such that its center-of-mass lies in the fundamental unit cell in the periodic domain. (**B**) Mean number of entanglements per chain, (*Z*⟩; the mean primitive path length, (*L*_pp_⟩; and the ratio (*L*_pp_⟩/*R*_e_, for each kink fraction, dihedral stiffness ( n = *K*_dihedral_/(*k*_B_*T*) ), and concentration ( *c* ). The corresponding mean values for random coils of the same length at the same concentration are given in the red boxes. Each mean value was obtained from a pooled ensemble of configurations from 10 independent MC runs. (**C**) Schematic of node-kink distance calculation. Given two entangled polymer chains (i), we can obtain their primitive paths using Z1+ (ii), with nodes shown in purple; we then identify the bead along the chain contour (orange) corresponding to each node in the primitive path (iii), then calculate the contour distance between this “node- associated bead” and the nearest kink (iv). (**D**) Node-kink distance distributions for *c* = *c*_0_, n = 0.5, and kink fractions of 5% and 10%. Distributions corresponding to internal nodes and terminal nodes are shown separately. Dashed lines within violins indicate the first, second, and third quartiles; the tail of each violin has been truncated for clarity. (**E**) Median node-kink distances for kink fractions of 5% and 10% and the indicated values of *c* and n, calculated in the same way as in panel **D**. Only internal nodes were used to calculate the medians on the left.

However, when the dihedrals are more strongly constrained, we found that (*L*_pp_⟩/*R*_e_ can change non-monotonically with the kink fraction: introducing a small number of kinks yields an increase in (*L*_pp_⟩/*R*_e_, indicating more tortuous primitive paths, but at higher kink fractions, (*L*_pp_⟩/*R*_e_begins to slowly decrease. We observed this non-monotonicity for dihedral stiffnesses as low as *K*_dihedral_ = *k*_B_*T*, albeit only at higher concentrations (*c* ≥ 3*c*_0_), suggesting that the crossover between the monotonic and non-monotonic regimes occurs in a concentration-dependent manner. We speculate that, in this latter regime, the dihedrals impart sufficiently high stiffness to the chains such that, when the kink fraction is small but nonzero, the increased tortuosity of the chains gives rise to increased tortuosity in their primitive paths, rather than an increased tube diameter.

To measure primitive path tortuosity more directly, we quantified the angles formed by the “nodes” along each primitive path, i.e., the points at which the primitive path changes direction. While VPS-like chains with no kinks exhibited long primitive paths with minor deflections that largely trace the end-to-end line segment, we found that increasing the kink fraction gives rise to progressively larger deviations in the primitive path’s direction (Fig. S10A and B). By contrast, we found that random coils exhibit short primitive paths that often do not deviate at all from the end-to-end line segment, as expected from their relative lack of entanglement. These results suggest that kinks can significantly reshape the tube confining each polymer, yielding—to an extent that depends on the dihedral stiffness and concentration—a tortuous tube geometry that qualitatively differs from those typically encountered in the flexible and semiflexible settings.

We then asked if the kinks directly modulate the locations of entanglements along the VPS-like chains. In particular, we hypothesized that introducing kinks along each chain causes the emergence of local geometries in which neighboring chain segments are more likely to be entangled. We reasoned that, if this were true, then these kinks would be colocalized with the “nodes” along the corresponding primitive paths—i.e., the points at which each primitive path changes direction—to a greater extent than would be expected by chance. To test this idea, we used the primitive paths obtained from Z1+ to map each node along each primitive path to its corresponding bead along the corresponding chain, which can be interpreted as the approximate location of the entanglement; then computed the distance along the chain contour between this bead and the nearest kink (Fig. 5C). Indeed, we found that these “node-associated beads” were generally closer to kinks than would be expected by chance (Fig. 5D and E; see Methods), consistent with our hypothesis. Note that this is only true for beads associated with internal nodes, which, unlike terminal nodes, represent genuine entanglements.

Finally, to ascertain the effects of kinks on the mechanics of the polymer network, we sought to estimate the entanglement length, *N*_e_, as a function of kink fraction and concentration (with *K*_dihedral_ fixed to 0.5*k*_B_*T*) by evaluating the “M-coil” and “M-kink” estimators^71^. To do this, we first generated additional configurations of solutions comprising shorter chains (50, 100, and 150 beads) for each choice of kink fraction and concentration, using the same procedure as described above; and we calculated the M-coil estimator as the value *n* such that

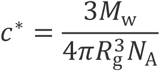

where (*l*_0_⟩ = 1.8 nm is the mean bond length and *L* is the contour length, and the M-kink estimator as the derivative *dN*/*d*(*Z*⟩ (see Methods). These quantifications revealed that introducing kinks increase *N*_e_ at lower concentrations (*c* = *c*_0_ and *c* = 2*c*_0_), but this effect diminishes as the concentration increases (Fig. S10C). This observation is qualitatively consistent with recent studies of the dependence of *L*_pp_ and *N*_e_ on chain stiffness in entangled polymer melts^63,72^. Estimating *N*_e_ at higher values of *K*_dihedral_, using the simpler “modified S- kink” estimator^71^, revealed similar trends (Fig. S10D).

## DISCUSSION

In this study, we used a combination of biophysical experiments and computer simulations to analyze the polymeric properties of VPS, the major matrix component in biofilms formed by the model pathogen *V. cholerae*. We obtained quantitative measurements for the molecular weight, radius of gyration, and persistence length, among other properties, all of which are critical for understanding the molecular basis of the rheological properties of VPS and the biofilm. By combining SAXS with multiscale simulations, we demonstrated that VPS is an intrinsically stiff polymer with a high persistence length but can also attain significant compaction through the spontaneous rotation of a glycosidic linkage within the VPS tetrasaccharide unit. Finally, we used MC simulations to demonstrate that this conformational flexibility may lead to a nonclassical mode of entanglement, not observed in flexible polymers or more conventional semiflexible polymers, at least in some regions of the underlying parameter space.

Our rheological measurements revealed that VPS behaves as a weakly charged polymer whose viscoelastic properties are sensitive to ionic screening by counterions in the solution. The exact nature of the crossover in the *η*_sp_ vs. *c* curve (Fig. 1D) is still unclear. We note that deviations from classical theories for polymer dynamics are common for polysaccharides such as dextran^73^, chitosan^74^, and hyaluronan^75^. We also note that the accuracy of our estimation of the scaling exponents is limited by the quality of our estimate of *η*_sp_, which is itself limited by the quality of our extrapolation-based estimate of *η*_0_, as well as the low torque limit of the shear rheometer, particularly with the small sample volume we are limited to. With these caveats in mind, we do note that the scaling exponents we observe beyond the crossover concentration, *c*^∗^, are roughly consistent with recent work by Lang and Frey on the dynamics of entangled semiflexible polymers^38^, in which they proposed a new scaling law for the terminal relaxation time, τ_R_, given by

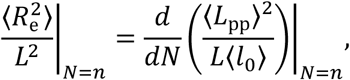

where ξ is the mesh size; according to a simple geometric argument, the mesh size scales as ξ ∼ *c*^−1/2^ in the semiflexible setting. We can now integrate this with Odijk–Semenov theory^40,41^, which predicts that the entanglement length, *N*_e_, of a semiflexible chain scales with the deflection length, *λ*, which in turn scales as *λ* ∼ *d*^2/3^*l_p_*^1/3^, where *d* is the tube diameter; the tube diameter, in turn, scales as *d* ∼ ξ^6/5^*l_p_*^−1/5^ , following Morse’s binary collision approximation^38,70,76,77^. Putting these pieces together, we first obtain a scaling relation for the plateau modulus^70,78^,

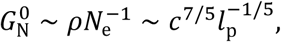

which we can then combine with Eqn. 1 to get

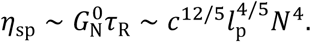

The resulting concentration scaling exponent (2.4) lies in between our measured exponents for VPS solutions in water (2.07) and in PBS (3.14). Alternatively, applying Morse’s effective medium approximation^70^, which implies that *d* ∼ ξ ∼ *c*^−1/2^ , yields a slightly different concentration scaling, *η*_sp_ ∼ *c*^7/3^ . Either way, this analysis suggests that the molecular processes underlying the Lang–Frey scaling—namely rotational diffusion of polymer chains following constraint release and disentanglement, as opposed to reptation along the tube^38^— may constitute the dominant mode of polymer dynamics in VPS solutions beyond *c*^∗^.

Regardless of the precise molecular mechanism, the crossover concentration (about 1 mg/mL, depending on the solvent) constitutes a quantitative threshold for the formation of a VPS polymer network. Interestingly, we found that *V. cholerae* biofilms produce and retain VPS at a concentration in the extracellular space near or slightly above the crossover concentration. This is relevant because it means that, even in the absence of any matrix proteins for crosslinking, VPS chains already begin to interact and form a percolating viscoelastic network. The high concentration of VPS in the extracellular space could potentially allow for VPS-interacting matrix protein molecules to bind multiple VPS chains^21,79^, leading to VPS crosslinking and facilitating the formation of higher-order structures in the *V. cholerae* biofilm matrix^20^.

Previously, we took a different and complementary approach for understanding biofilm mechanics, by measuring the rheological properties of the entire biofilm^48,49^. Those studies showed that the biofilm-dwelling cells also contribute significantly to biofilm mechanics: on the one hand, bacterial cells are much more rigid (with a modulus on the order of 30 MPa; ref. ^80^) compared to the polymer network. On the other hand, the cell-surface-anchored VPS molecules being actively secreted by the cells^26^, as well as the VPS-containing outer membrane vesicles in the matrix^81^, can engage in further interactions with the VPS network. As a consequence, the *G*′ and *G*′′ values measured for the biofilm are generally much larger (*G*′_p_ > 100 Pa) than those for the VPS solutions. How to consider the contributions of the biofilm-dwelling cells to biofilm mechanics is an interesting question for future studies; we suggest that drawing connections to recent theoretical work on the mechanical properties of disordered polymer networks with rigid inclusions^82,83^, as well as the broader study of polymer– nanoparticle composites^84,85^, may be illuminating.

Our MD simulations revealed that VPS attains conformational flexibility through both the intrinsic flexibility along its backbone and large, but infrequent, rotations of the β-(1 → 4) glycosidic linkage between the two central glucose residues within the VPS tetrasaccharide unit, which give rise to transitions between a dominant conformation and a rare anti-*ψ* rotamer for each linkage. A similar phenomenon has been observed in a previous simulation study of amylose chains, wherein transitions between a dominant conformation and an anti-*ψ* rotamer of the α-(1 → 4) glycosidic linkages along the chain causes it to collapse into a coiled, disordered state^86^. Our findings in this paper demonstrate that the flexibility of the β-(1 → 4) linkages between the two D-glucose units plays a somewhat analogous role in disrupting the ordered structure of each VPS tetrasaccharide unit and facilitating the compaction of VPS chains; this gives rise to a polymer that contains rod-like segments that are tens of nanometers long but exhibits a radius of gyration on the order of 100 nm. While our experimental and simulation-based measurements of *R*_g_ and *l*_p_—assuming a kink fraction of 5–10%—do not match exactly, they fall within the same order of magnitude; the discrepancy may be attributed to experimental measurement error, imprecise estimation of conformational probabilities from the all-atom simulations, or the loss of other potentially important molecular details, such as electrostatic interactions, in our coarse-grained simulations.

We also discovered that these transient kinks, together with the stiffness of the dihedral angles along each chain, can change how multiple chains interact with each other in a semidilute solution. Specifically, we found that introducing a small number of kinks can change the geometry of each chain’s confining tube by increasing its tortuosity, to an extent that also depends on the dihedral stiffness and concentration. Consistent with this effect, we also found that the kinks are often colocalized with the positions at which chains are entangled, suggesting that the kinks give rise to local geometries in which chains exhibit a greater propensity for entanglements. While a quantitative correspondence between our rheological measurements and simulations is yet to be achieved, our results indicate that these kinks constitute one plausible mechanism through which VPS chains can interact to form a network.

We also found that kinks mostly increase the entanglement length, *N*_e_ , in a concentration-dependent manner; we speculate that this arises from the decreased *l*_p_ of the kinked polymers, which has been observed to result in increased *N*_e_ in semiflexible polymer melts^63,72^. This, in turn, suggests that introducing kinks decreases the plateau modulus, *G*_N_^0^, which is related to *N*_e_ as 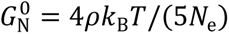 = 4*ρk*_B_*T*/(*ηN*_e_) (refs. ^68,69^). The implications of these findings for the viscosity of the VPS network are ultimately unclear, because *η*_sp_ depends not only on *G*_N_^0^ but also on the terminal relaxation time, τ_R_. We speculate that the kinks’ effects on the geometry of the tube confining each chain should in turn modulate this timescale; further work is required to reveal the precise nature of this effect and, more generally, whether entanglements between kinked polymers exhibit different dynamical behaviors from those involving more traditional semiflexible polymers^38^. Specifically, how does the increased tortuosity of a kinked polymer chain’s primitive path affect the chain’s ability to reptate within its confining tube? How are the tube geometry and reptation dynamics coupled with the interconversion between the linear and kinked conformations, and could this interconversion constitute a new form of constraint release? Upon shear, do neighboring kinks introduce intercalations between their respective polymer chains, further restricting chain motion? If so, how do such intercalations differ from more conventional entanglements (i.e., slip-links^87^) in their ability to transmit tension throughout the polymer network? Combined theoretical and experimental work is now underway in the lab to explore these tantalizing questions.

Placing our results back into the biological context, we note that, in growing *V. cholerae* biofilms, the VPS polymers are produced and secreted by the biofilm-dwelling bacteria in a non-equilibrium process: unlike in our simulations, their numbers and total occupied volume continually change throughout biofilm growth. A possible evolutionary advantage of the conformational flexibility we have detailed here is that it may allow VPS molecules to be secreted and incorporated into the extracellular matrix in a temporally coordinated manner: they are secreted as highly extended, close-to-linear polymers from their biosynthesis machinery, after which they immediately assume a more compact conformation and engage in entanglements with neighboring VPS chains before they can diffuse into the bulk solution. This hypothesis is consistent with our measured VPS concentration within biofilms being above the crossover concentration; it is also consistent with previous reports that VPS-producing cells are able to maintain and prioritize the VPS molecules they have secreted themselves, and prevent non-VPS-producing “cheater” cells from taking advantage of the secreted VPS^26,88,89^.

Our findings in this paper, which focused on the characterization of VPS by itself, lay the groundwork for understanding how *V. cholerae* biofilms derive their mechanical properties from VPS in concert with the key matrix proteins, RbmA, Bap1, and RbmC. An intriguing point of connection in this regard is the interaction between VPS and the β-propeller shared between RbmC and Bap1, which we have recently characterized with X-ray crystallography. This crystal structure revealed that a β-propeller-bound VPS tetrasaccharide unit adopts a fully kinked conformation, in which the β-(1 → 4) linkage between the glucose residues is bent in a similar fashion as in the minor conformation in Fig. 4B. In other words, binding to the β-propeller shifts the energy landscape of this glycosidic linkage, stabilizing the minor conformation in unbound VPS. We predict that such conformational changes that arise within VPS upon binding to different matrix proteins will dramatically modulate the mechanical properties of the VPS network and consequently the mechanics of the entire biofilm; further work will be required to dissect these effects in detail.

## METHODS

### Strains, media, and materials

All *V. cholerae* strains used in this study are derivatives of the wild-type *Vibrio cholerae* O1 biovar El Tor strain C6706 and are listed in Table S1. All strains harbor a missense mutation in the *vpvC* gene (*vpvC*^W240R^), which elevates intracellular cyclic diguanylate level and thus facilitates constitutive biofilm formation^90^. Additional mutations were genetically engineered using natural transformation^91^ unless indicated otherwise. In general, strains were grown overnight in lysogeny broth (LB) at 37°C with shaking. M9 minimal media (Sigma Aldrich) was supplemented with 2 mM MgSO4 (JT Baker) and 100 µM CaCl2 (JT Baker) (henceforth referred to as M9 medium).

### VPS purification

VPS purification was performed using a previously published protocol with several modifications^16,25^. First, a Δ*rbmA*Δ*bap1*Δ*rbmC*Δ*pomA* strain was grown in LB at 30°C overnight. 50 μL of this inoculum was added into 3 mL of LB liquid medium containing glass beads (4 mm, MP Biomedical), and the cultures were grown with shaking at 30°C for 3–3.5 h. 50 μL of this inoculum was applied to an agar plate containing M9 medium supplemented with 0.5% glucose and 0.5% casamino acids. The plates were incubated at 30°C for 2 days to form a continuous bacterial lawn. For each batch, 20 plates were used. The biofilms were carefully scraped off the agar plates and resuspended in 1× phosphate-buffered saline (PBS) buffer. Cells were removed by centrifugation (8,000 × g, 4°C, 45 min) and the supernatant was dialyzed for 3 days against distilled water using a dialysis cassette (10 kDa MWCO) with repeated water changes. The dialyzed sample was lyophilized to prepare crude VPS extract. The crude extract was dissolved in 10 mM Tris buffer at 1.5 mg/mL, treated with DNase and RNase (37°C, 24 h), then treated with Proteinase K (37°C, 48 h), followed by ultracentrifugation at 100,000 × g for 1 h to remove lipopolysaccharides. This solution was dialyzed against water for 3 days and lyophilized to finally obtain purified VPS. For each purification batch, typically 10–15 mg of VPS was obtained as a white powder after the final lyophilization step. The VPS was dissolved in either Milli-Q water or PBS, and the solutions were heated at 95°C for 10 min to denature Proteinase K before use.

### Rheological measurements

All rheological measurements were performed with a stress-controlled shear rheometer (Anton Paar Physica MCR502WESP) at 25°C. For each measurement, a VPS solution sample at a given concentration was transferred onto the lower plate of the rheometer. Measurements were initiated after sandwiching the VPS solution between the upper cone plate and the lower plate with a minimal gap size of 0.05 mm. Oscillatory shear tests were conducted by varying the amplitude of the oscillatory strain, *ε*, from 0.01% to 2,000% at a fixed frequency of 6.28 rad s⁻¹. The storage modulus *G′* and loss modulus *G′′* were extracted using RheoPlus software as a function of *ε*. To determine the plateau storage modulus *G′*_p_, segmented linear fits were applied to *G′*(*ε*) curves on a log–log scale^92^. The strain value at the intersection of the two fitted regimes was defined as the yield strain, *ε*Y. In the low-strain plateau region, *G′* varied minimally, and the fitted value at *ε* = 1% was taken as the plateau modulus, *G′*_p_ (ref. ^92^). The yield stress *σ*Y was defined as the stress corresponding to *ε*Y. Frequency sweep tests were performed with angular frequency varying from 0.1 to 200 rad s⁻¹ at a fixed strain of 1% within the linear viscoelastic regime, from which the complex viscosity |*η\**| was obtained. The zero-shear viscosity *η*_0_ was estimated by extrapolating to the low-frequency regime, and the corresponding specific viscosity *η*_sp_ was subsequently calculated using the solvent viscosity *η*_s_. Three batches of VPS were purified and combined for rheological measurements, and 3–6 technical replicates were performed for each concentration.

### Biofilm growth, imaging, and analysis

The *V. cholerae* strain with all matrix protein genes knocked out (Δ*rbmA*Δ*bap1*Δ*rbmC*) and constitutively expressing mNeonGreen was first cultured overnight at 37°C with shaking in LB, after which the OD_600_ was measured. Next, the bacterial cultures were diluted to OD_600_ = 0.01 in an M9 medium supplemented with 0.5% glucose and 0.5% casamino acids. 100 µL of this mixture was transferred into a well of a glass-bottomed 96-well plate and kept at 25°C using an on-stage heater. Fluorescence microscopy was performed using a Yokogawa W1 confocal scanner unit connected to a Nikon Ti2-E inverted microscope with a Perfect Focus System. Cells were excited at 488 nm with the corresponding filter. All fluorescence signals were recorded with an sCMOS camera (Photometrics Prime BSI). Confocal images were taken using a 60× water immersion objective (CFI Plan Apo 60XC, numerical aperture = 1.20) after overnight biofilm growth. A *z*-stack was captured with a *z*-step size of 0.43 µm, over a *z*-range of −2 µm to 80 µm above the glass surface. For analysis, a small imaging volume (around 10 µm × 10 µm × 10 µm) away from the image edge and from the top or bottom of the biofilm was selected to avoid edge effects. The number of cells in the imaging volume was manually counted to obtain the volume per cell in the biofilm. The analysis was repeated with 14 randomly chosen regions, all of which yielded statistically indistinguishable results. All images presented are raw images rendered with Nikon NIS-Elements (version 5.20).

### Quantification of VPS in *V. cholerae* biofilms

VPS content was quantified using a previously reported glycosaminoglycan hexosamine assay^45^. In this method, 2,5-anhydrohexoses generated by deamination of hexosamines react with 3-methyl-2-benzothiazolone hydrazone hydrochloride (MBTH) to form colored complexes that can be quantitatively detected at 650 nm. A standard calibration curve was first established using glucosamine-HCl, and the results were consistent with those reported in the literature, confirming that the method is effective and reproducible. A VPS calibration curve was subsequently constructed using purified VPS samples. We observed deviations between the two calibration curves, which can likely be attributed to differences in the reactivity of glucosamine-HCl and VPS under the assay conditions.

Bacterial samples containing VPS, as well as control samples lacking VPS, were then analyzed and quantified using the VPS calibration curve. Briefly, the Δ*rbmA*Δ*bap1*Δ*rbmC* mutant was grown overnight in LB, and regrown statically in M9 medium supplemented with 0.5% glucose and 0.5% casamino acids at 25°C to an OD_600_ of 1.0. This sample was first centrifuged at 18,500 rcf for 5 min at room temperature in Eppendorf tubes to remove cells and debris, and the resulting supernatant was used for VPS quantification. In parallel, the same procedure was repeated with the Δ*rbmA*Δ*bap1*Δ*rbmC*Δ*vpsL* mutant as a negative control, to account for other molecules in the cell culture that may react with MBTH. Using this procedure, we estimated the VPS concentration in the Δ*rbmA*Δ*bap1*Δ*rbmC* sample as 0.081 ± 0.030 mg/mL. The relatively large error arises from the strong background signal of the control sample without VPS, which may originate from other glycosaminoglycan hexosamine species such as those in fragmented cell walls.

### SEC-MALS analysis

Prior to the SEC-MALS analysis, the sample was washed with an 80:20 (v/v) ethanol:water solution and vortexed for several minutes to ensure thorough mixing. The mixture was then stored in a refrigerator overnight to allow the water-soluble components to settle at the bottom.

The following day, the sample was centrifuged at 2,500 RPM, and the aqueous layer was carefully decanted. Residual ethanol was removed by placing the sample in a low-temperature oven until the solvent had fully evaporated. The dried material was subsequently re-dissolved in water, frozen at −80°C, and lyophilized (freeze-dried) to obtain the final product.

Samples were prepared for size exclusion chromatography (SEC) by dissolving 5 mg of sample in 1 mL of 0.100 M NaNO_3_ and were filtered through a 0.45 μm syringe filter. 0.25 μL of the sample was loaded onto a Shodex LB-806M SEC column at a flow rate of 1 mL/min and a column temperature of 40°C. The molecular weight was determined using the multi-angle light scattering (MALS) intensity and refractive index change (dRI), which were obtained via in-line Wyatt detectors. All data were analyzed with Astra 8 software and normalized to commercial PEG and PEO standards (Wyatt).

### Dynamic light scattering

Purified VPS was dissolved in 1×PBS buffer at concentrations ranging from 5.0 mg/mL to 0.025 mg/mL. Samples were briefly vortexed before being left on a nutating shaker overnight and subsequently heated to 95°C for 10 min to ensure thorough dissolution. All measurements were performed on the Wyatt DynaPro II dynamic light scattering (DLS) plate reader at *λ* = 658 nm using a Corning 384 well plate. Wells were cleaned by rinsing with ultrapure water followed by dusting with compressed air. 40 µL of each VPS sample, each with a different concentration, was loaded across three wells and measurements obtained for each sample. Autocorrelation function half-lives were obtained using the regularization analysis feature in the Dynamics DLS software and fitted to the one phase decay model using GraphPad Prism. The hydrodynamic radius was calculated using the cumulants analysis feature in Dynamics DLS software. Reported half-lives are averages of three technical replicates.

### Digestion of VPS

Digestion of VPS was performed using a previously published procedure^27^. Briefly, reactions were incubated at 30°C using 20 mM HEPES and 150 mM NaCl buffer overnight and quenched by heating to 95°C for 5 min followed by centrifugation at 21,000 × g for 5 min. Following cleavage, reaction mixtures were adjusted using 20 mM HEPES and 150 mM NaCl buffer to 0.56 mg/mL of VPS. 200 μg of VPS and 160 μg of RbmB, purified according to published protocols^27^, were used for the reaction in a total volume of 360 μL. An undigested sample in the same buffer (20 mM HEPES and 150 mM NaCl buffer) was included in the same set of experiments as a control.

### Small-angle X-ray scattering

Small-angle X-ray scattering (SAXS) experiments were conducted at the Life Sciences X-ray Scattering (LiX) beamline (16-ID) at NSLS-II at Brookhaven National Laboratory, using a dual-detector setup—Pilatus 1M (in air) and Pilatus 900K (in vacuum)—operating simultaneously to provide a broad, continuous scattering vector magnitude (*q*) range of 0.005< *q* < 3.2 Å^−1^. Data were collected at a wavelength, *λ*, of 0.819 Å, and the momentum transfer, *q*, was defined as *q* = 4*π* sin(*θ*/2) /*λ*, where *θ* is the scattering angle. Detector calibration was performed using silver behenate, with its well-defined 5.8 nm lamellar spacing. Samples were measured using a flow cell to reduce radiation damage, and background subtraction was performed using measurements from matching buffers. Different exposure times (0.5–1 s) were used for samples at different concentrations, and the corresponding controls were used for subtraction. Data were processed using the py4xs Python package. In one of the SAXS samples, the sample was purified using a graphitized carbon solid-phase extraction column (Sigma-Aldrich) to remove the polysaccharides.

### Analysis of SAXS data

The SAXS data were best fitted to a hybrid scattering model comprising a Guinier–Porod scattering component^57,58^ and a cylindrical component^59^. The Guinier–Porod model is given by

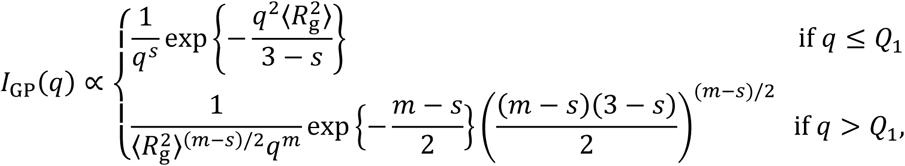

where 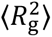, *m*, and *s* are the mean squared radius of gyration, Porod index, and dimensionality, respectively; and

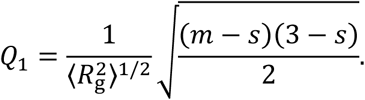

*s* = 0, 1, 2 correspond to the object in the Guinier regime being best described by a sphere, rod, or lamella, respectively. The cylindrical scattering model is given by

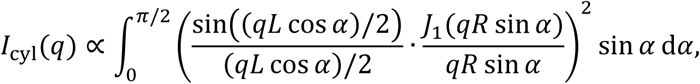

where *L* and *R* are the length and radius of the cylinder, and *J*_1_(·) is the first-order Bessel function of the first kind. The total scattering intensity, *I* (*q*), is a weighted sum of these two contributions:

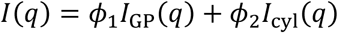

We used SasView 6.1.0 to fit the SAXS data and estimate the dimensions, *L* and *R*, of the rod-like structure.

### All-atom MD simulation of VPS segments

Initial coordinates for a 3D all-atom model of a 10-tetrasaccharide VPS segment were generated in three steps. First, a 3D structure for a related oligosaccharide with an unmodified L-gulose residue (L-Gulα), as a placeholder for the modified L-gulose present in VPS (L- GulNAcA(3Ac,6Gly)α), was generated using GLYCAM-Web^93^. Second, a molecular structure file in Amber PREP file format was generated for L-GulNAcAα, with ensemble-averaged atomic partial charges derived using procedures described previously for the GLYCAM-06j force field for carbohydrates^94^. The tleap module in the Amber software suite^60^ was then used to replace the L-Gulα coordinates with these coordinates. Finally, using tleap, a glycine moiety was attached via an amide linkage at the C6 position in L-GulNAcAα, and an acetyl moiety was attached at the O3 position, to create the required L-GulNAcA(3Ac,6Gly)α residue. Valence parameters associated with the amide linkage were assigned using the Generalized Amber Force Field^95^.

The resulting initial 3D structure was solvated in a periodic truncated octahedral box of TIP5P water^96^, with an 8 Å buffer and counterions added to neutralize the system using tleap. Energy minimization and MD simulations were carried out at 300 K following standard protocols^97^ with the Amber24 software suite^60^, using the GPU-accelerated implementation of particle mesh Ewald MD^98^ and the GLYCAM-06j force field^61^. Covalent bonds involving hydrogen atoms were constrained using the SHAKE algorithm^99^, enabling a timestep of 2 fs. The system was allowed to evolve for a total trajectory time of 500 ns, and frames were recorded every 100 ps for downstream analysis using the Amber CPPTRAJ program^100^. Three independent MD production runs were performed to generate averages and standard deviations.

### Coarse-grained model of VPS

We coarse-grained the all-atom MD simulations by defining a bead at the center-of-mass of each tetrasaccharide along the VPS segment, and tracking the beads’ positions over time (Fig. 4D). Here, the Python package MDTraj^101^ (version 1.11.0) was used to process the MD trajectories. The beads’ positions were then used to quantify empirical distributions of inter-bead “bond lengths,” “bond angles,” and “dihedral angles” (Fig. S6A). We fitted a two-component Gaussian mixture model to the bond length distribution using the Python package scikit-learn^102^ (version 1.4.1); this yielded a major peak at ∼ 1.8 nm, which is close to the end-to-end distance of a tetrasaccharide unit (Table S3). We fitted a symmetrically folded two-component von Mises mixture model to the bond angle distribution, using a custom implementation of a standard expectation maximization procedure^103,104^. More specifically, we “unraveled” the bond angle distribution—which is defined over a support of [0°, 180°)—onto a support of [—180°, 180°) by negating a random subset of half the angles, then fitted a four-component von Mises mixture model to this unraveled distribution. Noting that the resulting von Mises mixture is highly symmetric about zero, we then “folded” the mixture back onto [0°, 180°) by taking the two matching pairs of components about zero and averaging the means, concentrations, and weights within each pair, and renormalizing the weights of the two resulting components to sum to one. Finally, we fitted a simple von Mises distribution to the dihedral angle distribution, which, unlike the bond angles, are defined over a support of [—180°, 180°). The resulting fits are described in Table S3.

### MC conformational sampling of coarse-grained VPS conformations

To sample conformations from the canonical ensemble (at 300 K) of a single coarse-grained VPS chain, we implemented a configurational-bias Monte Carlo^64^ (MC) sampling procedure (Fig. S6B). Full details of this procedure are given in the Supplementary Information. Briefly, non-bonded pairs of beads were assumed to interact according to a truncated Lennard-Jones potential,

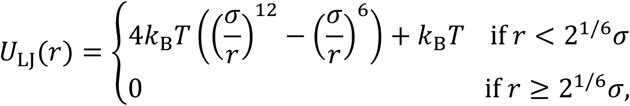

where *σ* is the interaction length scale; this is the repulsive component of the Weeks–Chandler– Andersen decomposition of the Lennard-Jones potential^105^. Bonded pairs of beads were assumed to interact according to a FENE potential^65^,

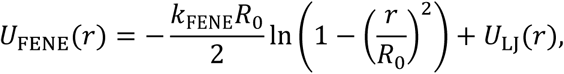

where *k*_FENE_ is a bond stiffness with units of energy/distance^2^ and *R*_0_ is a maximum bond length (at which the potential diverges). Bond angles were assumed to follow a two-component Gaussian potential^66^,

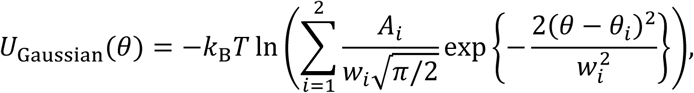

where *θ*_i_ , *w*_i_/2, and *A*_i_ are the mean, standard deviation, and weight of the *i*-th Gaussian component in the corresponding Boltzmann distribution, for *i* = 1, 2 . To model the spontaneous, sporadic emergence of kinks along the chain, we set *θ*_1_ = 160° (representing a nearly linear conformation) and *θ*_2_ = 90° (representing a kink), and set the weights *A*_1_ and *A*_2_ to reflect the relative frequency of kinks along the chain (the “kink fraction”), which we varied between zero and 0.3. Finally, dihedral angles were assumed to follow a single-welled harmonic potential,

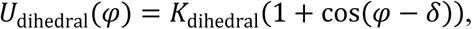

where the offset, *δ*, was set so that the potential is minimized at φ = 150°. Potential parameters were set to the values in Table S4, save for the weights *A*_1_ and *A*_2_ in the Gaussian angle potential.

For each MC run, an initial polymer configuration was first generated by randomly sampling bond lengths, bond angles, and dihedral angles from the Boltzmann distributions corresponding to their respective potentials, while enforcing a minimum distance of 2^1/6^*σ*— the cutoff distance for the truncated Lennard-Jones potential—between non-bonded pairs of beads. The configuration was then iteratively perturbed using three MC move types: reptation in either direction by a single bead, reptation in either direction by multiple beads, and deletion and regrowth of a terminal segment (Fig. S6B). Each move type was implemented using a configurational-bias Monte Carlo approach, wherein a collection of candidate moves were generated by sampling bond lengths, bond angles, and dihedral angles from the Boltzmann distributions corresponding to their respective potentials; one candidate move was chosen according to “residual” non-bonded interaction energies between the new beads in each candidate move and the remaining beads from the previous configuration; this move was then accepted or rejected with an acceptance probability that is defined so as to maintain detailed balance^64,106^. To ensure that the generated sample is representative of an equilibrium configurational ensemble, a burn-in of 100,000 configurations was discarded from each MC run, which far exceeded the number of iterations required for the configuration energy to apparently reach equilibrium. Configurations were collected every 500 iterations to reduce correlations between consecutive configurations in the sample; and a total of 999 configurations (excluding the initial configuration) were collected in each run. The sampling procedure was further validated by comparing the energies of an ensemble of sampled configurations for 100- bead and 200-bead chains against those obtained from corresponding MD simulations, which we implemented using LAMMPS^107,108^ (version 2Aug2023; Fig. S6C); by examining the distributions of bond lengths, bond angles, and dihedral angles from the sampled configurations, which exhibited the desired statistical properties (Fig. S6D); and by examining the energy distributions from configurations for the full 2,100-bead chain obtained from independent MC runs (Fig. S6E and F), which revealed minimal variation between runs, suggesting that they are sampling from a common equilibrium distribution.

### MC conformational sampling of VPS polymer solutions

To sample conformations of VPS chains in a polymer solution with a fixed concentration, we adapted the single-chain MC procedure to accommodate multiple chains. Full details of this procedure are provided in the Supplementary Information. Briefly, a fixed number of chains (72–120) were first situated within a periodic domain (with a cubic unit cell of appropriately chosen volume, based on the number of chains and the desired concentration), initially allowing for beads in different chains to be closer to each other than the non-bonded interaction cutoff distance (2^1/6^*σ*). The chains were then pushed apart by briefly equilibrating them (5 ns) in LAMMPS with a Langevin thermostat at 300 K, while applying a “soft” non-bonded interaction potential,

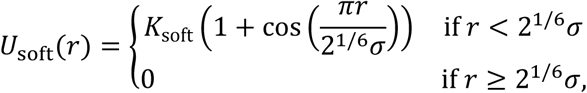

in lieu of the truncated Lennard-Jones potential. The prefactor, *K*_soft_, was gradually increased from 0 to 10*k*_B_*T* throughout this equilibration to avoid sudden jumps in the beads’ positions. Then, upon re-introducing the Lennard-Jones potential, the chains were energy-minimized and briefly re-equilibrated (5 ns), after which the resulting configuration was passed as input into the MC procedure. Each move in the MC procedure entailed randomly choosing one chain in the system, and applying one of the same three move types—single-bead reptation, multi-bead reptation, and terminal segment move—as in the single-chain procedure (Fig. S6B). Periodic boundary conditions were assumed throughout, and periodic inter-bead distances were used to calculate configurational energies and acceptance probabilities.

### Entanglement analysis

Entanglement-related properties were quantified using the Z1+ package^67^ (version 1), as described in the text. To generate the “null” distributions in Fig. 5D and E, we iteratively placed internal nodes (or, to be more precise, node-associated beads) and kinks along a 200-bead chain according to a uniform distribution, and calculated the distance between each node and its nearest kink. The number of nodes in each iteration was itself randomly sampled from the empirical distribution of the number of internal nodes ( *Z* ) along each chain in each configuration, whereas the number of kinks was fixed according to the kink fraction. The resulting distribution was then compared with the actual node-kink distance distributions for the given kink fraction and concentration.

To estimate the “M-coil” and “M-kink” estimators for the entanglement length *N*_e_ (ref. ^109^), additional configurations of solutions comprising 50-bead chains, 100-bead chains, or 150- bead chains were generated using the same MC procedure as described above. The M-coil estimator, 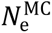, is defined as the value *n* such that

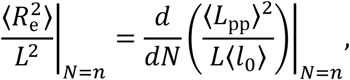

where (*l*_0_⟩ = 1.8 nm is the mean bond length and *L* is the contour length. To calculate the right- hand derivative, the values of (*L*_pp_⟩^2^/(*L*(*l*_0_⟩) obtained using Z1+ were fitted to a linear model of the form, (*L*_pp_⟩^2^/(*L*(*l*_0_⟩) = *mN* + *b* (*R*^2^ > 0.91), and the derivative was set to the slope, *m*, in the fitted model. Estimates for the left-hand quantity, 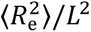, were then fitted to a decreasing function of the form *d* + *c*/(*N*— 1) , and 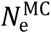 was finally set to 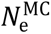 = 1 + *c*/(*m*— *d*). The M-kink estimator, 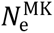, is defined as the derivative,

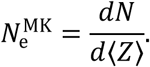

As in the M-coil calculation, the data were fitted to a linear model of the form, *N* = *m*(*Z*⟩ + *b* (*R*^2^ > 0.96), and 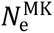 was set to the slope, *m*, in the fitted model.

### Statistical analysis

Errors correspond to standard deviations from measurements taken from distinct samples. All statistical analyses were performed using GraphPad Prism software.

## Acknowledgments

We thank Drs. Simon Mochrie and Robert Moore for helpful discussions. We acknowledge Caylyn McNaul from GlycoMIP for acquisition of the SEC-MALS data and Dr. Shirish Chodankar from Brookhaven National Laboratory for assistance with the SAXS experiments. The research reported in this publication was supported by the National Science Foundation awarded to J.Y. and Y. Li (DMR/MPS 2205006 and 2313746). J.Y. acknowledges support from the National Institute of General Medical Sciences of the National Institutes of Health (1DP2GM146253). Research was sponsored by the Army Research Office and was accomplished under Award Number: W911NF-25-1-0084 (to J.Y.). The VPS quantification was based upon work supported by the Air Force Office of Scientific Research under Award Number: FA9550-25-1-0293 (to M.Z.). The views and conclusions contained in this document are those of the authors and should not be interpreted as representing the official policies, either expressed or implied, of the Army Research Office or the U.S. Government. The U.S. Government is authorized to reproduce and distribute reprints for Government purposes notwithstanding any copyright notation herein. This work was supported in part by GlycoMIP, an NSF Materials Innovation Platform funded through Cooperative Agreement DMR-1933525 (R.J.W.). M.F.H. and M.-P.N. would like to thank the funding support for SAXS analysis from the NSF (DMR-2420040). This research used the LiX beamline at the National Synchrotron Light Source II, a U.S. Department of Energy (DOE) Office of Science User Facility operated for the DOE Office of Science by Brookhaven National Laboratory under Contract No. DE- SC0012704. The Center for BioMolecular Structure (CBMS) is primarily supported by the NIH, National Institute of General Medical Sciences (NIGMS) through a Center Core P30 Grant (P30GM133893), and by the DOE Office of Biological and Environmental Research (KP1607011). Beamtime was obtained through the Yale University Block Allocation Group #317429. All conformational sampling and simulations of coarse-grained VPS chains were performed on the Bouchet high-performance computing cluster, which is maintained by the Yale Center for Research Computing. This work benefited from the use of the SasView application, originally developed under NSF award DMR-0520547. SasView contains code developed with funding from the European Union’s Horizon 2020 research and innovation program under the SINE2020 project, grant agreement No. 654000.

## Author contributions

K.-M.N., A.M., and J.Y. conceptualized the project. A.M. and N.F. performed the VPS purification, DLS, and rheological characterizations. A.M. and J.Y. performed SAXS measurement. Y.Z. and M.Z. performed VPS quantification. A.M. and M.A. performed the biofilm growth and quantification. R.K. and R.J.W. performed all-atom MD simulations and associated analysis. K.-M.N. performed coarse-grained modeling and conformational sampling, under the guidance of Y. Li and J.Y. Y.-J.L., M.F.H., and M.-P.N. analyzed the SAXS data. E.G. and R.O. performed VPS digestion for SAXS analysis. J.Y., K.- M.N., A.M., and Y. Liu drafted the manuscript and all authors contributed to the final manuscript.

## Competing interests

The authors declare no competing interests.

## Supplementary Materials

Supplementary information is available for this paper. Correspondence and requests for materials should be addressed to.

---

**Fig. S1.**
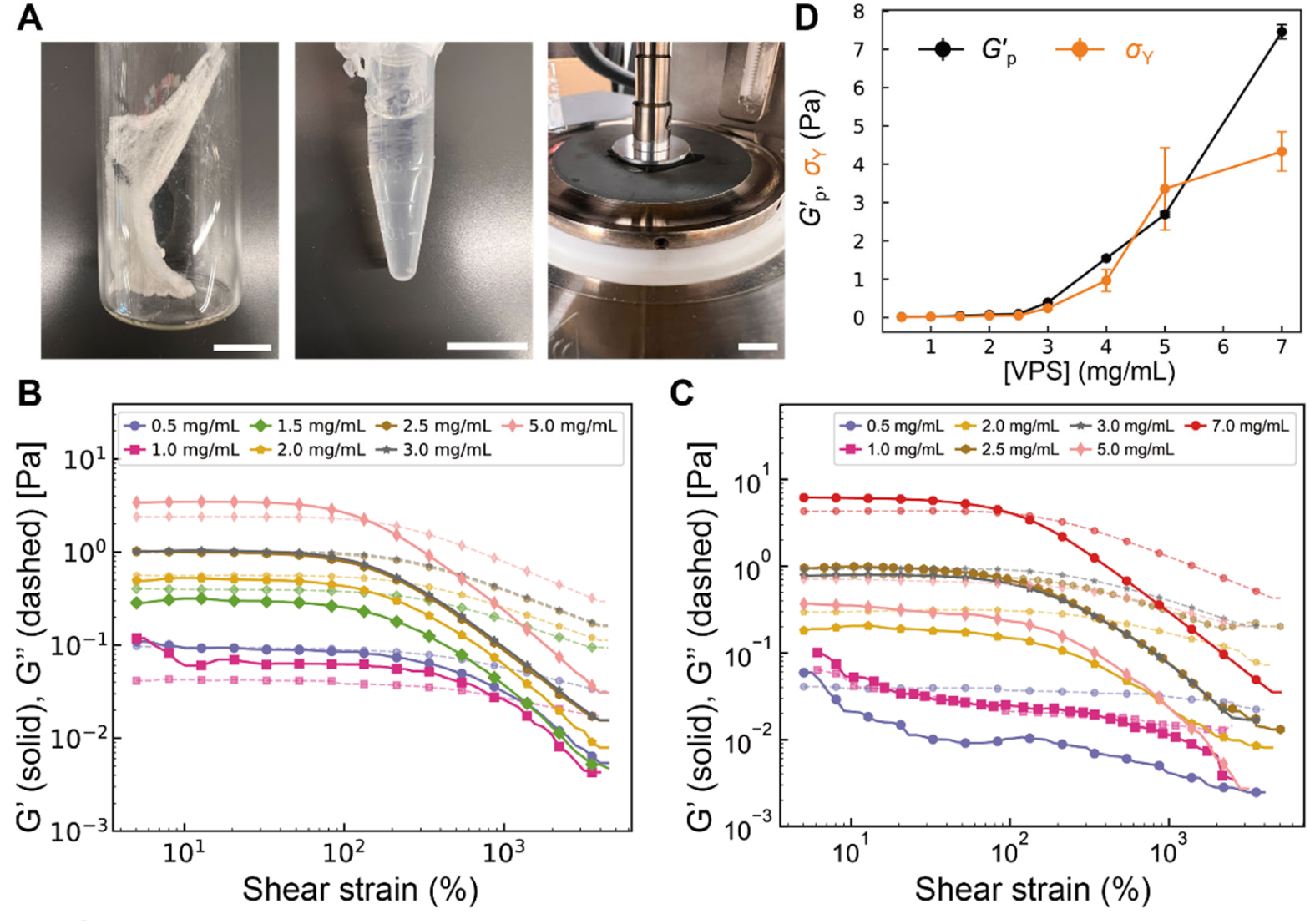
Rheological measurements of VPS solutions. (**A**) Representative images of purified VPS in solid state form (*top*), in solution (*middle*), and following transfer to the lower plate of a rheometer (*bottom*). Scale bars: 1 cm. (**B** and **C**) *G*′ and *G*′′ versus shear strain for VPS solutions in DI water (**B**) and PBS (**C**) at different concentrations. (**D**) Extracted plateau modulus *G*′_p_ and yield stress *σ*Y as a function of VPS concentration in PBS.

**Fig. S2.**
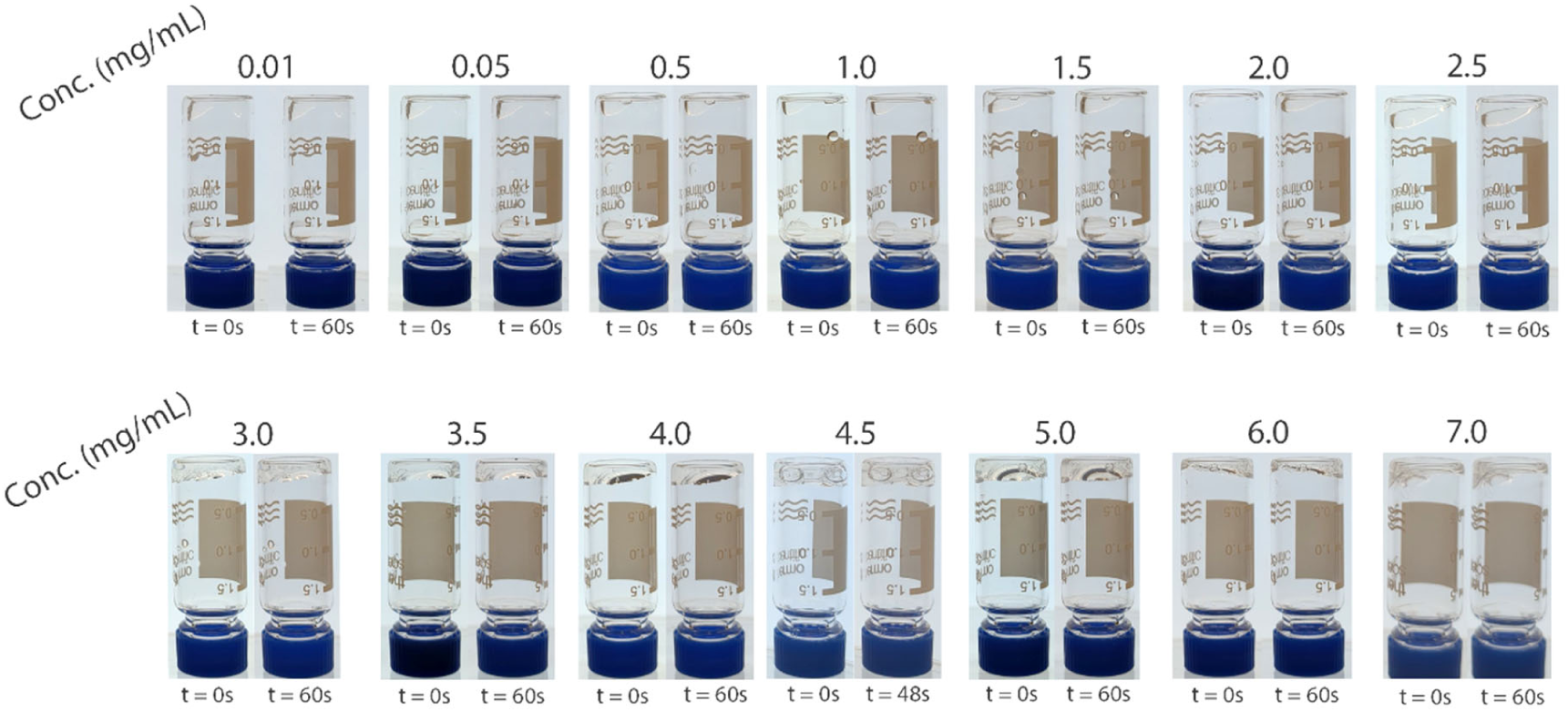
Pictures of VPS solutions in water at different concentrations, before and after flipping the vials containing them for 60. **s.** At or below 2 mg/mL, the solution quickly drips to the bottom. Above 2 mg/mL, the solution largely remains at the top.

**Fig. S3.**
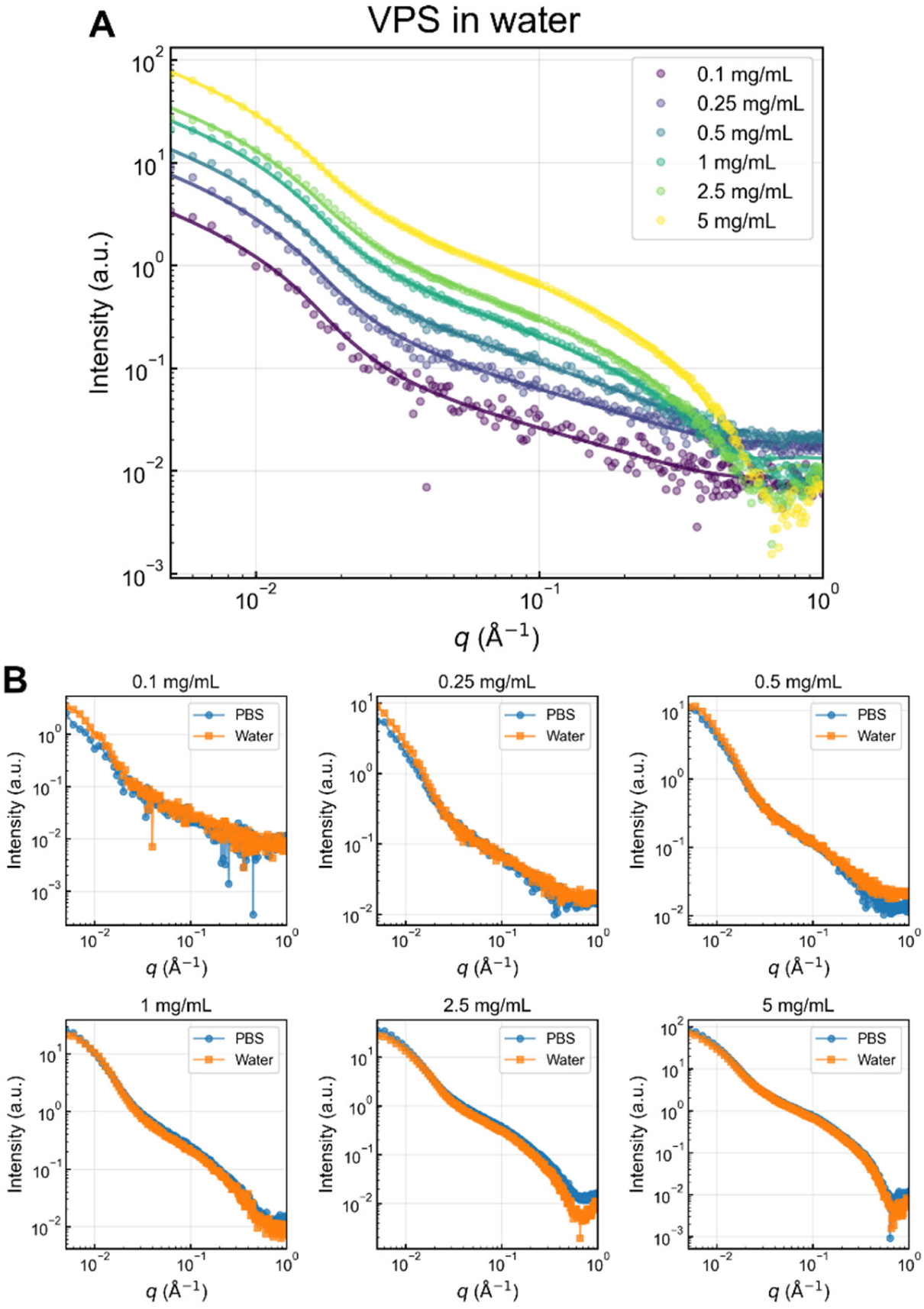
Additional SAXS data in DI water. (**A**) SAXS intensity versus *q* for different concentrations of VPS solutions in water. The same fitting procedure was applied to this dataset as in Fig. 3D. (**B**) Comparison between SAXS curves for VPS in water and in PBS at different concentrations.

**Fig. S4.**
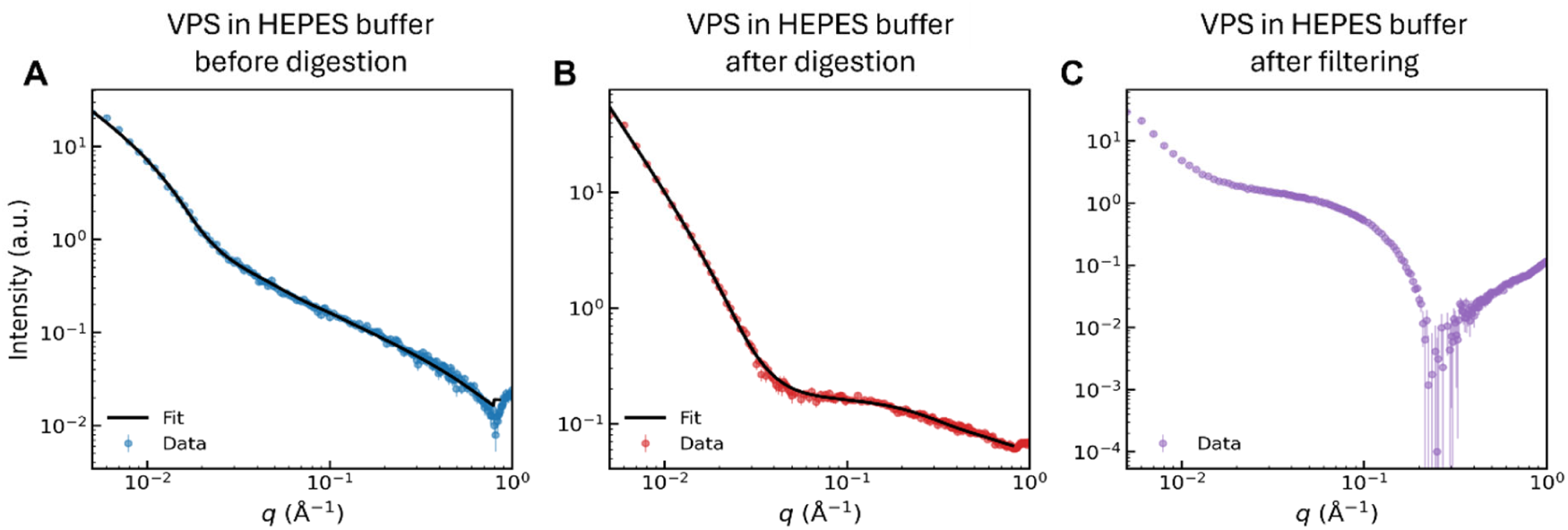
The observed features in SAXS curves arise from VPS polymers. (**A**) SAXS intensity curve for VPS solution in HEPES buffer, showing similar features to those in Fig. 3D. (**B**) SAXS curve for VPS solution after enzymatic digestion with RbmB in HEPES buffer into smaller units. The same fitting procedure was applied to the data in panels **A** and **B** as in Fig. 3D. (**C**) SAXS curve for VPS solution in HEPES buffer after filtering through a solid-phase extraction column to remove large polymers. No fitting can be appropriately obtained for this case. All initial concentrations of VPS solutions were ∼ 0.5 mg/mL.

**Fig. S5.**
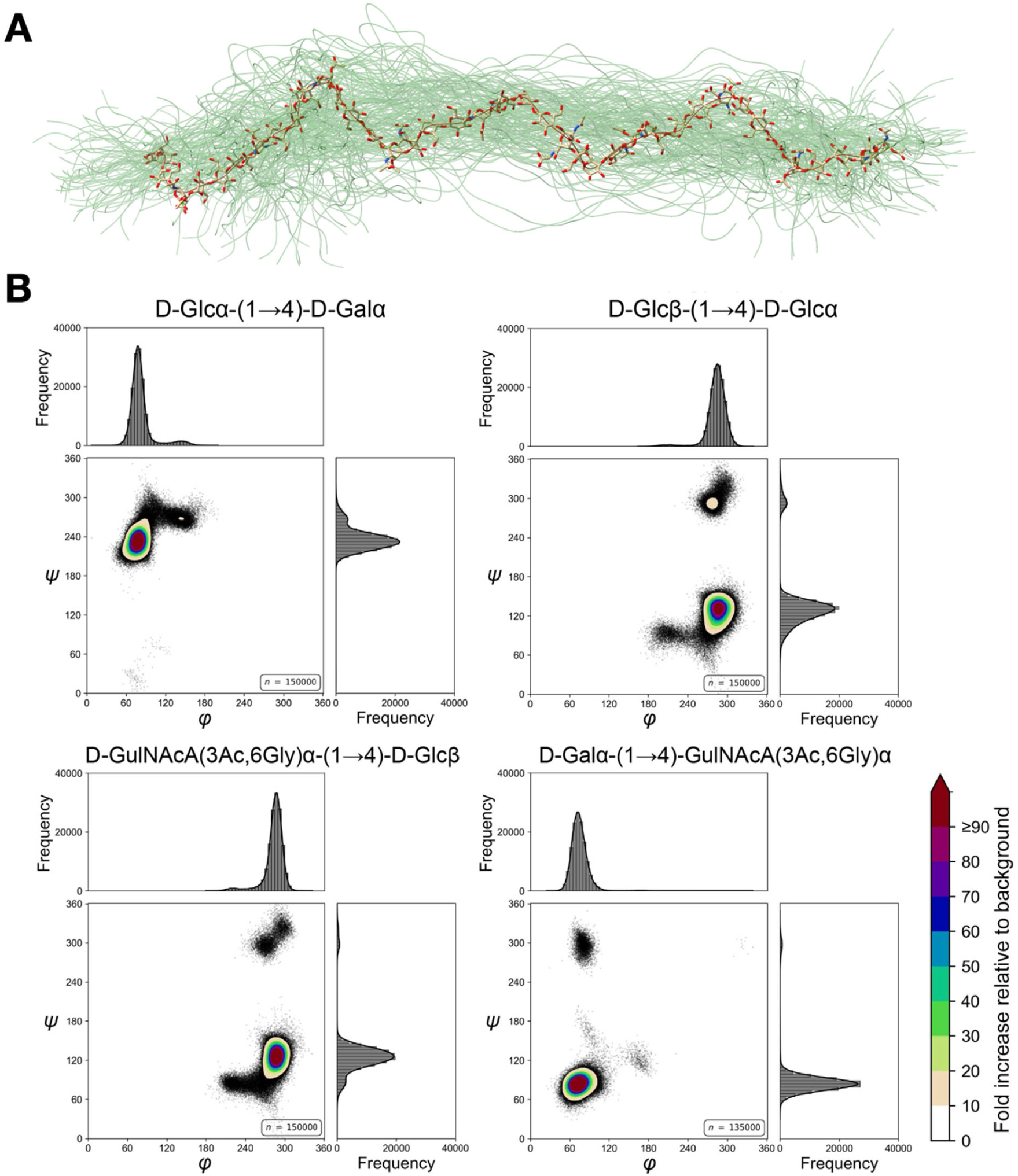
Dihedral angles for the four glycosidic linkage types. (**A**) Overlaid snapshots from the 10-tetrasaccharide VPS segment in the all-atom MD simulations, pooled over all three replicate trajectories. Each configuration is depicted as a spline interpolating the heavy-atom centroids of the monosaccharide rings along the segment. A representative configuration, namely the configuration with the least root-mean-square deviation to the average positions of the heavy atoms along the segment, is also shown. (**B**) Contour maps of the ф (O5–C1–O4’– C4’) and *ψ* (C1–O4’–C4’–C3’) dihedral angles along each of the four glycosidic linkage types in the VPS segment. For each linkage type, a histogram of dihedral angles was first computed on a 2D grid of 72 × 72 bins (5° resolution); the background frequency was then defined as the total number of points divided by the total number of bins. Each level set in the contour map represents the bins in the histogram that exceed the indicated fold-increase relative to this background frequency. See also Table S2.

**Fig. S6.**
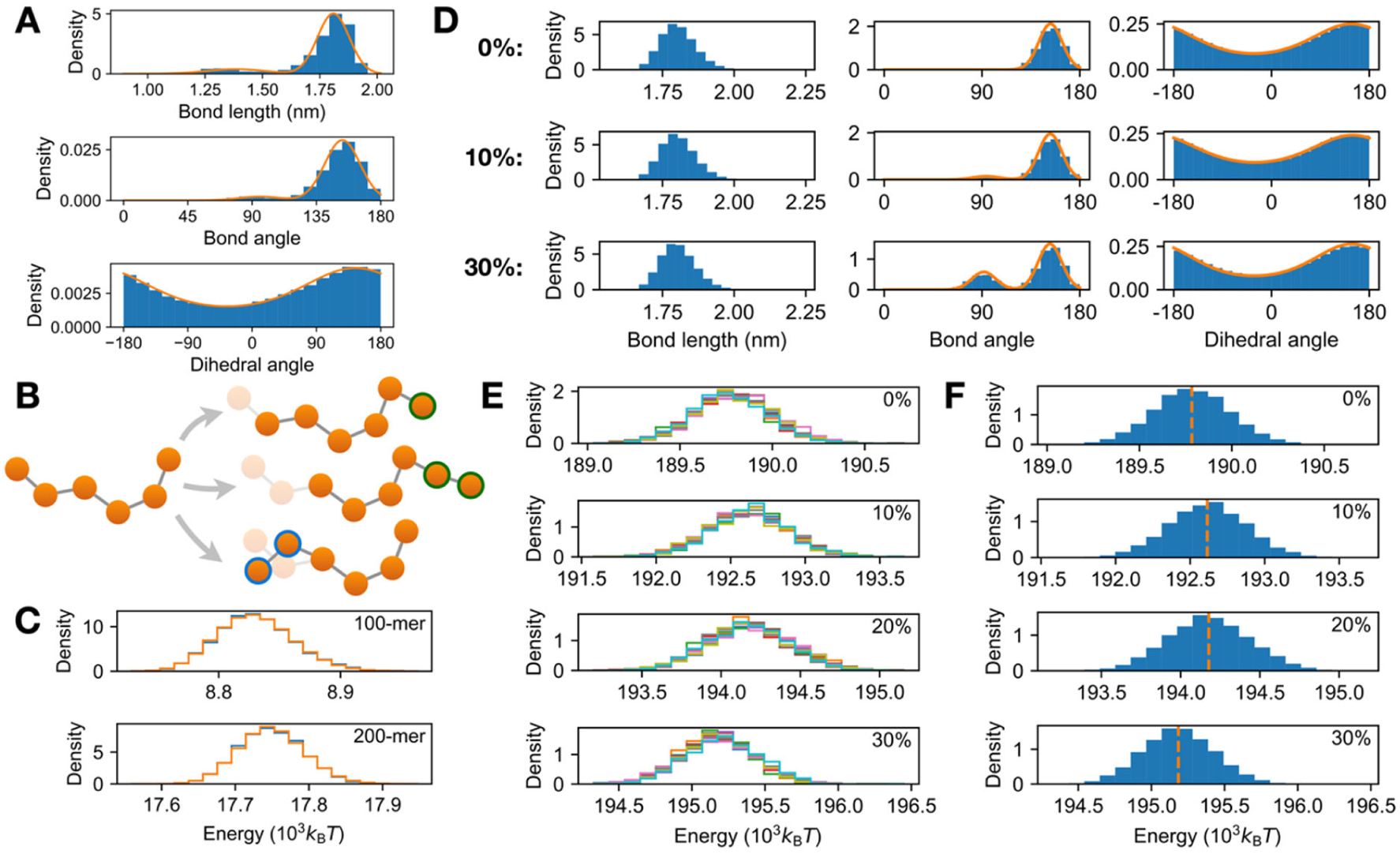
Coarse-grained VPS model and MC conformational sampling. (**A**) Empirical bond length, bond angle, and dihedral angle distributions along the coarse-grained VPS chain, obtained from each trajectory of the all-atom MD simulations. The bond length distribution was fitted to a two-component Gaussian mixture model; the bond angle distribution was fitted to a symmetrically folded two-component von Mises mixture model; and the dihedral angle distribution was fitted to a von Mises distribution (orange curves; see also Table S3). See Methods for details. (**B**) Schematic of the MC conformational sampling procedure. At each iteration, a set of new candidate polymer configurations was proposed using one of three move types: reptation by a single bead (*top*), reptation by multiple beads (*middle*), or terminal segment move (*bottom*). One candidate configuration was then selected and probabilistically accepted, as prescribed in configurational-bias MC^64^. Transparent beads indicate those in the original configuration that have been deleted or moved; beads with green outlines are new beads that have been introduced through reptation; and beads with blue outlines are pre-existing beads that have been moved. See Methods and Supplementary Information for details. (**C**) Distributions of configurational energy for a 100- or 200-bead random coil (*top* and *bottom*, respectively), obtained either from 10 independent MC runs (blue) or 20 independent MD trajectories (orange). The random coil was assumed to follow the same non-bonded interaction and bond length potentials as in the coarse-grained model for VPS (see Methods and Supplementary Information). All MD trajectories were run with a Langevin thermostat at 300 K for 500 ns each. (**D**) Bond length, bond angle, and dihedral angle distributions from 10 independent MC runs for the indicated kink fractions. Fitting each bond angle distribution to a folded two- component von Mises mixture model yields the desired fractions of linear (∼160°) and kinked (∼90°) bond angles; fitting each dihedral angle distribution to a von Mises distribution yields the desired mean dihedral angle (∼150°) and concentration (∼0.5). (**E** and **F**) Distributions of configurational energy from 10 independent MC runs for each kink fraction, shown as separate distributions for each run in panel **E** and as one distribution pooled over all 10 runs in panel **F**. The mean of each distribution in panel **F** is shown in orange. The overall similarity of each group of 10 distributions in panel **E** to each other, and to the corresponding pooled distribution in panel **F**, suggest that the procedure is effectively sampling from a common equilibrium distribution.

**Fig. S7.**
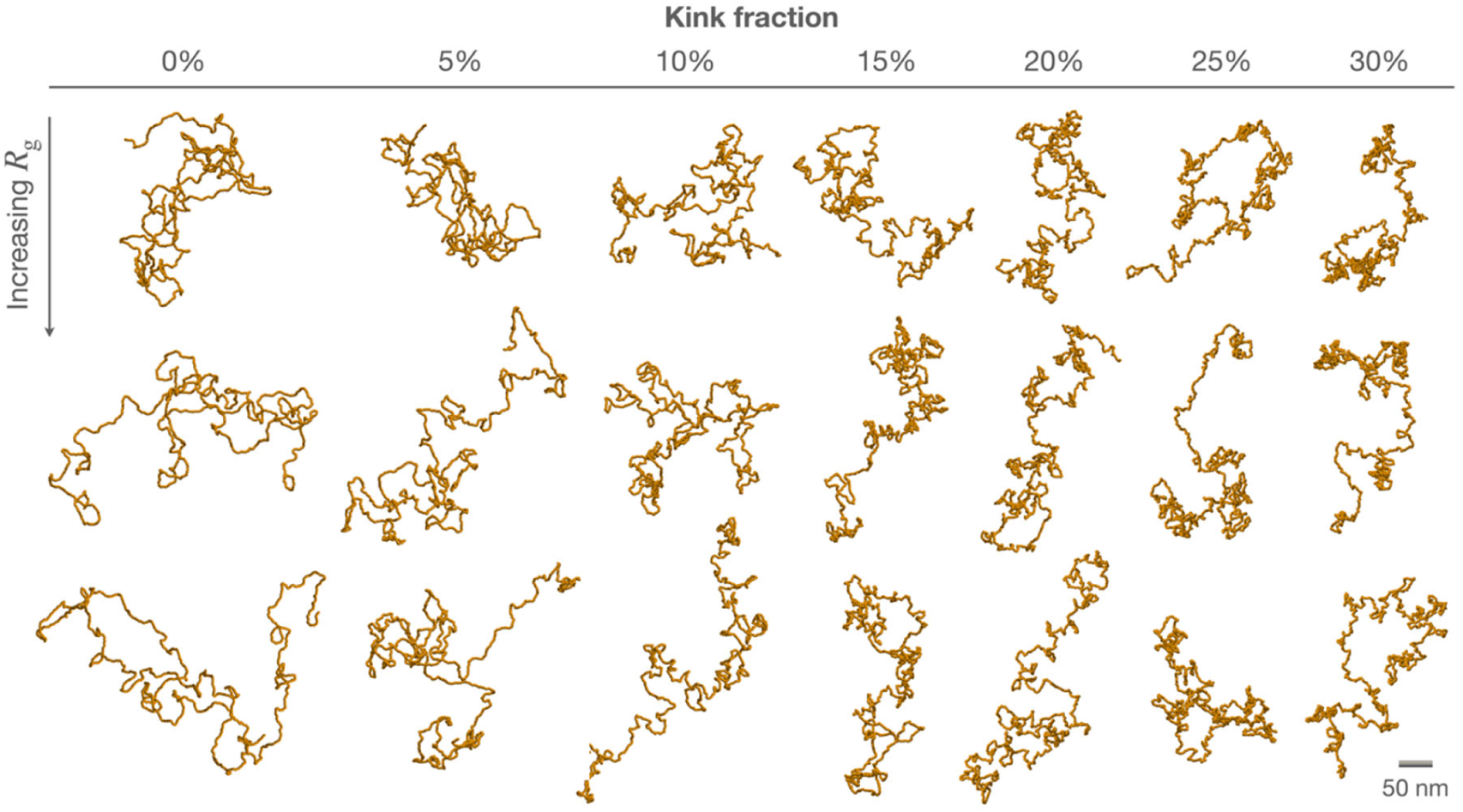
Example coarse-grained polymer configurations. For each kink fraction, the three configurations with radii of gyration closest to the first, second, and third quartiles (top, middle, and bottom rows, respectively) over the corresponding ensemble are shown. Each set of quartiles was quantified over a pooled ensemble obtained from 10 independent MC runs.

**Fig. S8.**
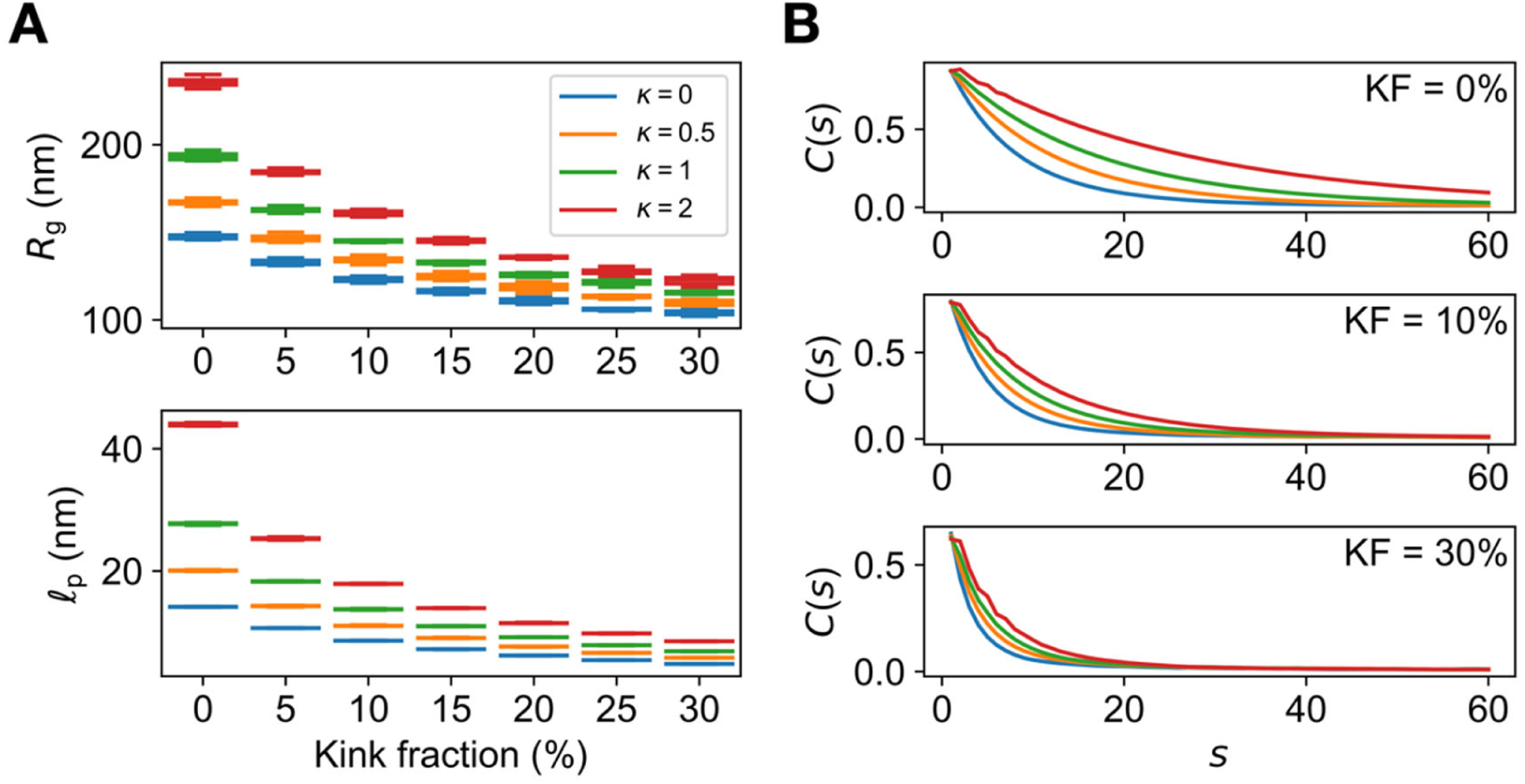
Effects of dihedral angle stiffness on the radius of gyration and persistence length. (**A**) Root-mean-square radius of gyration (*R*_g_) and persistence length (*l*_p_) as a function of the kink fraction, for different choices of dihedral angle stiffness, n = *K*_dihedral_/(*k*_B_*T*). n = 0.5 corresponds to the coarse-grained model for VPS (Fig. 4F; see also Table S4). Each distribution was obtained from 10 measurements, each taken from an independent MC run for a coarse- grained VPS polymer consisting of 2,100 units. Whiskers correspond to 1.5 times the interquartile range below and above the first and third quartiles, respectively. (**B**) Tangent vector autocorrelation function, *C*(*s*), along coarse-grained VPS polymers with the indicated kink fractions and the same values of n as in panel **A**. The autocorrelation for a fixed value of *s* was calculated as the average dot product over all pairs of unit tangent vectors *s* bonds apart along the polymer, averaged over all configurations in the ensemble (see Supplementary Information). Each curve corresponds to an ensemble of configurations obtained from one MC run. We note that, while the dihedral potential imposes some helicity on the VPS polymer (see Methods), a stiffness of *K*_dihedral_ ≤ *k*_B_*T* is sufficiently weak that it merely increases the persistence length of the polymer, without imparting any noticeable periodicity to the autocorrelation function; some weak periodicity is apparent at *K*_dihedral_ = 2*k*_B_*T* , but the autocorrelation function still appears to be well-described by an exponential decay.

**Fig. S9.**
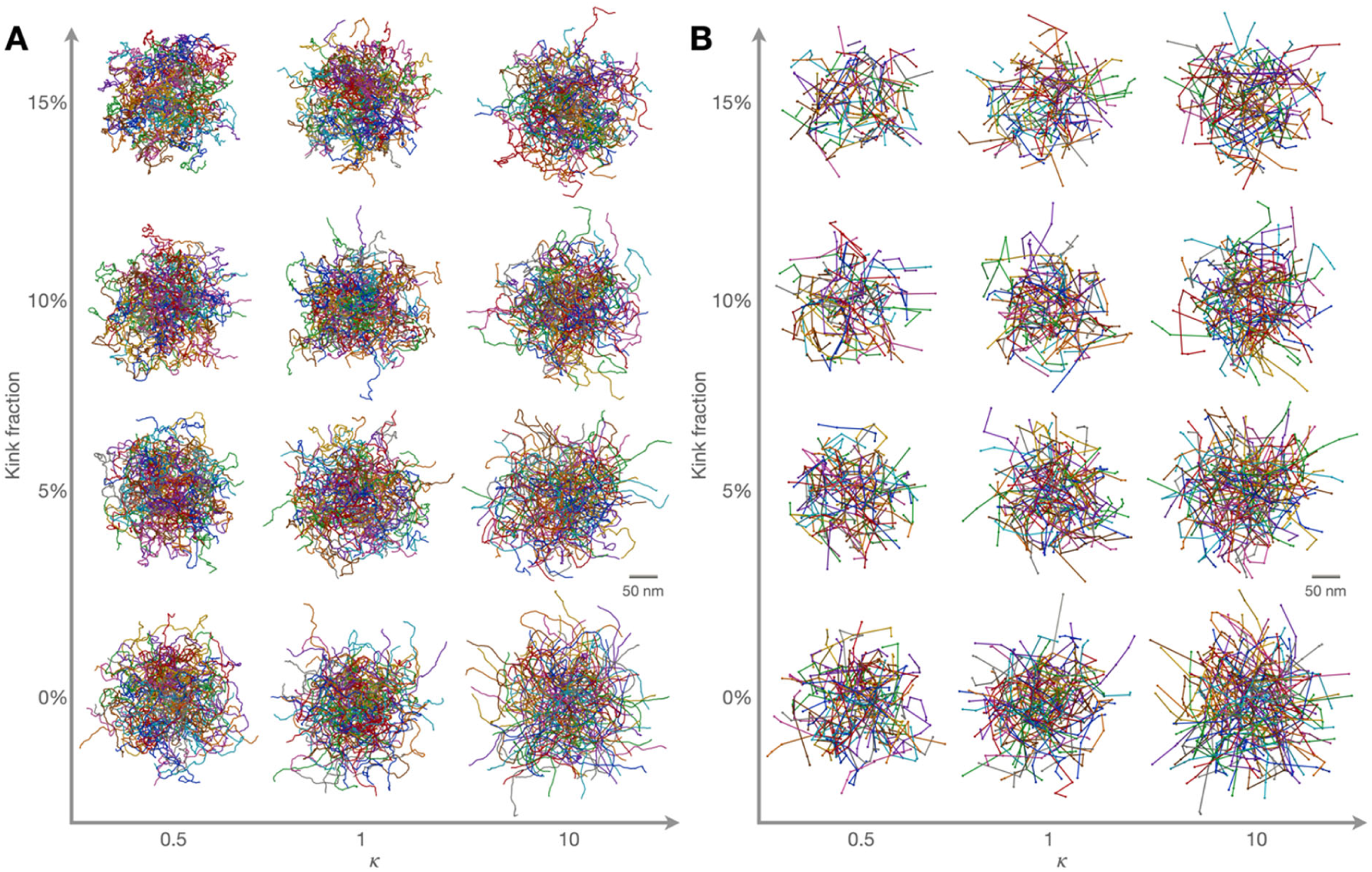
Primitive path analysis of coarse-grained polymer solution configurations. Representative configurations of solutions of VPS-like chains with the indicated values of kink fraction and dihedral stiffness, n = *K*_dihedral_/(*k*_B_*T*) (**A**), and the corresponding collections of primitive paths (**B**), each obtained using the Z1+ package. Each chain in panel **A** and each primitive path in panel **B** is plotted such that its center-of-mass lies in the fundamental unit cell in the corresponding periodic domain; as such, a chain and its corresponding primitive path may occasionally be visualized in different positions.

**Fig. S10.**
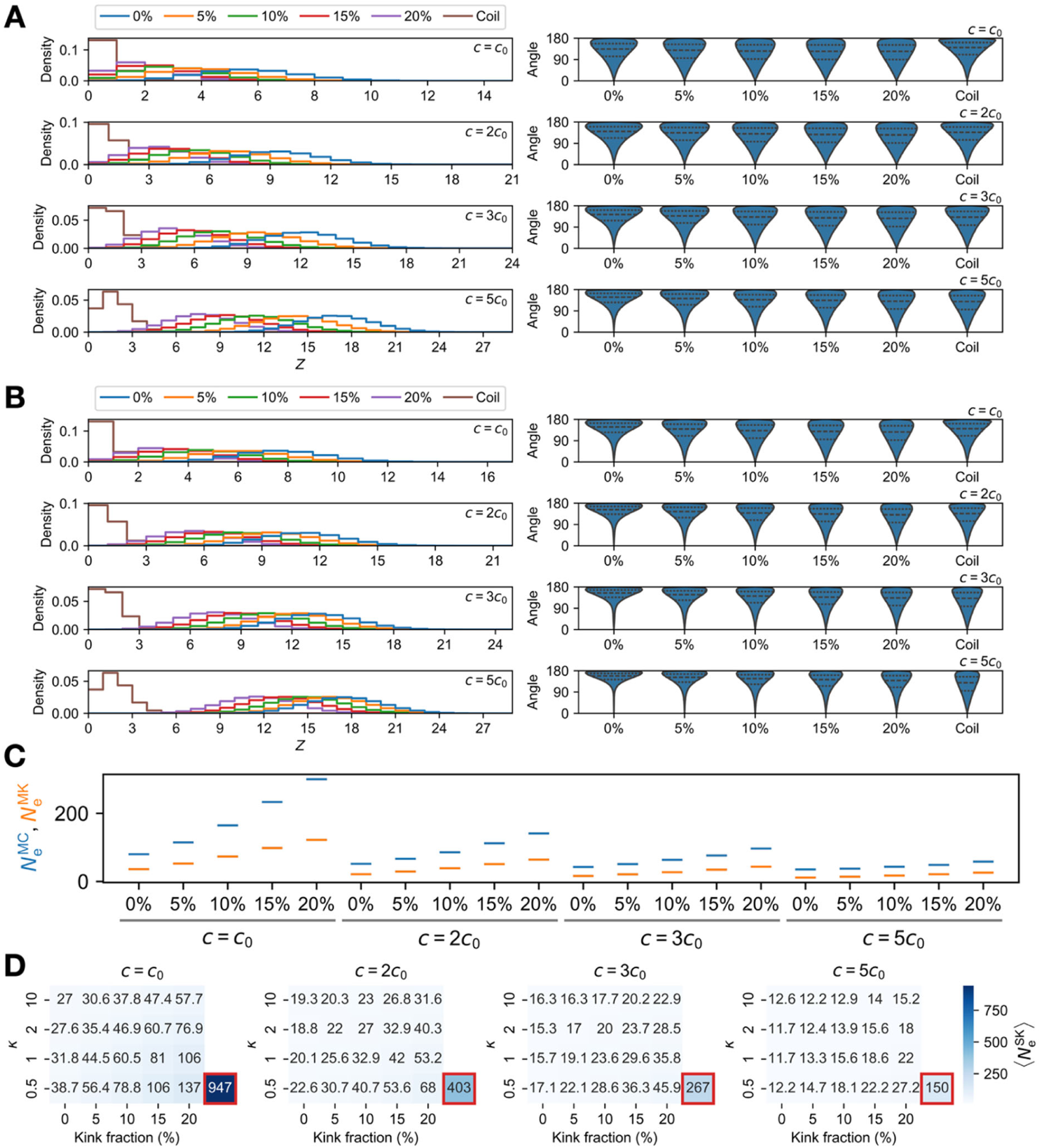
Entanglement-related quantities for different kink fractions and concentrations. (**A** and **B**) Distributions of the number, *Z*, of internal nodes along each primitive path (*left*) and the angles formed at these internal nodes (*right*) for each choice of kink fraction and concentration (*c*), for dihedral stiffnesses of *K*_dihedral_ = 0.5*k*_B_*T* (**A**) and *K*_dihedral_ = 2*k*_B_*T* (**B**). Analogous distributions for solutions of random coils following the same non-bonded interaction and bond length potentials are shown for comparison. (**C**) The “M-coil” and “M- kink” estimators, 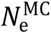 and 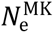, for the entanglement length, *N*, for *c* = *c*_0_, 2*c*_0_, 3*c*_0_, 5*c*_0_ and *K*_dihedral_ = 0.5*k*_B_*T* . We note that 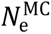 is around twice 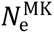 for all kink fractions and concentrations, which is consistent with previous work^110^; 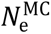 better corresponds to the “rheological” entanglement length that determines the plateau modulus. (**D**) The “modified S-kink” estimator, 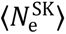, for *c* = *c*_0_, 2*c*_0_, 3*c*_0_, 5*c*_0_ and 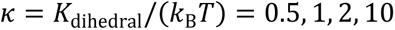. This estimator is simpler to evaluate than 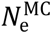 and 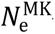—which require configurations of polymer solutions with different choices of polymer length, *N*—but is less statistically robust.

**Table S1.** List of *V. cholerae* strains used in this paper.

| Strain | Genotype | Source |
| --- | --- | --- |
| JY286 | *vpvC*<sup>W240R</sup>Δ*rbmA*Δ*bap1*Δ*rbmC*Δ*pomA* | 48 |
| JY462 | *vpvC*<sup>W240R</sup>Δ*rbmA*Δ*bap1*Δ*rbmC*Δ*VC1807::P<sub>tac</sub>-mNeonGreen*, Spec<sup>R</sup> | 43 |
| JY477 | *vpvC*<sup>W240R</sup>Δ*rbmA*Δ*bap1*Δ*rbmC*Δ*vpsL*Δ*VC1807::P<sub>tac</sub>-mNeonGreen*, Spec<sup>R</sup> | 43 |

**Table S2.**
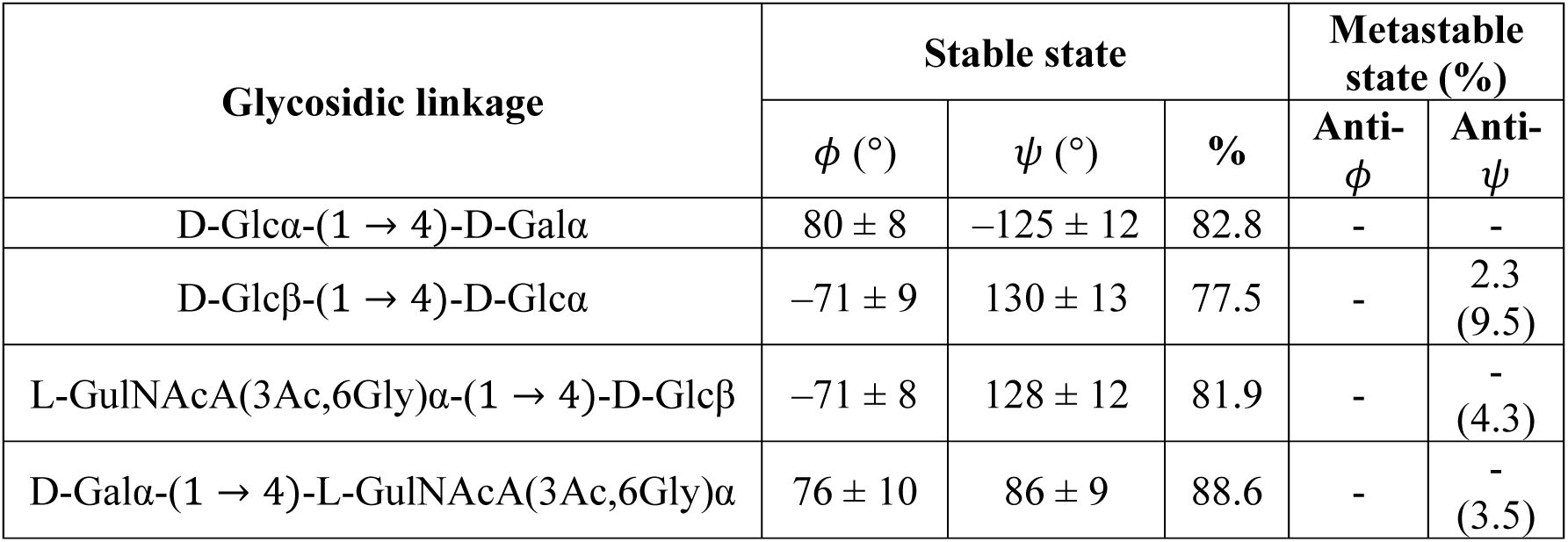
Conformational statistics of the four glycosidic linkage types. States and their empirical probabilities were identified according to the contour maps in Fig. S5B; in particular, the empirical probabilities correspond to the fraction of points in (ф, *ψ*)-space that lie within the level sets corresponding to a greater than 10-fold increase over the background frequency. We also quantified the percentage of points near the level set corresponding to the D-Glcβ-(1 → 4)-D-Glcα anti-*ψ* conformation (∼ 7.2%, yielding a total percentage of ∼ 9.5%), as well as the percentages of points that may represent analogous anti- *ψ* conformations for the L- GulNAcA(3Ac,6Gly)α- (1 → 4) -D-Glcβ and D-Galα- (1 → 4) -L-GulNAcA(3Ac,6Gly)α linkages, but did not meet the 10-fold threshold.

| Glycosidic linkage | Stable state |  |  | Metastable state (%) |  |
| --- | --- | --- | --- | --- | --- |
| | $\phi$ (°) | $\psi$ (°) | % | Anti-ϕ | Anti-ψ |
| D-Glcα-(1 → 4)-D-Galα | 80 ± 8 | -125 ± 12 | 82.8 | - | - |
| D-Glcβ-(1 → 4)-D-Glcα | -71 ± 9 | 130 ± 13 | 77.5 | - | 2.3<br>(9.5) |
| L-GulNAcA(3Ac,6Gly)α-(1 → 4)-D-Glcβ | -71 ± 8 | 128 ± 12 | 81.9 | - | -<br>(4.3) |
| D-Galα-(1 → 4)-L-GulNAcA(3Ac,6Gly)$\alpha$ | 76 ± 10 | 86 ± 9 | 88.6 | - | -<br>(3.5) |

**Table S3.** Bond length, bond angle, and dihedral angle statistics for the coarse-grained VPS chain. Only the major component of the bond length mixture model is specified.

| **Bond lengths (major component only, weight = 87.1%)** |  |  |
| --- | --- | --- |
| Mean |  | 1.81 nm |
| Standard deviation |  | 0.0690 nm |
| **Bond angles** |  |  |
| Peak 1 | Mean | 153° |
|  | Concentration | 21.5 |
|  | Weight | 92.7% |
| Peak 2 | Mean | 95.2° |
|  | Concentration | 15.0 |
|  | Weight | 7.3% |
| **Dihedral angles** |  |  |
| Mean |  | 144° |
| Concentration |  | 0.523 |

**Table S4.** Parameters for MC conformational sampling of coarse-grained VPS chains. See Supplementary Information for a full discussion of the parameters and their definitions. To set the Lennard-Jones and FENE potential parameters (*σ*, *k*_FENE_ , and *R*_0_ ), we first fixed *σ* = (*l*_0_⟩/0.965 ≈ 1.86*η* nm and *R*_0_ = 1.5*σ* ≈ 2.798 nm, as in Svaneborg and Everaers’ modified Kremer–Grest model^62,63^, and set *k*_FENE_ so as to obtain a Boltzmann distribution with mean bond length (*l*_0_⟩ = 1.8 nm. This yielded an optimal value of *k*_FENE_/(*k*_B_*T*) ≈ 9.432 nm^−2^, and so we set *k*_FENE_/(*k*_B_*T*) = 9 nm^−2^. To set *w*_1_ and *w*_2_, we recalled that a von Mises distribution with a large concentration parameter, n » 1, is approximately Gaussian with variance *σ*^2^ = 1/n. As such, using the concentration parameter of ∼ 20 for the major component in the bond angle mixture in Table S3, we set 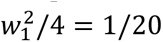, or 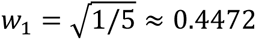; for simplicity, we set *w*_2_ = *w*_1_. Finally, we set *K*_dihedral_ = 0.5*k*_B_*T* to match the concentration parameter of the von Mises fit in Table S3, and set ð = —30° to accommodate an equilibrium dihedral angle of 150°.

| Parameter | Meaning | Value |
| --- | --- | --- |
| $T$ | Temperature | 300 K |
| $\langle l_0 \rangle$ | Mean bond length | 1.8 nm |
| $\sigma$ | Non-bonded interaction length scale | $\langle l_0 \rangle / 0.965 \approx 1.865$ nm |
| $k_{\text{FENE}}$ | Bond stiffness | $(9 \text{ nm}^{-2}) \cdot k_{\text{B}}T$ |
| $R_0$ | Maximum bond length | $1.5\sigma \approx 2.798$ nm |
| $w_1$ | Two times the standard deviation of the larger bond angles in the bimodal angle potential | $\sqrt{1/5} \approx 0.4472$ |
| $w_2$ | Two times the standard deviation of the smaller bond angles in the bimodal angle potential | $\sqrt{1/5} \approx 0.4472$ |
| $K_{\text{dihedral}}$ | Dihedral angle stiffness | $0.5k_{\text{B}}T$ |
| $\delta$ | Dihedral angle offset | $-30^\circ$ |

## Notes

https://github.com/kmnam/biofilm-matrix-crosslinking

### Competing Interest Statement

The authors have declared no competing interest.

## REFERENCES

1. George, A., Sanjay, M. R., Srisuk, R., Parameswaranpillai, J. & Siengchin, S. A comprehensive review on chemical properties and applications of biopolymers and their composites. Int. J. Biol. Macromol. 154, 329–338 (2020).

2. Varki, A. et al. Glycomics. in Essentials of Glycobiology (Cold Spring Harbor Laboratory Press, Cold Spring Harbor (NY), 2009).

3. Wanke, A., Malisic, M., Wawra, S. & Zuccaro, A. Unraveling the sugar code: the role of microbial extracellular glycans in plant–microbe interactions. JXB 72, 15–35 (2021).

4. Calles-Garcia, D. & Dube, D. H. Chemical biology tools to probe bacterial glycans. Curr. Opin. Chem. Biol. 80, 102453 (2024).

5. Drickamer, K. & Taylor, M. E. Glycan arrays for functional glycomics. Genome Biol. 3, REVIEWS1034 (2002).

6. Geissner, A., Anish, C. & Seeberger, P. H. Glycan arrays as tools for infectious disease research. Curr. Opin. Chem. Biol. 18, 38–45 (2014).

7. Flemming, H.-C. & Wingender, J. The biofilm matrix. Nat. Rev. Microbiol. 8, 623–633 (2010).

8. Dragoš, A. & Kovács, Á. T. The peculiar functions of the bacterial extracellular matrix. Trends Microbiol. 25, 257–266 (2017).

9. Flemming, H.-C., Neu, T. R. & Wozniak, D. J. The EPS matrix: the ‘house of biofilm cells’. J. Bacteriol. 189, 7945–7947 (2007).

10. Hobley, L., Harkins, C., Macphee, C. E. & Stanley-Wall, N. R. Giving structure to the biofilm matrix: an overview of individual strategies and emerging common themes. FEMS Microbiol. Rev. 39, 649–669 (2015).

11. Billings, N., Birjiniuk, A., Samad, T. S., Doyle, P. S. & Ribbeck, K. Material properties of biofilms — a review of methods for understanding permeability and mechanics. Rep. Prog. Phys. 78, 036601 (2015).

12. Gordon, V. D., Davis-Fields, M., Kovach, K. & Rodesney, C. A. Biofilms and mechanics: a review of experimental techniques and findings. J. Phys. D 50, 223002 (2017).

13. Yan, J. & Bassler, B. L. Surviving as a community: antibiotic tolerance and persistence in bacterial biofilms. Cell Host Microbe 26, 15–21 (2019).

14. Qian, P.-Y., Cheng, A., Wang, R. & Zhang, R. Marine biofilms: diversity, interactions and biofouling. Nat. Rev. Microbiol. 20, 671–684 (2022).

15. Nelson, E. J., Harris, J. B., Morris, J. G., Calderwood, S. B. & Camilli, A. Cholera transmission: the host, pathogen and bacteriophage dynamic. Nat. Rev. Microbiol. 7, 693–702 (2009).

16. Yildiz, F., Fong, J., Sadovskaya, I., Grard, T. & Vinogradov, E. Structural characterization of the extracellular polysaccharide from *Vibrio cholerae* O1 El-Tor. PLoS ONE 9, e86751 (2014).

17. Fong, J. C. N., Syed, K. A., Klose, K. E. & Yildiz, F. H. Role of *Vibrio* polysaccharide (vps) genes in VPS production, biofilm formation and *Vibrio cholerae* pathogenesis. Microbiology 156, 2757–2769 (2010).

18. Teschler, J. K. et al. Living in the matrix: assembly and control of *Vibrio cholerae* biofilms. Nat. Rev. Microbiol. 13, 255–268 (2015).

19. Teschler, J. K., Nadell, C. D., Drescher, K. & Yildiz, F. H. Mechanisms underlying *Vibrio cholerae* biofilm formation and dispersion. Annu. Rev. Microbiol. 76, 503–532 (2022).

20. Berk, V. et al. Molecular architecture and assembly principles of *Vibrio cholerae* biofilms. Science 337, 236–239 (2012).

21. Fong, J. C. et al. Structural dynamics of RbmA governs plasticity of *Vibrio cholerae* biofilms. eLife 6, e1002210 (2017).

22. Fong, J. C. N., Karplus, K., Schoolnik, G. K. & Yildiz, F. H. Identification and characterization of RbmA, a novel protein required for the development of rugose colony morphology and biofilm structure in *Vibrio cholerae*. J. Bacteriol. 188, 1049–1059 (2006).

23. Fong, J. C. N. & Yildiz, F. H. The *rbmBCDEF* gene cluster modulates development of rugose colony morphology and biofilm formation in *Vibrio cholerae*. J. Bacteriol. 189, 2319–2330 (2007).

24. Absalon, C., Van Dellen, K. & Watnick, P. I. A communal bacterial adhesin anchors biofilm and bystander cells to surfaces. PLOS Pathog. 7, e1002210 (2011).

25. Huang, X., et al. *Vibrio cholerae* biofilms use modular adhesins with glycan-targeting and nonspecific surface binding domains for colonization. Nat. Commun. 14, 2104 (2023).

26. Moreau, A. et al. Surface remodeling and inversion of cell-matrix interactions underlie community recognition and dispersal in *Vibrio cholerae* biofilms. Nat. Commun. 16, 327 (2025).

27. Weerasekera, R., et al. *Vibrio cholerae* RbmB is an α-1,4-polysaccharide lyase with biofilm-disrupting activity against Vibrio Polysaccharide (VPS). PLOS Pathog. 20, e1012750 (2024).

28. Weerasekera, R. et al. Crystal structure of *Vibrio cholerae* polysaccharide lyase RbmB bound to *Vibrio* polysaccharide (VPS) fragments provides insights into substrate recognition and cleavage. Proc. Natl. Acad. Sci. USA 123, e2534280123 (2026).

29. Moon, H., Yamani, S., Lee, J. & McKinley, G. H. Tuning the shear and extensional rheology of semi-flexible polyelectrolyte solutions. arXiv:2410.15132v1 (2024) doi:arXiv:2410.15132v1.

30. Hossain, M. T. & Ewoldt, R. H. Protorheology. J. Rheol. 68, 113–144 (2024).

31. Hossain, M. T., Tiwari, R. & Ewoldt, R. H. Protorheology in practice: Avoiding misinterpretation. Curr. Opin. Colloid Interface Sci. 74, 101866 (2024).

32. Lopez, C. G., Colby, R. H., Graham, P. & Cabral, J. T. Viscosity and scaling of semiflexible polyelectrolyte NaCMC in aqueous salt solutions. Macromolecules 50, 332–338 (2017).

33. Rubinstein, M. & Colby, R. H. Polymer Physics. vol. 23 (Oxford university press New York, 2003).

34. Ganesan, M. et al. Molar mass, entanglement, and associations of the biofilm polysaccharide of *Staphylococcus epidermidis*. Biomacromolecules 14, 1474–1481 (2013).

35. Rubinstein, M. & Semenov, A. N. Thermoreversible gelation in solutions of associating polymers. 2. Linear dynamics. Macromolecules 31, 1386–1397 (1998).

36. Heo, Y. & Larson, R. G. The scaling of zero-shear viscosities of semidilute polymer solutions with concentration. J. Rheol. 49, 1117–1128 (2005).

37. Rubinstein, M. & Semenov, A. N. Dynamics of entangled solutions of associating polymers. Macromolecules 34, 1058–1068 (2001).

38. Lang, P. & Frey, E. Disentangling entanglements in biopolymer solutions. Nat. Commun. 9, 494 (2018).

39. Romo-Uribe, A. Shear rheology and scaling of semiflexible polymers: Effect of polymer- solvent interactions in the semidilute regime. J. Appl. Polym. Sci. 138, 49712 (2021).

40. Semenov, A. N. Dynamics of concentrated solutions of rigid-chain polymers. Part 1.— Brownian motion of persistent macromolecules in isotropic solution. J. Chem. Soc., Faraday Trans. 2 82, 317–329 (1986).

41. Odijk, T. The statistics and dynamics of confined or entangled stiff polymers. Macromolecules 16, 1340–1344 (1983).

42. Ganesan, M., Knier, S., Younger, J. G. & Solomon, M. J. Associative and entanglement contributions to the solution rheology of a bacterial polysaccharide. Macromolecules 49, 8313–8321 (2016).

43. Nijjer, J. et al. Biofilms as self-shaping growing nematics. Nat. Phys. 19, 1936–1944 (2023).

44. Phillips, R. & Milo, R. A feeling for the numbers in biology. Proc. Natl. Acad. Sci. USA 106, 21465–21471 (2009).

45. Smith, R. L. & Gilkerson, E. Quantitation of glycosaminoglycan hexosamine using 3- methyl-2-benzothiazolone hydrazone hydrochloride. Anal. Biochem. 98, 478–480 (1979).

46. Sanz-Jiménez, A. et al. High-throughput determination of dry mass of single bacterial cells by ultrathin membrane resonators. *Commun*. Biol. 5, 1227 (2022).

47. Yan, J., Nadell, C. D. & Bassler, B. L. Environmental fluctuation governs selection for plasticity in biofilm production. ISME J. 11, 1569–1577 (2017).

48. Yan, J. et al. Bacterial biofilm material properties enable removal and transfer by capillary peeling. Adv. Mater. 30, e1804153 (2018).

49. Zhang, Q. et al. Mechanical resilience of biofilms toward environmental perturbations mediated by extracellular matrix. Adv. Funct. Mater. 32, 2110699 (2022).

50. Yan, J., Sharo, A. G., Stone, H. A., Wingreen, N. S. & Bassler, B. L. *Vibrio cholerae* biofilm growth program and architecture revealed by single-cell live imaging. Proc. Natl. Acad. Sci. USA 113, E5337–E5343 (2016).

51. Stephens, Z., Wilson, L. F. L. & Zimmer, J. Diverse mechanisms of polysaccharide biosynthesis, assembly and secretion across kingdoms. Curr. Opin. Struct. Biol. 79, 102564 (2023).

52. Schwechheimer, C. et al. A tyrosine phosphoregulatory system controls exopolysaccharide biosynthesis and biofilm formation in *Vibrio cholerae*. PLOS Pathog. 16, e1008745 (2020).

53. Tianjiao, M. et al. Structural characterization and synergistic mechanism of *Bacillus thuringiensis* G033A exopolysaccharides enhancing insecticidal protein efficacy. Int. J. Biol. Macromol. 358, 151710 (2026).

54. Wang, W., Ju, Y., Liu, N., Shi, S. & Hao, L. Structural characteristics of microbial exopolysaccharides in association with their biological activities: a review. Chem. Biol. Technol. Agric. 10, 137 (2023).

55. Sutthasupa, S., Shiotsuki, M. & Sanda, F. Recent advances in ring-opening metathesis polymerization, and application to synthesis of functional materials. Polym. J. 42, 905–915 (2010).

56. Matyjaszewski, K. Controlled Radical Polymerization: State-of-the-Art in 2014. in Controlled Radical Polymerization: Mechanisms vol. 1187 1–17 (American Chemical Society, 2015).

57. Hammouda, B. A new Guinier-Porod model. J. Appl. Crystallogr. 43, 716–719 (2010).

58. Hammouda, B. Analysis of the Beaucage model. J. Appl. Crystallogr. 43, 1474–1478 (2010).

59. Pedersen, J. S. Analysis of small-angle scattering data from colloids and polymer solutions: modeling and least-squares fitting. Adv. Colloid Interface Sci. 70, 171–210 (1997).

60. Case, D. A. et al. AmberTools. J. Chem. Inf. Model. 63, 6183–6191 (2023).

61. Kirschner, K. N. et al. GLYCAM06: A generalizable biomolecular force field. Carbohydrates. J. Comput. Chem. 29, 622–655 (2008).

62. Everaers, R., Karimi-Varzaneh, H. A., Fleck, F., Hojdis, N. & Svaneborg, C. Kremer– Grest models for commodity polymer melts: linking theory, experiment, and simulation at the Kuhn scale. Macromolecules 53, 1901–1916 (2020).

63. Svaneborg, C. & Everaers, R. Characteristic time and length scales in melts of Kremer– Grest bead–spring polymers with wormlike bending stiffness. Macromolecules 53, 1917–1941 (2020).

64. Siepmann, J. I. & Frenkel, D. Configurational bias Monte Carlo: a new sampling scheme for flexible chains. Mol. Phys. 75, 59–70 (1992).

65. Kremer, K. & Grest, G. S. Dynamics of entangled linear polymer melts: A molecular- dynamics simulation. J. Chem. Phys. 92, 5057–5086 (1990).

66. Milano, G., Goudeau, S. & Müller-Plathe, F. Multicentered Gaussian-based potentials for coarse-grained polymer simulations: Linking atomistic and mesoscopic scales. J. Polym. Sci. B 43, 871–885 (2005).

67. Kröger, M., Dietz, J. D., Hoy, R. S. & Luap, C. The Z1+ package: Shortest multiple disconnected path for the analysis of entanglements in macromolecular systems. Comput. Phys. Commun. 283, 108567 (2023).

68. Everaers, R. et al. Rheology and microscopic topology of entangled polymeric liquids. Science 303, 823–826 (2004).

69. Doi, M. & Edwards, S. F. The Theory of Polymer Dynamics. (Clarendon Press, Oxford, 2013).

70. Morse, D. C. Tube diameter in tightly entangled solutions of semiflexible polymers. *Phys*. Rev. E 63, 031502 (2001).

71. Hoy, R. S., Foteinopoulou, K. & Kröger, M. Topological analysis of polymeric melts: Chain-length effects and fast-converging estimators for entanglement length. *Phys*. Rev. E 80, 031803 (2009).

72. Dietz, J. D., Kröger, M. & Hoy, R. S. Validation and refinement of unified analytic model for flexible and semiflexible polymer melt entanglement. Macromolecules 55, 3613–3626 (2022).

73. Antoniou, E. & Tsianou, M. Solution properties of dextran in water and in formamide. J. Appl. Polym. Sci. 125, 1681–1692 (2012).

74. Xin, H., Yu, X., Lu, Y. & Cai, J. Rheological properties and gelation behavior of chitosan in a weakly alkaline NaHCO_3_/urea aqueous solution. Macromolecules 58, 5283–5295 (2025).

75. Krause, W. E., Bellomo, E. G. & Colby, R. H. Rheology of sodium hyaluronate under physiological conditions. Biomacromolecules 2, 65–69 (2001).

76. Hinsch, H., Wilhelm, J. & Frey, E. Quantitative tube model for semiflexible polymer solutions. Eur. Phys. J. E 24, 35–46 (2007).

77. Isambert, H. & Maggs, A. C. Dynamics and rheology of actin solutions. Macromolecules 29, 1036–1040 (1996).

78. Broedersz, C. P. & MacKintosh, F. C. Modeling semiflexible polymer networks. Rev. Mod. Phys. 86, 995–1036 (2014).

79. Maestre-Reyna, M., Wu, W.-J. & Wang, A. H. J. Structural insights into RbmA, a biofilm scaffolding protein of *V. cholerae*. PLoS ONE 8, e82458 (2013).

80. Deng, Y., Sun, M. & Shaevitz, J. W. Direct measurement of cell wall stress stiffening and turgor pressure in live bacterial cells. Phys. Rev. Lett. 107, 158101 (2011).

81. Potapova, A. et al. Outer membrane vesicles and the outer membrane protein OmpU govern *Vibrio cholerae* biofilm matrix assembly. mBio 15, e03304–23 (2024).

82. Shivers, J. L. et al. Compression stiffening of fibrous networks with stiff inclusions. Proc. Natl. Acad. Sci. USA 117, 21037–21044 (2020).

83. Shivers, J. L., Feng, J. & MacKintosh, F. C. Criticality enhances the reinforcement of disordered networks by rigid inclusions. Phys. Rev. X 15, 031061 (2025).

84. Kumar, S. K., Ganesan, V. & Riggleman, R. A. Perspective: Outstanding theoretical questions in polymer-nanoparticle hybrids. J. Chem. Phys. 147, 020901 (2017).

85. Kumar, S. K. & Krishnamoorti, R. Nanocomposites: structure, phase behavior, and properties. Annu. Rev. Chem. Biomol. Eng. 1, 37–58 (2010).

86. Sattelle, B. M. & Almond, A. Microsecond kinetics in model single- and double-stranded amylose polymers. Phys. Chem. Chem. Phys. 16, 8119–8126 (2014).

87. Schieber, J. D. & Andreev, M. Entangled polymer dynamics in equilibrium and flow modeled through slip links. Annu. Rev. Chem. Biomol. Eng. 5, 367–381 (2014).

88. Nadell, C. D. & Bassler, B. L. A fitness trade-off between local competition and dispersal in *Vibrio cholerae* biofilms. Proc. Natl. Acad. Sci. USA 108, 14181–14185 (2011).

89. Nadell, C. D., Drescher, K., Wingreen, N. S. & Bassler, B. L. Extracellular matrix structure governs invasion resistance in bacterial biofilms. ISME J. 9, 1700–1709 (2015).

90. Beyhan, S. & Yildiz, F. H. Smooth to rugose phase variation in *Vibrio cholerae* can be mediated by a single nucleotide change that targets c-di-GMP signalling pathway. Mol. Microbiol. 63, 995–1007 (2007).

91. Dalia, A. B., McDonough, E. & Camilli, A. Multiplex genome editing by natural transformation. Proc. Natl. Acad. Sci. USA 111, 8937–8942 (2014).

92. Kovach, K. et al. Evolutionary adaptations of biofilms infecting cystic fibrosis lungs promote mechanical toughness by adjusting polysaccharide production. npj Biofilms Microbiomes 3, 1 (2017).

93. Grant, O. C. et al. Generating 3D models of complex carbohydrates with GLYCAM-Web. Nat. Methods 23, 720–723 (2026).

94. Kirschner, K. N. et al. GLYCAM06: A generalizable biomolecular force field. Carbohydrates. J. Comp. Chem. 29, 622–655 (2008).

95. Wang, J., Wolf, R. M., Caldwell, J. W., Kollman, P. A. & Case, D. A. Development and testing of a general amber force field. J. Comput. Chem. 25, 1157–1174 (2004).

96. Khalak, Y., Baumeier, B. & Karttunen, M. Improved general-purpose five-point model for water: TIP5P/2018. J. Chem. Phys. 149, 224507 (2018).

97. Roe, D. R. & Brooks, B. R. A protocol for preparing explicitly solvated systems for stable molecular dynamics simulations. J. Chem. Phys. 153, 054123 (2020).

98. Götz, A. W. et al. Routine microsecond molecular dynamics simulations with AMBER on GPUs. 1. Generalized born. J. Chem. Theory Comput. 8, 1542–1555 (2012).

99. Andersen, H. C. Rattle: A “velocity” version of the shake algorithm for molecular dynamics calculations. J. Comp. Phys. 52, 24–34 (1983).

100. Roe, D. R. & Cheatham III, T. E. PTRAJ and CPPTRAJ: Software for processing and analysis of molecular dynamics trajectory data. J. Chem. Theory Comput. 9, 3084–3095 (2013).

101. McGibbon, R. T. et al. MDTraj: a modern open library for the analysis of molecular dynamics trajectories. Biophys. J. 109, 1528–1532 (2015).

102. Pedregosa, F. et al. Scikit-learn: machine learning in Python. J. Mach. Learn. Res. 12, 2825–2830 (2011).

103. Bishop, C. M. Pattern Recognition and Machine Learning. (Springer New York, New York, NY, USA, 2006).

104. Banerjee, A., Dhillon, I. S., Ghosh, J. & Sra, S. Clustering on the unit hypersphere using von Mises-Fisher distributions. J. Mach. Learn. Res. 6, 1345–1382 (2005).

105. Weeks, J. D., Chandler, D. & Andersen, H. C. Role of repulsive forces in determining the equilibrium structure of simple liquids. J. Chem. Phys. 54, 5237–5247 (1971).

106. Allen, M. P. & Tildesley, D. J. Computer Simulation of Liquids. (Oxford University Press, Oxford, United Kingdom, 2017).

107. Plimpton, S. Fast parallel algorithms for short-range molecular dynamics. J. Comp. Phys. 117, 1–19 (1995).

108. Thompson, A. P. et al. LAMMPS - a flexible simulation tool for particle-based materials modeling at the atomic, meso, and continuum scales. Comput. Phys. Commun. 271, 108171 (2022).

109. Hoy, R. S., Foteinopoulou, K. & Kröger, M. Topological analysis of polymeric melts: Chain-length effects and fast-converging estimators for entanglement length. *Phys*. Rev. E 80, 031803 (2009).

110. Everaers, R. Topological versus rheological entanglement length in primitive-path analysis protocols, tube models, and slip-link models. *Phys*. Rev. E 86, 022801 (2012).

