## Supplementary Information for "Conformational dynamics of exopolysaccharides underlie biofilm matrix mechanics in *Vibrio cholerae*"

### Coarse-grained modeling of VPS chains and solutions

#### Supplementary Information for: Conformational dynamics of exopolysaccharides underlie biofilm matrix mechanics in *Vibrio cholerae*

Kee-Myoung Nam, Nathan Fowler, Rajan Kandel, Yongqi Zhu, Yuchu Liu, Yi-Jhen Lai, Muhammad Faheem Hassan, Emma Gerace, Merrill Asp, Rich Olson, Ying Li, Mu-Ping Nieh, Mingjiang Zhong, Robert J. Woods, Alexis Moreau, Jing Yan

**Abstract**

This document describes: (1) a modified Kremer–Grest model for semiflexible polymers, due to Svaneborg and Everaers 2020, which provides a simple estimate for the Kuhn length of VPS; and (2) the configurational-bias Monte Carlo (CBMC) procedure we used to sample configurations of VPS chains and solutions.

### 1 Modified Kremer–Grest model for semiflexible polymers

To obtain an initial estimate for the persistence length of VPS, we considered Svaneborg and Everaers' modified Kremer–Grest model for a semiflexible polymer (Svaneborg and Everaers 2020; Everaers et al. 2020). This model assumes that all non-bonded pairs of beads interact according to the repulsive component in the Weeks–Chandler–Andersen decomposition of the Lennard-Jones potential (Weeks et al. 1971), which is given by

$$U_{\mathrm{LJ}}(r) = \begin{cases} 4\epsilon\left(\left(\dfrac{\sigma}{r}\right)^{12} - \left(\dfrac{\sigma}{r}\right)^{6}\right) + \epsilon & \text{if } r < 2^{1/6}\sigma \\ 0 & \text{if } r \geq 2^{1/6}\sigma, \end{cases} \tag{1}$$

where $\epsilon = k_{\mathrm{B}}T$ and $\sigma$ is an interaction length scale. In addition, we assume that all bonds follow a FENE potential (Kremer and Grest 1990), which is given by

$$U_{\mathrm{FENE}}(r) = -\frac{k_{\mathrm{FENE}}R_0^2}{2}\ln\left(1 - \left(\frac{r}{R_0}\right)^2\right) + U_{\mathrm{LJ}}(r), \tag{2}$$

where $R_0$ is a maximum bond length—at which the potential diverges—and $k_{\mathrm{FENE}}$ is a stiffness parameter with units of energy/distance$^2$. Following Svaneborg and Everaers 2020, we set $R_0 = 1.5\sigma$, $k_{\mathrm{FENE}} = 30k_{\mathrm{B}}T\sigma^{-2}$, and assume an average bond length, $\langle \ell_0 \rangle$, of $\langle \ell_0 \rangle = 0.965\sigma$. Finally, we assume that all bond angles follow a harmonic potential with an equilibrium angle of $\pi$,[1]

$$U_{\text{angle}}(\theta) = \kappa_{\text{angle}} k_{\text{B}} T \left(1 + \cos\theta\right), \tag{3}$$

where $\kappa_{\text{angle}} k_{\text{B}} T$ is the bending stiffness.

Svaneborg and Everaers 2020 proposed the following empirical formula for the Kuhn length of the resulting polymer:

$$\begin{aligned} \ell_{\text{K}} &= \ell_{\text{K}}^{(0)} + 0.77\sigma \left(\tanh\left(-0.03\kappa_{\text{angle}}^2 - 0.41\kappa_{\text{angle}} + 0.16\right) + 1\right) \\ &= \ell_{\text{K}}^{(0)} + \frac{0.77}{0.965} \langle \ell_0 \rangle \left(\tanh\left(-0.03\kappa_{\text{angle}}^2 - 0.41\kappa_{\text{angle}} + 0.16\right) + 1\right) \end{aligned} \tag{4}$$

where $\ell_{\text{K}}^{(0)}$ is the bare Kuhn length, which is given here by

$$\ell_{\text{K}}^{(0)} = \langle \ell_0 \rangle \times \begin{cases} \left( \dfrac{2\kappa_{\text{angle}} + e^{-2\kappa_{\text{angle}}} - 1}{1 - e^{-2\kappa_{\text{angle}}} \left(2\kappa_{\text{angle}} + 1\right)} \right) & \text{if } \kappa_{\text{angle}} \neq 0 \\ 1 & \text{if } \kappa_{\text{angle}} = 0. \end{cases} \tag{5}$$

This formula for $\ell_{\text{K}}$ depends only on the mean bond length, $\langle \ell_0 \rangle$, and the dimensionless bending stiffness, $\kappa_{\text{angle}}$. The coarse-grained bond length distribution (Fig. S6A) suggest that we set $\langle \ell_0 \rangle = 1.8\,\text{nm}$, which corresponds to the head-to-tail length of a tetrasaccharide unit. For $\kappa_{\text{angle}}$, we note that the Boltzmann distribution corresponding to the bond angle potential (Eqn. 3) is proportional to

$$\begin{aligned} \exp\left\{-\frac{U_{\text{angle}}}{k_{\text{B}} T}\right\} &= \exp\left\{-\kappa_{\text{angle}}\left(1 + \cos\theta\right)\right\} \\ &= \exp\left\{-\kappa_{\text{angle}}\right\} \exp\left\{-\kappa_{\text{angle}} \cos\theta\right\} \\ &\propto \exp\left\{-\kappa_{\text{angle}} \cos\theta\right\} \\ &= \exp\left\{\kappa_{\text{angle}} \cos\left(\theta - \pi\right)\right\}, \end{aligned}$$

which implies that the Boltzmann distribution is in fact a von Mises distribution,

$$\frac{\exp\left\{\kappa_{\text{angle}} \cos\left(\theta - \pi\right)\right\}}{2\pi I_0(\kappa_{\text{angle}})},$$

with mean $\pi$ and concentration $\kappa_{\text{angle}}$; here, $I_0(\cdot)$ is the modified Bessel function of the first kind of order zero. As such, we set $\kappa_{\text{angle}}$ to the concentration of the major component in the von Mises mixture model for the coarse-grained bond angle distribution (Fig. S6A), which is ~20. Substituting $\langle \ell_0 \rangle = 1.8\,\text{nm}$ and $\kappa_{\text{angle}} = 20$ into Eqns. 4 and 5 yields $\ell_{\text{K}} \approx 70.2\,\text{nm}$.

---

[1] Note that Svaneborg and Everaers 2020 define this potential as

$$U_{\text{angle}}(\theta') = \kappa_{\text{angle}} k_{\text{B}} T \left(1 - \cos\theta'\right),$$

where $\theta' = \pi - \theta$ is the angle between consecutive bond vectors.

### 2 Configurational-bias Monte Carlo

#### 2.1 Preliminaries

##### 2.1.1 Potentials

We first describe the potentials that we used to model VPS. A more condensed discussion of this material is given in the Methods, and the values of all parameter values described below are given in Table S4.

We assumed that all non-bonded pairs of beads interact according to the repulsive component in the Weeks–Chandler–Andersen decomposition of the Lennard-Jones potential (Eqn. 1), with $\epsilon = k_\mathrm{B} T$; and that all bonds follow a FENE potential (Eqn. 2). We further assumed that all bond angles follow a two-component Gaussian potential (Milano et al. 2005),

$$U_{\mathrm{Gaussian}}(\theta) = -k_\mathrm{B} T \ln\left(\sum_{i=1}^{2} \frac{A_i}{w_i\sqrt{\pi/2}} \exp\left\{-\frac{2\left(\theta-\theta_i\right)^2}{w_i^2}\right\}\right). \tag{6}$$

The meanings of the parameters in this potential can be seen by taking the corresponding Boltzmann distribution,

$$\exp\left\{-\frac{U_{\mathrm{Gaussian}}(\theta)}{k_\mathrm{B} T}\right\} = \sum_{i=1}^{2} \frac{A_i}{w_i\sqrt{\pi/2}} \exp\left\{-\frac{2\left(\theta-\theta_i\right)^2}{w_i^2}\right\}, \tag{7}$$

which reveals that the Boltzmann distribution is a two-component Gaussian mixture, in which the $i$-th component has mean $\theta_i$, variance $w_i^2/4$, and weight $A_i$. Finally, we assume that all dihedral angles follow a single-welled, shifted harmonic potential of the form,

$$U_{\mathrm{dihedral}}(\varphi) = K_{\mathrm{dihedral}}\left(1+\cos\left(\varphi-\delta\right)\right), \tag{8}$$

which is minimized at $\varphi = \pi + \delta$.

To generate new chain configurations during MC sampling that have high acceptance probabilities, we must first be able to sample bond lengths, bond angles, and dihedral angles that follow these potentials (or rather, the corresponding Boltzmann distributions). Specifically, we sampled each bond length from a density of the form,

$$p(r) \propto r^2 \exp\left\{-\frac{U_{\mathrm{FENE}}(r)}{k_\mathrm{B} T}\right\}, \tag{9}$$

where the prefactor, $r^2$, is the radial component of the Jacobian of the transformation of a volume element in three-dimensional space, $d^3\mathbf{r}$, into spherical coordinates, $r^2\left(\sin\theta\right) dr\, d\theta\, d\varphi$ (Allen and Tildesley 2017). This prefactor accounts for the fact that we are not merely sampling a bond length $r$, but rather a *bond vector with length* $r$, and given a fixed starting point $\mathbf{u}$, the probability of selecting a bond vector with length $r$ should scale with the surface area of the sphere of radius $r$ centered at $\mathbf{u}$.

To sample each bond angle $\theta$, we first sampled an angle $\theta' \in [-\pi, \pi)$ from an "unraveled" four-component von Mises mixture of the form,

$$p(\theta') \propto \left|\sin\theta'\right| \sum_{i=1}^{4} \frac{B_i \exp\left\{\left(4/w_i^2\right)\cos\left(\theta' - \theta_i\right)\right\}}{2\pi I_0(4/w_i^2)}, \tag{10}$$

where we have added two new components (indices 3 and 4) that "mirror" the components in the true bond angle distribution (indices 1 and 2) about $\theta' = 0$, with

$$\theta_3 = -\theta_1, \qquad \theta_4 = -\theta_2, \qquad w_3 = w_1, \qquad w_4 = w_2;$$

and the mixture weights have been apportioned appropriately through the unraveling,

$$B_1 = B_3 = \frac{A_1}{2} \qquad \text{and} \qquad B_2 = B_4 = \frac{A_2}{2}.$$

We have also introduced a similar geometric prefactor of $\left|\sin\theta'\right|$ in Eqn. 10 to that in Eqn. 9, to similarly account for the degeneracy in bond vectors that give rise to the same bond angle. We then set $\theta$ to the absolute value, $\theta = \left|\theta'\right|$, to ensure that the bond angle lies in $[0, \pi]$. Here, we have used the fact that a von Mises distribution with mean $\mu$ and large concentration, $\kappa \gg 1$, is well-approximated by a Gaussian distribution with mean $\mu$ and variance $1/\kappa$; this assumption is appropriate since the variances in the Gaussian potential, $w_i^2/4$, are quite small ($\sim 1/20$, Table S4).

Finally, we sampled each dihedral angle from the Boltzmann distribution corresponding to $U_{\text{dihedral}}(\varphi)$, which is the von Mises distribution with mean $\pi + \delta$ and concentration $\kappa_{\text{dihedral}} = K_{\text{dihedral}}/(k_B T)$:

$$p(\varphi) = \frac{\exp\left\{\kappa_{\text{dihedral}} \cos\left(\varphi - \pi - \delta\right)\right\}}{2\pi I_0(\kappa_{\text{dihedral}})}.$$

Sampling from $p(\varphi)$ is straightforward (Best and Fisher 1979). To sample from $p(\theta)$, we used rejection sampling: we iteratively sampled an angle, $\theta' \in [-\pi, \pi)$, from the von Mises mixture (without the $\left|\sin\theta'\right|$ prefactor), accepted $\theta'$ with probability $\left|\sin\theta'\right|$, then set $\theta = \left|\theta'\right|$. To sample from $p(r)$, we first calculated an empirical cumulative distribution function for the density,

$$F_{\text{FENE}}(r) = \left(\int_0^r p(r')\,dr'\right) \Big/ \left(\int_0^{R_0} p(r')\,dr'\right), \qquad r \in (0, R_0),$$

and returned $F_{\text{FENE}}^{-1}(u)$, where $u$ is sampled uniformly from $[0,1]$. In practice, we calculated and normalized $F_{\text{FENE}}(r)$ over a mesh $\mathscr{M}$ of $10^4$ values over $[10^{-6}, R_0 - 10^{-6}]$ and, for each sampled value of $u$, we approximated $F_{\text{FENE}}^{-1}(u)$ by linearly interpolating between the two values, $u_1, u_2 \in F_{\text{FENE}}(\mathscr{M})$, nearest to $u$ in the image of the mesh (with $u_1 < u < u_2$), as

$$F_{\text{FENE}}^{-1}(u) \approx F_{\text{FENE}}^{-1}(u_1) + (u - u_1)\,\frac{F_{\text{FENE}}^{-1}(u_2) - F_{\text{FENE}}^{-1}(u_1)}{u_2 - u_1}.$$

##### 2.1.2 Generating new bead positions

Let $\mathbf{r}_1$, $\mathbf{r}_2$, and $\mathbf{r}_3$ be the positions of three beads, with bonds between $\mathbf{r}_1$ and $\mathbf{r}_2$ and between $\mathbf{r}_2$ and $\mathbf{r}_3$. Let $\mathbf{u}_1 = \mathbf{r}_2 - \mathbf{r}_1$ and $\mathbf{u}_2 = \mathbf{r}_3 - \mathbf{r}_2$ be the vectors along these bonds, and let $\hat{\mathbf{u}}_1 = \mathbf{u}_1 / \|\mathbf{u}_1\|$ and $\hat{\mathbf{u}}_2 = \mathbf{u}_2 / \|\mathbf{u}_2\|$ be the corresponding unit vectors. We now describe our procedure for generating a new bead position,

$$\mathbf{r}_4 = \mathbf{q}(\mathbf{r}_1, \mathbf{r}_2, \mathbf{r}_3, r, \theta, \varphi), \tag{11}$$

such that the bond length between $\mathbf{r}_3$ and $\mathbf{r}_4$ is $r$,

$$\|\mathbf{r}_4 - \mathbf{r}_3\| = r;$$

the bond angle between $\mathbf{r}_2$, $\mathbf{r}_3$, and $\mathbf{r}_4$ is $\theta$,

$$\angle(\mathbf{r}_2, \mathbf{r}_3, \mathbf{r}_4) = \theta;$$

and the dihedral angle along the four-bead segment is $\varphi$, which we denote by

$$\angle_{\mathrm{D}}(\mathbf{r}_1, \mathbf{r}_2, \mathbf{r}_3, \mathbf{r}_4) = \varphi.$$

To do this, we first define a (right-handed) orthonormal basis for $\mathbb{R}^3$ centered at $\mathbf{r}_3$, given by

$$\hat{\mathbf{e}}_1 = -\hat{\mathbf{u}}_2, \qquad \hat{\mathbf{e}}_2 = \frac{\hat{\mathbf{u}}_1 \times \hat{\mathbf{u}}_2}{\|\hat{\mathbf{u}}_1 \times \hat{\mathbf{u}}_2\|}, \qquad \hat{\mathbf{e}}_3 = \hat{\mathbf{e}}_1 \times \hat{\mathbf{e}}_2.$$

Then we can define a unit vector, $\hat{\mathbf{v}}$, from $\mathbf{r}_3$ that enforces the correct bond angle and dihedral angle, as

$$\hat{\mathbf{v}} = (\cos\theta)\,\hat{\mathbf{e}}_1 + (\sin\theta)\left(\left(\cos\varphi\right)\hat{\mathbf{e}}_3 + \left(\sin\varphi\right)\hat{\mathbf{e}}_2\right). \tag{12}$$

To see that $\hat{\mathbf{v}}$ specifies the desired direction, we can turn to Rodrigues' rotation formula, which states that rotating a vector, $\mathbf{v} \in \mathbb{R}^3$, by an angle $\gamma$ about an axis specified by a unit vector $\hat{\mathbf{n}}$ yields the vector,

$$\mathbf{v}' = \mathbf{v}\cos\gamma + (\hat{\mathbf{n}} \times \mathbf{v})\sin\gamma + \hat{\mathbf{n}}\,(\hat{\mathbf{n}} \cdot \mathbf{v})\left(1 - \cos\gamma\right). \tag{13}$$

Now, to first enforce a bond angle of $\theta$, we can first rotate $\hat{\mathbf{e}}_1 = -\hat{\mathbf{u}}_2 \propto \mathbf{r}_2 - \mathbf{r}_3$ by $\theta$ within the plane spanned by $\hat{\mathbf{e}}_1$ and $\hat{\mathbf{e}}_3$, to get

$$\hat{\mathbf{w}} = (\cos\theta)\,\hat{\mathbf{e}}_1 + (\sin\theta)\,\hat{\mathbf{e}}_3.$$

It is easy to check that $\hat{\mathbf{w}}$ is a unit vector with $\hat{\mathbf{w}} \cdot \hat{\mathbf{e}}_1 = \cos\theta$. Moreover, it lies within the plane spanned by $\hat{\mathbf{e}}_1$ and $\hat{\mathbf{e}}_3$, which, by construction, is exactly the plane spanned by $\hat{\mathbf{u}}_1$ and $\hat{\mathbf{u}}_2$. Because $\hat{\mathbf{e}}_1$, $\hat{\mathbf{e}}_2$, and $\hat{\mathbf{e}}_3$ form a right-handed basis, the dihedral angle along the four-bead segment formed by $\mathbf{r}_1$, $\mathbf{r}_2$, $\mathbf{r}_3$, and $\mathbf{r}_3 + a\hat{\mathbf{w}}$ for any $a > 0$ is zero,

$$\angle_{\mathrm{D}}(\mathbf{r}_1, \mathbf{r}_2, \mathbf{r}_3, \mathbf{r}_3 + a\hat{\mathbf{w}}) = 0.$$

We can now enforce a dihedral angle of $\varphi$ by rotating $\hat{\mathbf{w}}$ about the axis specified by $-\hat{\mathbf{e}}_1 = \hat{\mathbf{u}}_2$. Using Eqn. 13, we obtain

$$\hat{\mathbf{v}} = \hat{\mathbf{w}}\cos\varphi + (-\hat{\mathbf{e}}_1 \times \hat{\mathbf{w}})\sin\varphi - \hat{\mathbf{e}}_1\,(-\hat{\mathbf{e}}_1 \cdot \hat{\mathbf{w}})\left(1 - \cos\varphi\right)$$

$$
\begin{aligned}
&= ((\cos\theta)\,\hat{\mathbf{e}}_1 + (\sin\theta)\,\hat{\mathbf{e}}_3)\cos\varphi - (\hat{\mathbf{e}}_1 \times ((\cos\theta)\,\hat{\mathbf{e}}_1 + (\sin\theta)\,\hat{\mathbf{e}}_3))\sin\varphi + \hat{\mathbf{e}}_1\,(\cos\theta)\left(1-\cos\varphi\right)\\
&= \left(\cos\theta\cos\varphi\right)\hat{\mathbf{e}}_1 + \left(\sin\theta\cos\varphi\right)\hat{\mathbf{e}}_3 - \left(\sin\theta\sin\varphi\right)(\hat{\mathbf{e}}_1 \times \hat{\mathbf{e}}_3) + (\cos\theta)\,\hat{\mathbf{e}}_1 - \left(\cos\theta\cos\varphi\right)\hat{\mathbf{e}}_1\\
&= (\cos\theta)\,\hat{\mathbf{e}}_1 + \left(\sin\theta\cos\varphi\right)\hat{\mathbf{e}}_3 + \left(\sin\theta\sin\varphi\right)\hat{\mathbf{e}}_2\\
&= (\cos\theta)\,\hat{\mathbf{e}}_1 + (\sin\theta)\left(\left(\cos\varphi\right)\hat{\mathbf{e}}_3 + \left(\sin\varphi\right)\hat{\mathbf{e}}_2\right),
\end{aligned}
$$

i.e., we recover Eqn. 12. Now, we can finally define the new bead position as

$$\mathbf{r}_4 = \mathbf{q}(\mathbf{r}_1, \mathbf{r}_2, \mathbf{r}_3, r, \theta, \varphi) = \mathbf{r}_3 + r\hat{\mathbf{v}}.$$

##### 2.1.3 Determining mixture weights in the bond angle distribution

We now consider the problem of setting the mixture weights, $A_1$ and $A_2$, in the two-component Gaussian potential to obtain a desired "kink fraction." This is not an immediately trivial problem, due to the presence of a $\sin\theta'$ geometric prefactor that biases the sampling distribution in Eqn. 10. Specifically, recall that our sampling procedure for the bond angles involves first sampling an angle, $\theta'$, from an "unraveled" distribution of the form (Eqn. 10),

$$p(\theta') \propto \left|\sin\theta'\right| \sum_{i=1}^{4} \frac{B_i \exp\left\{\left(4/w_i^2\right)\cos\left(\theta' - \theta_i\right)\right\}}{2\pi I_0(4/w_i^2)},$$

where $\theta_3 = -\theta_1$, $\theta_4 = -\theta_2$, $w_3 = w_1$, $w_4 = w_2$, $B_1 = B_3 = A_1/2$, and $B_2 = B_4 = A_2/2$; and setting the bond angle, $\theta$, as $\theta = \left|\theta'\right|$. As such, $A_1$ and $A_2$ specify the relative contributions of the two pairs of matching components before the geometric re-weighting by the $\sin\theta'$ prefactor, which generally do not coincide with their contributions after the re-weighting.

For notational simplicity, let us write $f_{\mathrm{VM}}(\cdot)$ for the von Mises density,

$$f_{\mathrm{VM}}(\theta' \mid \mu, \kappa) = \frac{\exp\left\{\kappa\cos\left(\theta' - \mu\right)\right\}}{2\pi I_0(\kappa)},$$

so that the $i$-th component in the above distribution is given by $f_{\mathrm{VM}}(\theta' \mid \theta_i, 4/w_i^2)$. Then the total contribution of components 1 and 3 to the sampling distribution is given by

$$\int_{-\pi}^{\pi} \left|\sin\theta'\right| \left(B_1 f_{\mathrm{VM}}(\theta' \mid \theta_1, 4/w_1^2) + B_3 f_{\mathrm{VM}}(\theta' \mid \theta_3, 4/w_3^2)\right) d\theta',$$

which we can rewrite as

$$
\begin{aligned}
&B_1 \int_{-\pi}^{\pi} \left|\sin\theta'\right| f_{\mathrm{VM}}(\theta' \mid \theta_1, 4/w_1^2)\, d\theta' + B_3 \int_{-\pi}^{\pi} \left|\sin\theta'\right| f_{\mathrm{VM}}(\theta' \mid \theta_3, 4/w_3^2)\, d\theta'\\
&= \frac{A_1}{2} \int_{-\pi}^{\pi} \left|\sin\theta'\right| f_{\mathrm{VM}}(\theta' \mid \theta_1, 4/w_1^2)\, d\theta' + \frac{A_1}{2} \int_{-\pi}^{\pi} \left|\sin\theta'\right| f_{\mathrm{VM}}(\theta' \mid -\theta_1, 4/w_1^2)\, d\theta'\\
&= \frac{A_1 Z_1}{2},
\end{aligned}
$$

where we have defined

$$Z_1 = \int_{-\pi}^{\pi} \left|\sin\theta'\right| \left(f_{\mathrm{VM}}(\theta' \mid \theta_1, 4/w_1^2) + f_{\mathrm{VM}}(\theta' \mid -\theta_1, 4/w_1^2)\right) d\theta'.$$

Similarly, the total contribution of components 2 and 4 to the sampling distribution is $A_2 Z_2/2$, where

$$Z_2 = \int_{-\pi}^{\pi} \left|\sin\theta'\right| \left( f_{\text{VM}}(\theta' \mid \theta_2, 4/w_2^2) + f_{\text{VM}}(\theta' \mid \theta_4, 4/w_4^2) \right) d\theta'.$$

Now, the fractions of bond angles arising from the two pairs of matching components in the sampling distribution are

$$F_1 = \frac{A_1 Z_1}{A_1 Z_1 + A_2 Z_2} \qquad \text{and} \qquad F_2 = \frac{A_2 Z_2}{A_1 Z_1 + A_2 Z_2}.$$

We would like to set $A_1$ and $A_2$ such that we obtain a desired set of values for $F_1$ and $F_2$, the latter of which represents the kink fraction. To do this, we first note that

$$\frac{F_1}{F_2} = \frac{A_1 Z_1}{A_2 Z_2},$$

which we can rearrange to get

$$\frac{A_1}{A_2} = \frac{F_1 Z_2}{F_2 Z_1}.$$

We can then use the fact that $A_2 = 1 - A_1$ and rearrange, to get

$$A_1 = \frac{F_1 Z_2}{F_2 Z_1}(1 - A_1) \qquad \Longrightarrow \qquad A_1 = \frac{(F_1 Z_2)/(F_2 Z_1)}{1 + (F_1 Z_2)/(F_2 Z_1)},$$

and

$$A_2 = \frac{1}{1 + (F_1 Z_2)/(F_2 Z_1)}.$$

We can now numerically evaluate $Z_1$ and $Z_2$ and utilize these formulas to calculate $A_1$ and $A_2$ for any given choice of $F_1$ and $F_2$. For instance, undertaking this calculation for $F_1 = 0.95$, corresponding to a kink fraction of $F_2 = 5\%$, yields $A_1 \approx 0.98168$.

#### 2.2 Configurational-bias Monte Carlo

Here, we describe our configurational-bias Monte Carlo (CBMC) procedure for sampling polymer configurations (Siepmann and Frenkel 1992). The basic procedure iteratively generates a sequence of configurations,

$$\left\{ X^{(n)} = \left( \mathbf{r}_1(X^{(n)}), \ldots, \mathbf{r}_N(X^{(n)}) \right) : n = 0, 1, \ldots \right\},$$

given an initial configuration $X^{(0)}$. In particular, we construct $X^{(n+1)}$ from $X^{(n)}$ as follows:

1. Randomly choose one of three move types: *single-bead reptation*, *multi-bead reptation*, or *terminal segment move*.

2. Randomly choose the direction of the chosen move type (reptation towards the head or tail of the chain, or moving the terminal segment at the head or tail of the chain).

3. Generate a new candidate configuration, $X_{\text{new}}$, according to the procedure for each move type, and calculate their corresponding *forward and reverse Rosenbluth factors*, $W_{\text{fwd}}$ and $W_{\text{rev}}$, as described below.

4. Accept the candidate configuration (i.e., set $X^{(n+1)} = X_{\text{new}}$), with probability

$$p_{\text{accept}} = \min\left(1, \frac{W_{\text{fwd}}}{W_{\text{rev}}}\right), \tag{14}$$

and reject otherwise (in which case we set $X^{(n+1)} = X^{(n)}$).

We now describe how to generate the new configuration, $X_{\text{new}}$, and calculate $W_{\text{fwd}}$ and $W_{\text{rev}}$, for each move type. In what follows, we let $b_1, \ldots, b_N$ be the beads in the $n$-th configuration and, to simplify our notation, we denote their positions by $\mathbf{r}_j^{(n)} = \mathbf{r}_j$. We also direct the reader to Allen and Tildesley's discussion of CBMC (Allen and Tildesley 2017, §9.3.4) for further background.

##### 2.2.1 Single- and multi-bead reptation

In these move types, we reptate the chain by $K \geq 1$ beads: we generate a segment of $K \geq 1$ new beads, with positions

$$\mathbf{r}^{\text{new}}(1), \qquad \ldots, \qquad \mathbf{r}^{\text{new}}(K),$$

where $\mathbf{r}^{\text{new}}(1)$ is bonded to the terminal bead at one end of the chain ($b_N$ or $b_1$), and remove the $K$ beads at the other end of the chain ($b_1, \ldots, b_K$ or $b_{N-K+1}, \ldots, b_N$, respectively). We call these cases "reptating towards the tail" and "reptating towards the head" of the chain, respectively. Since the procedure for single-bead reptation ($K = 1$) is merely an example of the general procedure for $K \geq 1$, we first describe the general procedure, then explain how it reduces in the single-bead case.

Let us suppose that we are reptating towards the tail; the analogous procedure for reptation towards the head can be inferred straightforwardly. Following the standard prescription for CBMC (Allen and Tildesley 2017), we generate the segment sequentially, by conditioning each bead position, $\mathbf{r}^{\text{new}}(i)$, on the previously generated bead positions, $\mathbf{r}^{\text{new}}(1), \ldots, \mathbf{r}^{\text{new}}(i-1)$. For each $i = 1, \ldots, K$, we generate $\mathbf{r}^{\text{new}}(i)$ by first proposing $M$ candidate positions,

$$\mathbf{r}_j^{\text{new}}(i) \qquad \text{for } j = 1, \ldots, M,$$

each of which is specified as

$$\mathbf{r}_j^{\text{new}}(i) = \begin{cases} \mathbf{q}(\mathbf{r}_{N-2}, \mathbf{r}_{N-1}, \mathbf{r}_N, r_{i,j}, \theta_{i,j}, \varphi_{i,j}) & \text{if } i = 1 \\ \mathbf{q}(\mathbf{r}_{N-1}, \mathbf{r}_N, \mathbf{r}^{\text{new}}(1), r_{i,j}, \theta_{i,j}, \varphi_{i,j}) & \text{if } i = 2 \\ \mathbf{q}(\mathbf{r}_N, \mathbf{r}^{\text{new}}(1), \mathbf{r}^{\text{new}}(2), r_{i,j}, \theta_{i,j}, \varphi_{i,j}) & \text{if } i = 3 \\ \mathbf{q}(\mathbf{r}^{\text{new}}(i-3), \mathbf{r}^{\text{new}}(i-2), \mathbf{r}^{\text{new}}(i-1), r_{i,j}, \theta_{i,j}, \varphi_{i,j}) & \text{otherwise,} \end{cases} \tag{15}$$

where each bond length $r_{i,j}$, bond angle $\theta_{i,j}$, and dihedral angle $\varphi_{i,j}$ are sampled as described previously.

We then select one of these $M$ candidate positions according to their Boltzmann weights, $w_j(i)$, each of which is given by

$$w_j(i) = \exp\left\{-\frac{V_j(i)}{k_\mathrm{B}T}\right\}, \tag{16}$$

where $V_j(i)$ is the non-bonded interaction energy between the new bead and every bead in the original configuration, $X^{(n)}$, that would remain in the reptated configuration, plus the new beads that have been generated thus far. To be more precise,

$$V_j(i) = \sum_{k=K+1}^{N_\mathrm{max}} U_\mathrm{LJ}\left(\left\|\mathbf{r}_j^\mathrm{new}(i) - \mathbf{r}_k\right\|\right) + \sum_{k=1}^{i-2} U_\mathrm{LJ}\left(\left\|\mathbf{r}_j^\mathrm{new}(i) - \mathbf{r}^\mathrm{new}(k)\right\|\right), \tag{17}$$

where

$$N_\mathrm{max} = \begin{cases} N & \text{if } i > 1 \\ N-1 & \text{if } i = 1. \end{cases}$$

The first sum in Eqn. 17 omits the $K$ beads in $X^{(n)}$ that would be removed upon reptation, as well as the final bead, $b_N$, in the case where we are generating the very first bead in the growing segment ($i = 1$). Furthermore, the second sum omits the $(i-1)$-th bead in the growing segment, which is bonded to the $i$-th bead.

From here, we obtain the $i$-th *Rosenbluth weight*,

$$W_i = \sum_{j=1}^{M} w_j(i), \tag{18}$$

and we select the $j$-th candidate bead position, $\mathbf{r}^\mathrm{new}(i) = \mathbf{r}_j^\mathrm{new}(i)$, with probability $w_j(i)/W_i$. Continuing in this fashion for each $i = 1, \ldots, K$, we obtain the full segment of $K$ beads and the Rosenbluth weights $W_1, \ldots, W_K$, whose product is the *forward Rosenbluth factor*:

$$W_\mathrm{fwd} = \prod_{i=1}^{K} W_i. \tag{19}$$

We now turn to the task of calculating $W_\mathrm{rev}$. Let $\mathbf{r}'_1, \ldots, \mathbf{r}'_N$ denote the bead positions in the *reptated* configuration, which are given by

$$\mathbf{r}'_m = \begin{cases} \mathbf{r}_{m+K} & \text{if } m \le N-K \\ \mathbf{r}^\mathrm{new}(N-m) & \text{if } m > N-K. \end{cases} \tag{20}$$

We now undertake the same calculation as before, but in reverse: we start from the reptated configuration, and calculate the Rosenbluth factor arising from *reversion* to the original configuration. For each $i = 1, \ldots, K$, we first propose $M-1$ positions for the $i$-th bead in this segment, which we define, for $j = 1, \ldots, M-1$, as

$$\mathbf{r}_j^\mathrm{rev}(i) = \begin{cases} \mathbf{q}(\mathbf{r}'_3, \mathbf{r}'_2, \mathbf{r}'_1, r_{i,j}, \theta_{i,j}, \varphi_{i,j}) & \text{if } i = 1 \\ \mathbf{q}(\mathbf{r}'_2, \mathbf{r}'_1, \mathbf{r}_K, r_{i,j}, \theta_{i,j}, \varphi_{i,j}) & \text{if } i = 2 \\ \mathbf{q}(\mathbf{r}'_1, \mathbf{r}_K, \mathbf{r}_{K-1}, r_{i,j}, \theta_{i,j}, \varphi_{i,j}) & \text{if } i = 3 \\ \mathbf{q}(\mathbf{r}_{K-i+4}, \mathbf{r}_{K-i+3}, \mathbf{r}_{K-i+2}, r_{i,j}, \theta_{i,j}, \varphi_{i,j}) & \text{otherwise,} \end{cases} \tag{21}$$

where, again, each bond length $r_{i,j}$, bond angle $\theta_{i,j}$, and dihedral angle $\varphi_{i,j}$ are sampled as described previously. We also include reversion to the original configuration as an additional bead position: for each $i = 1, \ldots, K$, we set

$$\mathbf{r}_M^{\text{rev}}(i) = \mathbf{r}_{K-i+1}. \tag{22}$$

We note that the key difference between the forward and reverse calculations is that, in the former, we *generate a new segment* of $K$ beads, and each successive collection of candidate bead positions, $\mathbf{r}_j^{\text{new}}(i)$, depends on the previously chosen bead positions, $\mathbf{r}^{\text{new}}(1), \ldots, \mathbf{r}^{\text{new}}(i-1)$. Here, while we do generate candidate bead positions for each $i$, we merely use these candidate positions to calculate the corresponding Rosenbluth weights (as described below), and forgo stitching together a new segment of beads.

We then calculate the Boltzmann weight, $w_j^{\text{rev}}(i)$, which is given by

$$w_j^{\text{rev}}(i) = \exp\left\{-\frac{V_j^{\text{rev}}(i)}{k_{\text{B}} T}\right\}, \tag{23}$$

where $V_j^{\text{rev}}(i)$ follows a similar form to $V_j(i)$ in Eqn. 17:

$$V_j^{\text{rev}}(i) = \sum_{k=N_{\min}}^{N-K} U_{\text{LJ}}\left(\left\|\mathbf{r}_j^{\text{rev}}(i) - \mathbf{r}'_k\right\|\right) + \sum_{k=1}^{i-2} U_{\text{LJ}}\left(\left\|\mathbf{r}_j^{\text{rev}}(i) - \mathbf{r}_{K-k+1}\right\|\right), \tag{24}$$

where

$$N_{\min} = \begin{cases} 2 & \text{if } i = 1 \\ 1 & \text{if } i > 1. \end{cases}$$

From here, we can calculate the $i$-th *reverse Rosenbluth weight*,

$$W_i^{\text{rev}} = \sum_{j=1}^{M} w_j^{\text{rev}}(i), \tag{25}$$

which we can combine to arrive at the *reverse Rosenbluth factor*,

$$W_{\text{rev}} = \prod_{i=1}^{K} W_i^{\text{rev}}, \tag{26}$$

which we can use to finally calculate the acceptance probability via Eqn. 14.

**The single-bead case.** It is instructive to consider how this procedure simplifies in the single-bead case ($K = 1$). In this case, we generate a single new bead position, $\mathbf{r}^{\text{new}} \equiv \mathbf{r}^{\text{new}}(1)$, by first randomly sampling $M$ candidate bead positions (Eqn. 15),

$$\mathbf{r}_j^{\text{new}} \equiv \mathbf{r}_j^{\text{new}}(1) = \mathbf{q}(\mathbf{r}_{N-2}, \mathbf{r}_{N-1}, \mathbf{r}_N, r_{1,j}, \theta_{1,j}, \varphi_{1,j}).$$

The corresponding Boltzmann weight is given by (Eqn. 16)

$$w_j(1) = \exp\left\{-\frac{V_j(1)}{k_\mathrm{B} T}\right\},$$

where the non-bonded interaction energy, $V_j(1)$, is given by (Eqn. 17)

$$V_j(1) = \sum_{k=2}^{N-1} U_\mathrm{LJ}\left(\left\|\mathbf{r}_j^\mathrm{new} - \mathbf{r}_k\right\|\right).$$

Summing the Boltzmann weights yields the Rosenbluth weight, $W_1$, which in this case is also the forward Rosenbluth factor (Eqns. 18 and 19):

$$W_\mathrm{fwd} = W_1 = \sum_{j=1}^{M} w_j(1).$$

To calculate $W_\mathrm{rev}$, we again calculate the Rosenbluth factor arising from reversion to the original configuration. The bead positions in the reptated configuration are given by (Eqn. 20)

$$\mathbf{r}'_1 = \mathbf{r}_2, \qquad \ldots, \qquad \mathbf{r}'_{N-1} = \mathbf{r}_N, \qquad \mathbf{r}'_N = \mathbf{r}^\mathrm{new},$$

and we randomly sample $M-1$ new bead positions at the *head* of the reptated configuration (Eqn. 21),

$$\mathbf{r}_j^\mathrm{rev} \equiv \mathbf{r}_j^\mathrm{rev}(1) = \mathbf{q}(\mathbf{r}'_3, \mathbf{r}'_2, \mathbf{r}'_1, r_{1,j}, \theta_{1,j}, \varphi_{1,j}).$$

We also add reversion to the original configuration as the final reverse move, by setting (Eqn. 22)

$$\mathbf{r}_M^\mathrm{rev}(1) = \mathbf{r}_1.$$

We can then calculate the Boltzmann weight of each reverse move, as (Eqns. 23 and 24)

$$w_j^\mathrm{rev}(1) = \exp\left\{-\frac{V_j^\mathrm{rev}(1)}{k_\mathrm{B} T}\right\}, \qquad \text{where} \qquad V_j^\mathrm{rev}(1) = \sum_{k=2}^{N-1} U_\mathrm{LJ}\left(\left\|\mathbf{r}_j^\mathrm{rev} - \mathbf{r}'_k\right\|\right),$$

whose sum is the reverse Rosenbluth factor (Eqns. 25 and 26),

$$W_\mathrm{rev} = W_1^\mathrm{rev} = \sum_{j=1}^{M} w_j^\mathrm{rev}(1).$$

##### 2.2.2 Terminal segment move

In this move type, we move the $K > 1$ beads at either the head or tail of the chain. In other words, we seek to remove either the first $K$ beads or the final $K$ beads in the chain, and "regrow" a new segment of $K$ beads, with positions

$$\mathbf{r}^\mathrm{new}(1), \qquad \ldots, \qquad \mathbf{r}^\mathrm{new}(K),$$

where $\mathbf{r}^{\text{new}}(1)$ is bonded to $b_{K+1}$ (if we have removed the first $K$ beads) or to $b_{N-K}$ (if we have removed the final $K$ beads).

To do this, we follow a very similar procedure to that for multi-bead reptation. Let us assume that we are moving the final $K$ beads in the chain. Specifically, for each $i = 1, \ldots, K$, we generate $\mathbf{r}^{\text{new}}(i)$ by proposing $M$ candidate positions, $\mathbf{r}_j^{\text{new}}(i)$ for $j = 1, \ldots, M$, as

$$\mathbf{r}_j^{\text{new}}(i) = \begin{cases} \mathbf{q}(\mathbf{r}_{N-K-2}, \mathbf{r}_{N-K-1}, \mathbf{r}_{N-K}, r_{i,j}, \theta_{i,j}, \varphi_{i,j}) & \text{if } i = 1 \\ \mathbf{q}(\mathbf{r}_{N-K-1}, \mathbf{r}_{N-K}, \mathbf{r}^{\text{new}}(1), r_{i,j}, \theta_{i,j}, \varphi_{i,j}) & \text{if } i = 2 \\ \mathbf{q}(\mathbf{r}_{N-K}, \mathbf{r}^{\text{new}}(1), \mathbf{r}^{\text{new}}(2), r_{i,j}, \theta_{i,j}, \varphi_{i,j}) & \text{if } i = 3 \\ \mathbf{q}(\mathbf{r}^{\text{new}}(i-3), \mathbf{r}^{\text{new}}(i-2), \mathbf{r}^{\text{new}}(i-1), r_{i,j}, \theta_{i,j}, \varphi_{i,j}) & \text{otherwise,} \end{cases} \tag{27}$$

where each bond length $r_{i,j}$, bond angle $\theta_{i,j}$, and dihedral angle $\varphi_{i,j}$ are sampled as described previously. (Note that the only difference between this proposal and that in Eqn. 15 is that $\mathbf{r}_j^{\text{new}}(1)$ is bonded to $\mathbf{r}_{N-K}$, not $\mathbf{r}_N$.) We then calculate the corresponding Boltzmann weight, $w_j(i)$, as

$$w_j(i) = \exp\left\{-\frac{V_j(i)}{k_{\text{B}}T}\right\},$$

where, again, $V_j(i)$ is the non-bonded interaction energy between the new bead and every bead in the original configuration, $X^{(n)}$, that would remain in the new configuration, plus the new beads that have been generated thus far. Naturally, this energy follows a similar form to that in Eqn. 17:

$$V_j(i) = \sum_{k=1}^{N_{\max}} U_{\text{LJ}}\left(\left\|\mathbf{r}_j^{\text{new}}(i) - \mathbf{r}_k\right\|\right) + \sum_{k=1}^{i-2} U_{\text{LJ}}\left(\left\|\mathbf{r}_j^{\text{new}}(i) - \mathbf{r}^{\text{new}}(k)\right\|\right), \tag{28}$$

where

$$N_{\max} = \begin{cases} N-K & \text{if } i > 1 \\ N-K-1 & \text{if } i = 1. \end{cases}$$

The first sum in Eqn. 28 omits the final $K$ beads in $X^{(n)}$, as well as the final bead in the fixed segment, $b_{N-K}$, in the case where we are generating the very first bead in the growing segment ($i = 1$). Furthermore, the second sum omits the $(i-1)$-th bead in the growing segment, which is bonded to the $i$-th bead.

From here, we follow precisely the same procedure for choosing $\mathbf{r}^{\text{new}}(i)$ and calculating $W_{\text{fwd}}$ as in multi-bead reptation: we calculate the $i$-th Rosenbluth weight,

$$W_i = \sum_{j=1}^{M} w_j(i),$$

and we select the $j$-th candidate bead position, $\mathbf{r}^{\text{new}}(i) = \mathbf{r}_j^{\text{new}}(i)$, with probability $w_j(i)/W_i$. Upon generating the full segment of $K$ beads, we obtain the forward Rosenbluth factor,

$$W_{\text{fwd}} = \prod_{i=1}^{K} W_i.$$

We now calculate $W_{\text{rev}}$, which is again the Rosenbluth factor arising from reversion to the original configuration. Again, let $\mathbf{r}'_1, \ldots, \mathbf{r}'_N$ denote the bead positions in the new configuration, which are given here by

$$\mathbf{r}'_m = \begin{cases} \mathbf{r}_m & \text{if } m \leq N-K \\ \mathbf{r}^{\text{new}}(m) & \text{if } m > N-K. \end{cases}$$

We now randomly sample, for each $i = 1, \ldots, K$, a collection of $M-1$ candidate positions for the $i$-th beads at the *same end* of the chain, with positions $\mathbf{r}^{\text{rev}}(1), \ldots, \mathbf{r}^{\text{rev}}(K)$. For each $i = 1, \ldots, K$, we first propose $M-1$ positions for the $i$-th bead in this segment, which we define, for $j = 1, \ldots, M-1$, as (cf. Eqn. 27)

$$\mathbf{r}_j^{\text{rev}}(i) = \begin{cases} \mathbf{q}(\mathbf{r}'_{N-K-2}, \mathbf{r}'_{N-K-1}, \mathbf{r}'_{N-K}, r_{i,j}, \theta_{i,j}, \varphi_{i,j}) & \text{if } i = 1 \\ \mathbf{q}(\mathbf{r}'_{N-K-1}, \mathbf{r}'_{N-K}, \mathbf{r}^{\text{rev}}(1), r_{i,j}, \theta_{i,j}, \varphi_{i,j}) & \text{if } i = 2 \\ \mathbf{q}(\mathbf{r}'_{N-K}, \mathbf{r}^{\text{rev}}(1), \mathbf{r}^{\text{rev}}(2), r_{i,j}, \theta_{i,j}, \varphi_{i,j}) & \text{if } i = 3 \\ \mathbf{q}(\mathbf{r}^{\text{rev}}(i-3), \mathbf{r}^{\text{rev}}(i-2), \mathbf{r}^{\text{rev}}(i-1), r_{i,j}, \theta_{i,j}, \varphi_{i,j}) & \text{otherwise,} \end{cases}$$

where, again, each bond length $r_{i,j}$, bond angle $\theta_{i,j}$, and dihedral angle $\varphi_{i,j}$ are sampled as described previously. Furthermore, we again reserve reversion to the original configuration as the final reverse move: for each $i = 1, \ldots, K$, we set (cf. Eqn. 22)

$$\mathbf{r}_M^{\text{rev}}(i) = \mathbf{r}_{N-K+i}.$$

We then calculate the Boltzmann weight, $w_j^{\text{rev}}(i)$, which is given by

$$w_j^{\text{rev}}(i) = \exp\left\{-\frac{V_j^{\text{rev}}(i)}{k_B T}\right\},$$

where $V_j^{\text{rev}}(i)$ follows a similar form to $V_j(i)$ in Eqn. 28:

$$V_j^{\text{rev}}(i) = \sum_{k=1}^{N_{\max}} U_{\text{LJ}}\left(\left\|\mathbf{r}_j^{\text{rev}}(i) - \mathbf{r}'_k\right\|\right) + \sum_{k=1}^{i-2} U_{\text{LJ}}\left(\left\|\mathbf{r}_j^{\text{rev}}(i) - \mathbf{r}_{N-K+k}\right\|\right),$$

where

$$N_{\max} = \begin{cases} N-K & \text{if } i > 1 \\ N-K-1 & \text{if } i = 1. \end{cases}$$

From here, we can calculate the $i$-th reverse Rosenbluth weight,

$$W_i^{\text{rev}} = \sum_{j=1}^{M} w_j^{\text{rev}}(i),$$

which we can combine to arrive at the total reverse Rosenbluth factor,

$$W_{\text{rev}} = \prod_{i=1}^{K} W_i^{\text{rev}},$$

which we can use to finally calculate the acceptance probability via Eqn. 14.

##### 2.2.3 Initialization

To initialize each run of the CBMC procedure, we generated a polymer configuration by iteratively sampling bond lengths, bond angles, and dihedrals and generating bead positions as described in §2.1.2, while also enforcing a minimum-distance requirement between non-bonded pairs of beads. In particular, starting from a position for the first bead, $\mathbf{r}_1 = \mathbf{0}$, we generated a candidate position for the $i$-th bead, $\mathbf{r}_i$, along the chain by sampling a bond length, $r \in (0, R_0)$; a bond angle, $\theta \in [0, \pi]$; and a dihedral angle, $\varphi \in [-\pi, \pi)$, as described in §2.1.1, and setting[2]

$$\mathbf{r}_i = \mathbf{q}(\mathbf{r}_{i-3}, \mathbf{r}_{i-2}, \mathbf{r}_{i-1}, r, \theta, \varphi).$$

We then accepted this candidate position if $\left\|\mathbf{r}_i - \mathbf{r}_j\right\| < \epsilon$ for all $j = 1, \ldots, i-2$, where the minimum distance $\epsilon$ was set to the length-scale of the non-bonded interaction potential (Eqn. 1), $\epsilon = 2^{1/6}\sigma$. To enable escape from neighborhoods in configurational space where unacceptably close candidate positions were repeatedly encountered, we also allowed for backtracking to and resampling the previous bead position whenever a maximum number of candidate positions was exceeded.

##### 2.2.4 CBMC for a polymer solution

To adapt our CBMC procedure to a solution of multiple chains, we defined a system as $N_{\text{chains}}$ chains of the same length $N$, situated within a cubic domain $\Omega = [x_{\min}, x_{\max}] \times \left[y_{\min}, y_{\max}\right] \times [z_{\min}, z_{\max}]$ with periodic boundary conditions. As such, all interaction energies were computed with respect to norms and angles derived from periodic distance vectors,

$$\mathbf{d}_\Omega(\mathbf{p}, \mathbf{q}) = \left(u_x - \Delta x \cdot \lfloor u_x/\Delta x \rceil, u_y - \Delta y \cdot \lfloor u_y/\Delta y \rceil, u_z - \Delta z \cdot \lfloor u_z/\Delta z \rceil\right),$$

where $\mathbf{u} = \mathbf{p} - \mathbf{q}$, $\Delta x = x_{\max} - x_{\min}$, $\Delta y = y_{\max} - y_{\min}$, $\Delta z = z_{\max} - z_{\min}$, and $\lfloor \cdot \rceil$ denotes rounding to the nearest integer.

During each sampling iteration, we uniformly sampled a chain to perturb, and perturbed the chain using the same move set as in the single-chain setting—single-bead reptation, multi-bead reptation, and terminal segment moves—while also incorporating *inter-chain* non-bonded interaction energies into the Boltzmann weight calculations. For instance, suppose we are applying a $K$-bead reptation move on the $p$-th chain in the solution (§2.2.1). Then the Boltzmann weight of the $j$-th candidate position for the $i$-th bead in the proposed segment is calculated as (cf. Eqn. 16)

$$w_j(i) = \exp\left\{-\frac{V_j(i)}{k_B T}\right\},$$

where $V_j(i)$ is the non-bonded interaction energy between the candidate position, $\mathbf{r}_j^{\text{new}}(i)$, and:

[2]For $i = 2$, we set $\mathbf{r}_2$ by uniformly sampling a vector $\hat{\mathbf{u}}$ on the unit sphere, and setting $\mathbf{r}_2 = \mathbf{r}_1 + r\hat{\mathbf{u}}$. For $i = 3$, we set $\mathbf{r}_3$ by generating an orthonormal basis for $\mathbb{R}^3$ containing the vector $\hat{\mathbf{u}} = (\mathbf{r}_1 - \mathbf{r}_2) / \|\mathbf{r}_1 - \mathbf{r}_2\|$, then transforming the vector $(\cos\theta, \sin\theta, 0)$ according to the change-of-basis which sends $\hat{\mathbf{x}} \mapsto \hat{\mathbf{u}}$, $\hat{\mathbf{y}} \mapsto \hat{\mathbf{v}}$, and $\hat{\mathbf{z}} \mapsto \hat{\mathbf{w}}$. We then used the resulting vector, $\hat{\mathbf{q}}$, to define $\mathbf{r}_3 = \mathbf{r}_2 + r\hat{\mathbf{q}}$.

1. every bead in the original configuration for the $p$-th chain, $X_p^{(n)}$, that would survive the reptation move;

2. every bead that has been generated thus far in the segment except for the $(i-1)$-th bead, i.e., $\mathbf{r}^{\text{new}}(1), \dots, \mathbf{r}^{\text{new}}(i-2)$; and

3. every bead in every other chain in the configuration.

In other words, we have (cf. Eqn. 17)

$$V_j(i) = \sum_{k=K+1}^{N_{\max}} U_{\text{LJ}}\left(\left\|\mathbf{r}_j^{\text{new}}(i) - \mathbf{r}_k\right\|\right) + \sum_{k=1}^{i-2} U_{\text{LJ}}\left(\left\|\mathbf{r}_j^{\text{new}}(i) - \mathbf{r}^{\text{new}}(k)\right\|\right)$$
$$+ \sum_{\substack{1 \le q \le N_{\text{chains}} \\ q \neq p}} \sum_{k=1}^{N} U_{\text{LJ}}\left(\left\|\mathbf{r}_j^{\text{new}}(i) - \mathbf{r}_k(X_q^{(n)})\right\|\right),$$

where $\mathbf{r}_k(X_q^{(n)})$ is the $k$-th bead position in the $q$-th chain in $X^{(n)}$, and, as before,

$$N_{\max} = \begin{cases} N & \text{if } i > 1 \\ N-1 & \text{if } i = 1. \end{cases}$$

The Boltzmann weight calculations for the other moves were adapted similarly.

The final difference between the single-chain and multi-chain sampling procedures lies in the initialization, as described in the Methods. First, we generated a starting configuration by following a variant of the procedure described in §2.2.3 but iterated $N_{\text{chains}}$ times, to generate bead positions $\left\{\left(\mathbf{r}_1^{(p)}, \dots, \mathbf{r}_N^{(p)}\right) : p = 1, \dots, N_{\text{chains}}\right\}$. We also introduced the following adjustments:

1. We checked that each candidate bead position, $\mathbf{r}_i^{(p)}$, is further than $\epsilon_{\text{intra}}$ from all beads along the $p$-th chain generated thus far other than $\mathbf{r}_{i-1}^{(p)}$,

$$\left\|\mathbf{r}_i^{(p)} - \mathbf{r}_j^{(p)}\right\| < \epsilon_{\text{intra}} \qquad \text{for all } j = 1, \dots, i-2,$$

   *and* that it is further than $\epsilon_{\text{inter}}$ from all beads along every other chain generated thus far,

$$\left\|\mathbf{r}_i^{(p)} - \mathbf{r}_j^{(q)}\right\| < \epsilon_{\text{inter}} \qquad \text{for all } j = 1, \dots, N \text{ and } q = 1, \dots, p-1.$$

   We set $\epsilon_{\text{intra}} = 2^{1/6}\sigma$ as in the single-chain setting, but we set $\epsilon_{\text{inter}}$ to a range of values less than $2^{1/6}\sigma$ (0.5–1.5 nm; compare with $2^{1/6}\sigma \approx 2.0937$ nm), depending on the concentration.

2. In addition to allowing for backtracking along each chain, we also allowed for re-seeding a chain from scratch if a maximum number of backtracks was exceeded along that chain, or if backtracking led to encroaching back into the first three beads of the chain.

The configuration obtained via this procedure was then equilibrated using LAMMPS (Thompson et al. 2022) to gradually increase inter-chain distances, as described in the Methods. The final configuration obtained from this equilibration was then used as the initial configuration for the CBMC procedure.

**References.**


Allen, Michael P. and Dominic J. Tildesley (2017). *Computer Simulation of Liquids*. 2nd. Oxford, United Kingdom: Oxford University Press.

Best, D.J. and N.I. Fisher (1979). "Efficient simulation of the von Mises distribution". In: *J. R. Stat. Soc. C* 28, pp. 152–157. DOI: 10.2307/2346732.

Everaers, Ralf, Hossein Ali Karimi-Varzaneh, Frank Fleck, Nils Hojdis, and Carsten Svaneborg (2020). "Kremer–Grest models for commodity polymer melts: linking theory, experiment, and simulation at the Kuhn scale". In: *Macromolecules* 53, pp. 1901–1916. DOI: 10.1021/acs.macromol.9b02428.

Kremer, Kurt and Gary S. Grest (1990). "Dynamics of entangled linear polymer melts: A molecular-dynamics simulation". In: *J. Chem. Phys.* 92, pp. 5057–5086. DOI: 10.1063/1.458541.

Milano, Giuseppe, Sylvain Goudeau, and Florian Müller-Plathe (2005). "Multicentered Gaussian-based potentials for coarse-grained polymer simulations: Linking atomistic and mesoscopic scales". In: *J. Polym. Sci. B* 43, pp. 871–885. DOI: 10.1002/polb.20380.

Siepmann, Jörn Ilja and Daan Frenkel (1992). "Configurational bias Monte Carlo: a new sampling scheme for flexible chains". In: *Mol. Phys.* 75, pp. 59–70. DOI: 10.1080/00268979200100061.

Svaneborg, Carsten and Ralf Everaers (2020). "Characteristic time and length scales in melts of Kremer–Grest bead-spring polymers with wormlike bending stiffness". In: *Macromolecules* 53, pp. 1917–1941. DOI: 10.1021/acs.macromol.9b02437.

Thompson, Aidan P. et al. (2022). "LAMMPS – a flexible simulation tool for particle-based materials modeling at the atomic, meso, and continuum scales". In: *Comput. Phys. Commun.* 271, p. 108171. DOI: 10.1016/j.cpc.2021.108171.

Weeks, John D., David Chandler, and Hans C. Andersen (1971). "Role of repulsive forces in determining the equilibrium structure of simple liquids". In: *J. Chem. Phys.* 54, pp. 5237–5247. DOI: 10.1063/1.1674820.
